# Revisiting the Evolution of Lactase Persistence: Insights from South Asian Genomes

**DOI:** 10.1101/2025.11.05.686799

**Authors:** Elise Kerdoncuff, Meaghan Marohn, Nathan Cramer, Sharmistha Dey, Sharon Kardia, Kumarasamy Thangaraj, Laure Ségurel, Jinkook Lee, Aparajit Ballav Dey, Priya Moorjani

## Abstract

Lactase persistence (LP), the ability to digest lactose from milk into adulthood, is a classic example of natural selection in humans. Multiple mutations upstream of the *LCT* gene are associated with LP and have been previously shown to be under selection in Europeans and Africans. South Asia is the world’s largest producer of dairy, and milk and dairy products are widely consumed throughout the subcontinent. However, the origin, evolutionary history and selective pressures associated with LP in South Asia remain elusive. We assembled genome-wide data from ∼8,000 present-day and ancient genomes from India, Pakistan, and Bangladesh, spanning diverse timescales (∼3300 BCE–1650 CE), geographic regions, and ethnolinguistic and subsistence groups. We find that the Eurasian LP-associated variant,-13.910:C>T, is widespread across South Asia, exhibiting clinal variation along north-south and east-west gradients. Ancient DNA analysis reveals that this variant first appeared in South Asia during the historical and medieval periods through Steppe pastoralist-related gene flow. Interestingly, unlike in other worldwide populations, the LP prevalence is almost entirely explained by Steppe ancestry—not selection––in most contemporary South Asians. A notable exception is the only two pastoralist groups, Toda in South India and Gujjar in Pakistan, that have unexpectedly high frequencies of-13.910*T, comparable to estimates in Northern Europeans. By performing local ancestry inference, we find significant enrichment for Steppe pastoralist ancestry around the *LCT* locus in these two geographically-distant pastoralist groups, indicative of strong selection. Together, these findings highlight the complex role of ancestry and natural selection in shaping the prevalence of lactase persistence on the subcontinent.

## Introduction

Lactase persistence (LP)—the ability to digest lactose into adulthood— is a classic example of positive selection in humans. This trait is associated with regulatory mutations upstream of the *LCT* gene, which encodes the lactase enzyme^1,2^. Among these,-13.910:C>T and-22.018:G>A located in introns 9 and 13 of the *MCM6* gene are significantly associated to LP in Eurasians, although functional studies have indicated that the key variant regulating maintenance of *LCT* expression in adults is-13.910:C>T^3–5^. Additional independent variants including-13.915:T>G,-14.010:G>C,-13.907:C>G and-14.009:T>G have been identified in African and Middle Eastern pastoralist groups^6–9^. Genetic analysis has demonstrated that LP-associated variants,-13.910:C>T and-14.010:G>C, arose independently and spread to high frequency in European and African populations respectively, exhibiting strong signatures of natural selection^6–9^. Together, these results suggest convergent evolution for LP in humans.

The prevailing evolutionary model posits that, following the domestication of milk-producing animals, adults capable of digesting lactose gained a fitness advantage—particularly in environments where milk provided essential calories, hydration, or nutrients such as calcium^1,10,11^. Consequently, LP alleles experienced strong positive selection, representing a classic case of gene–culture coevolution^10,11^. However, this model does not fully explain the global diversity of LP or its selective advantage^12^. Analysis of ancient dental calculus and pottery indicates that milk use in Europe preceded the rise of LP alleles by thousands of years, suggesting that cultural practices such as fermentation or reduced lactose intake may have mitigated lactose intolerance. Nonetheless, during times of nutritional stress (e.g., famine) or increased pathogen exposure, the inability to digest lactose may have conferred a survival disadvantage, driving LP selection^13^. Furthermore, recent findings suggest that the *LCT* locus harbors genes related to immune function that might be under selection, independent of lactase activity^14^.

South Asia presents a particularly intriguing context for studying the evolution of LP. As the world’s largest producer of dairy, the region supports extensive use of milk and dairy products across a wide range of cultural and geographic settings^15,16^. Cattle domestication, especially *Bos indicus* (zebu) dates to ∼7,000–6,000 years ago in Mehrgarh (present-day Pakistan)^17^. Yet, despite the centrality of dairy in contemporary diets, the evolutionary history and selective dynamics of LP in South Asia remain poorly understood. Recent studies have shown that the Eurasian variant-13.910:C>T accounts for nearly all the genetic variation in LP in India, and correlates strongly with phenotypic differences in lactose digestion across regions^18,19^. However, many open questions remain: What do patterns of LP look like in South Asian populations outside India, e.g., Pakistan and Bangladesh? Did LP arise in South Asia at the onset of agriculture during the Neolithic or did it spread through Steppe pastoralists-related gene flow as seen in Europe^12^? Is there evidence for selection in South Asia and what are the underlying cultural and ecological drivers?

In this study, we analyze genome-wide data from approximately 8,000 present-day and ancient individuals from India, Pakistan, and Bangladesh, spanning a wide range of temporal, geographic, urban, rural and tribal groups including two pastoralist groups (Toda and Gujjar). The Toda, traditional buffalo herders from the Nilgiri Hills of South India, consume milk in diverse forms, including fresh milk, butter, buttermilk, yogurt, and cheese^20^. Similarly, the Gujjars, semi-nomadic cattle and buffalo herders from North India and Pakistan, base both their subsistence and economy on milk production and dairy products^21,22^. We map the frequency and distribution of the-13.910:C>T variant across the subcontinent and explore its correlation with ancestry and subsistence strategies. By integrating ancient DNA, haplotype analysis, and demographic modeling, we aim to disentangle the roles of gene flow, drift, and selection in shaping the landscape of LP in South Asia.

## Results

### Data Description and Population Structure in South Asia

We assembled a genome-wide dataset of 8,091 present-day and ancient genomes from India, Pakistan, and Bangladesh, including diverse timescales, geographic regions, ethnolinguistic, and subsistence groups (Figure 1A). For present-day individuals, we used 7,962 whole genome sequences from unrelated individuals from three published datasets: the Longitudinal Aging Study of India-Harmonized Diagnosis of Dementia (LASI-DAD)^23^, Genome Asia wave 1 (GAsP1)^24^ and Genome Asia wave 2 (GAsP2)^25^ (Supplementary Note 1; Supplementary Table 1.1; Methods). We retained 45 endogamous groups with at least 5 individuals (*n* = 5–34 individuals per group), including two pastoralists groups (*n* = 18–20 per group). We analyzed 129 ancient individuals from the Swat Valley in present-day Pakistan, dated to ∼3300 BCE–1650 CE, from the Allen Ancient DNA Resource (AADR) captured on the 1240K SNP array^26^. For most analyses, we grouped individuals by sampling age, endogamous community, geographic region or country of origin––India (*n* = 5,535), Pakistan (*n* = 1,905) or Bangladesh (*n* = 522). To account for the dense sampling in India, we further grouped individuals from India by region of sampling–– North (*n* = 571), Central (*n* = 429), East (*n* = 1,791), North-East (*n* = 87), West (*n* = 416) and South (*n* = 2,241).

**Figure 1.**
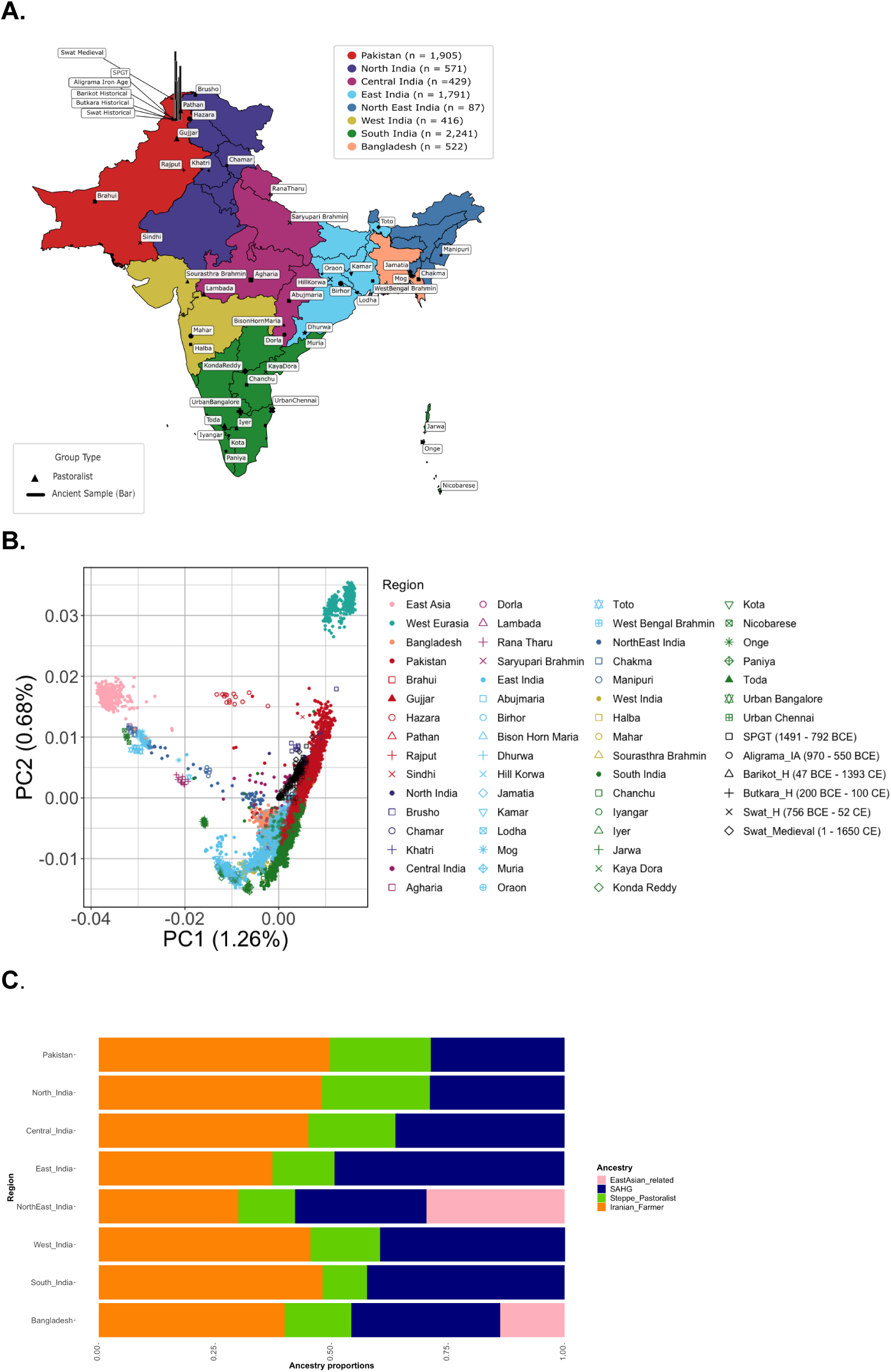
Dataset presentation and evolutionary history of South Asia A. Map of South Asia Data Sampling. Each geographic region of India, all of Pakistan, and all of Bangladesh are colored to reflect whole genome sequences that are sampled based on population density from LASI-DAD and medical cohorts from GAsP2 (the legend for each region in the upper left). Whole genome sequences of endogamous groups from GAsP1 are labeled and plotted in different symbols to match the PCA in B and are scaled according to sample size (n = 5–34). Pastoralist groups are shown in solid triangles, while non-pastoralists are shown in different symbols per region as in (B). Ancient DNA samples from the Swat Valley in Pakistan are represented by vertical bars that vary in length according to the age of the samples (∼1491 BCE - 1650 CE). **B. Principal Components Analysis of Present-Day and Ancient South Asians** Principal Components Analysis was done on unrelated present-day South Asian individuals (India, Pakistan, and Bangladesh), to demonstrate how South Asians relate to other global groups, East Asians and West Eurasians. All individuals are plotted in color according to their geographic region (teal for West Eurasians, pink for East Asians, and South Asians following the same color scheme in (A)). Endogamous groups are plotted by the color of their geographic region and the same symbol as the map in (A), with pastoralists represented as a solid triangle. Ancient South Asians are projected in black in different symbols for each time period. **C. Ancestry Modeling of Present-Day South Asians** For each region in South Asia (Pakistan, regions of India, and Bangladesh), the average proportion Iranian farmer-related (orange), Steppe pastoralist-related (green), SAHG-related (navy), and East Asian-related (pink) ancestries are shown in a bar summing to 1 (100%). Along the y-axis, the groups are arranged geographically from north to south and west to east.

To investigate the patterns of genetic variation in South Asia and their relationship to other worldwide populations, we combined our dataset with the 1000 Genomes Project (1000G)^27^ and the Human Genome Diversity Panel (HGDP)^28^ and performed Principal Component Analysis (PCA) and ADMIXTURE (Methods). As seen in previous studies, we find most South Asians fall on a cline of relatedness to West Eurasians, referred to as the ‘Indian cline’^29,30^. Individuals from South India cluster towards one end of the cline and individuals from North India and Pakistan fall on the other end closer to West Eurasians (Figure 1B). Individuals from Bangladesh, East India, and North-East India are shifted toward East Asians. Individuals within endogamous communities tend to be genetically homogeneous and generally cluster by geographic region (Figure 1B, Figure S1.2 A)^29–31^. Some individuals from North India and Pakistan exhibit recent relatedness to sub-Saharan African individuals, potentially due to gene flow from the Middle East as observed in earlier surveys (Figure S1.1)^29,32^. We observe qualitatively similar patterns of population relationships using ADMIXTURE analysis (Supplementary Note 1, Figure S1.4).

Ancient DNA analysis has recently shown that the individuals on the Indian cline have ancestry from three ancestral groups related to ancient Neolithic Iranian farmers, ancient Eurasian Steppe pastoralists, and South Asian hunter-gatherers (*SAHG*) that are related to Andamanese islanders^23,31^. To model the ancestry in South Asia, we used *qpAdm*^33,34^ that compares allele frequency correlations between a population of interest and a set of reference and outgroup populations (Methods; Supplementary Note 1). We used *Indus_Periphery_West* (the highest coverage individual among the Iranian farmer-related ancient DNA samples) as the proxy for Neolithic Iranian farmers; Central Steppe individuals from the Middle to late Bronze age (*Central_Steppe_MLBA* dated ∼2000-900 BCE) for Steppe pastoralist-related ancestry; and present-day Andamanese islanders (*Onge* from GAsP1) to represent *SAHG*-related ancestry. The three-way model provides a good fit for 4,946 individuals on the Indian Cline (>83%), and an additional 580 individuals that do not fall on the cline (we define ‘good fit’ as *p*-value > 0.01 and ancestry coefficients > 0 for *qpAdm* analysis; Supplementary Table 1.2, Supplementary Note 1). For individuals on the Indian cline, we infer Iranian farmer-related ancestry ranges from 26.3-69.7%, Steppe pastoralist-related ancestry ranges from 0.3-46.4%, and *SAHG*-related ancestry ranges from 3.7-64.1% (Figure 1C; Supplementary Table 1.2. Because *Indus_Periphery_West* harbors ∼15–20% *SAHG*-related ancestry, our estimates reflect correspondingly higher SAHG ancestry, aligning more closely with earlier surveys^23,31^). Steppe pastoralist-related ancestry is higher in North India and Pakistan (∼22%) compared to South India (∼10%) (*p*-value < 0). Conversely, *SAHG*-related shows the reciprocal pattern (*p*-value < 1e-12). For individuals from Bangladesh (*n* = 495) and North-East India (*n* = 71) that fall outside the Indian cline, we also applied a four-way model with the addition of East Asian-related ancestry (Han Chinese in Beijing (CHB) from 1000G) and found it varies between 0.1–80.9% across South Asia (Supplementary Table 1.2; Figure S1.6 A).

For ancient DNA samples from Swat Valley, the three-way model provides a good fit to the majority (>87%) of the individuals. The *SAHG*-related ancestry ranges from 1.7–24.9%, Iranian farmer-related ancestry ranges from 36.6–86.1% and Steppe pastoralist-related ancestry ranges from 0.8–50.7% (Figure S1.6 B; Supplementary Table 1.2). The ancestry composition of ancient individuals is consistent with their geographic location and inferred estimates for present-day individuals from Pakistan and North India (Figure 1). Together, our dataset provides the most comprehensive view to date of population diversity and ancestry variation in South Asia, capturing both temporal shifts and regional heterogeneity in genetic composition.

### Distribution of LP-associated variants varies by Geography, Timescale, and Subsistence Practices

To understand the patterns of LP in South Asians, we first examined thirteen LP-associated variants–-13.910:C>T,-22.018:G>A,-13.915:T>G,-13.907:C>G,-14.009:T>G,-14.010:G>C,-13.779:G>C,-13.801:C>T,-13.879:G>A,-13.915:T>C,-14.011:C>T,-14.012:A>G,-14.026:T>C––identified in previous surveys^3,6–9,18^. We also examined the frequency of-13.910:C>T in 85 ancient South Asians and 23 Central Asians from the AADR dataset^26^, using two approaches (i) high-quality imputation and (ii) read count data requiring at least three reads at the locus of interest (we note other LP-associated variants are not included on the 1240K array). Both approaches give consistent results, and we report frequencies based on imputed genotypes for ancient individuals below (Supplementary Table 2.1).

Among the known LP-associated variants, we found that-13.910:C>T variant, the key variant regulating maintenance of *LCT* expression in Europeans, is the most widespread across South Asia, with frequencies on average reaching ∼10-15% compared to < 1% for other variants (Supplementary Note 2). Because this allele confers a dominant phenotype, we estimated the frequency of lactase persistence trait (hereafter “LP”, shown in brackets) by weighting the heterozygous and homozygous genotypes equally (see Methods). The 13.910*T allele exhibits a clinal distribution across the subcontinent, with highest frequencies above 25% in North India and Pakistan (LP = 44.5%), decreasing to less than 10% in East India (LP = 13.77%) and less than 5% in South India (LP = 6.56%) (Figure 2A; Supplementary Table 2.1). Similar trends are observed in earlier surveys with fewer individuals^18^ (Supplementary Table 2.1) and in 1000G South Asians (Figure S2.6, Supplementary Table 2.1). Across endogamous groups, nearly half––predominantly from the South and East India––completely lack-13.910*T (Figure 2B; Supplementary Table 2.1). Strikingly, the only two pastoralist groups from our dataset–– Toda (South India) and Gujjar (Pakistan)–– exhibit exceptionally high frequencies reaching 61% (LP = 89%) and 67.5% (LP = 90%), respectively (Figure 2B; Supplementary Table 2.1), similar to estimates seen in Northern Europeans^3^.

**Figure 2.**
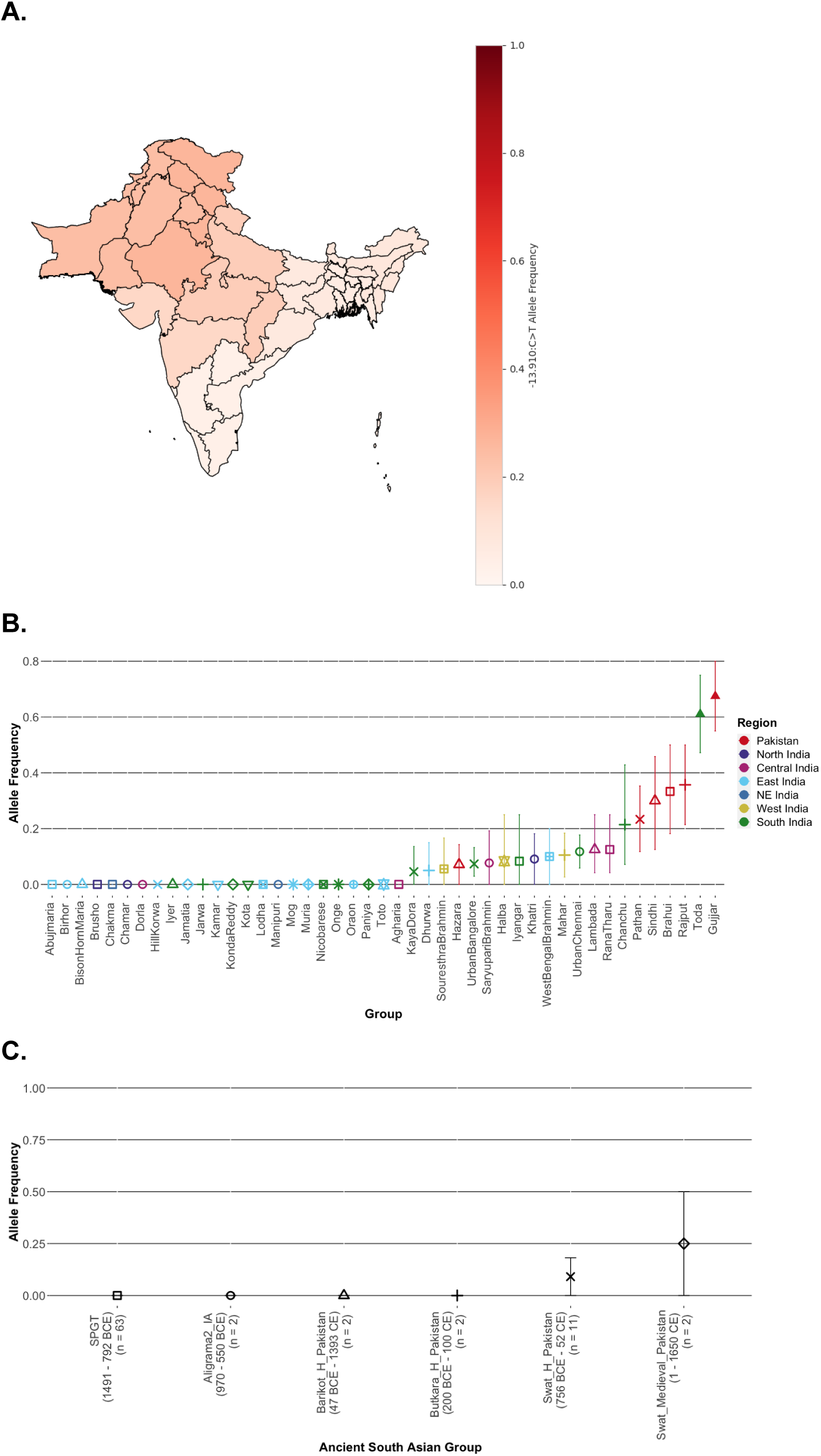
Allele frequency of-13.910:C>T across space and time in South Asia **A. Allele Frequencies across South Asia** The allele frequency (red scale) was calculated across Pakistan, India, and Bangladesh. The allele frequency is highest in Pakistan and North India, and decreases in South India, East India, and Bangladesh. **B. Allele Frequency in Endogamous Groups** The allele frequency was calculated for each endogamous group from GAsP1 across India and Pakistan. The allele frequency of each endogamous group increased with geography, that is, groups in South India have low allele frequencies or are zero, and groups in North India and Pakistan have higher allele frequencies. The pastoralist groups Toda (South India) and Gujjar (Pakistan) are notable outliers. Groups are plotted by color and symbol as in Figure 1A and Figure 1B, representing geographic regions. **C. Allele Frequency in Ancient South Asians** The allele frequency was calculated for ancient South Asians from Pakistan (∼1491 BCE - 1650 CE) for individuals that had at least 3 reads at-13.910:C>T. Each ancient group is shown in the same symbol as plotted in Figure 1B. We note the allele is absent in samples from the Bronze and Iron Ages, but increases in frequency in the Historic and Medieval periods.

The-22.018:G>A variant, also discovered in Europeans, displays a geographic distribution that closely mirrors that of-13.910:C>T. This concordance reflects strong, but incomplete, linkage disequilibrium (LD) between these two variants in South Asia—unlike the complete LD observed in Europeans (*r^2^* = 0.79–1; Supplementary Table 2.1; Figure S2.1, Figure S2.7, Supplementary Note 2)^5^. In contrast, LP-associated variants discovered in individuals from the Middle East and Africa (-13.915:T>G,-13.907:C>G,-14.009:T>G,-14.010:G>C)^6–9^ were absent in our dataset (Supplementary Table 2.1; Figure S2.2-4, Supplementary Note 2). Moreover, a recent study identified seven additional candidate variants in the enhancer of *LCT* which may contribute to regional differences in India—-13.779:G>C,-13.801:C>T,-13.879:G>A,-13.915:T>C,-14.011:C>T,-14.012:A>G,-14.026:T>C^18^. We find these variants were either absent or extremely rare (< 5% on average, Supplementary Table 2.1; Figure S2.4, Supplementary Note 2) as seen earlier^18^.

In ancient DNA samples from South Asia, we find-13.910*T is absent in Bronze Age and Iron Age individuals but the allele frequency rises to an average of 9.1% in the historic period (LP = 18.18%) and 25% in the medieval period (LP = 50%) (Figure 2C; Supplementary Table 2.1). Among the three ancestral proxies used in *qpAdm* analysis,-13.910*T is absent in *SAHG*-related individuals (present-day Onge) and Neolithic Iranian farmers (*Sarazm_EN* and *Indus_Periphery_Pool*), but reaches 10.9% in Bronze Age Steppe pastoralist-related individuals (*Central_Steppe_MLBA,* LP = 17.4%) (Supplementary Table 2.1). These results show that among the known LP-associated variants, only –13.910:C>T (and –22.018:G>A) reached appreciable frequencies in South Asia, with strong geographic and temporal variation. Based on these observations, we focused on-13.910:C>T for the majority of downstream analyses as it has also been previously shown to be the driver of *LCT* expression in adults^19^.

### Steppe Pastoralist Origin of Lactase Persistence in South Asia

To trace the origin of lactase persistence in South Asia, we analyzed the haplotype structure around-13.910:C>T in present-day South Asians and compared it to haplotypes in ancestral reference groups––*SAHG*, Iranian farmer, and Steppe pastoralist (see Methods). We focused on the core region (chr2:135,787,850–135,876,443) that spans 88 Kb and includes the *LCT* and *MCM6* genes, and stratified individuals based on the presence or absence of the derived allele at-13.910:C>T.

We find the present-day South Asian haplotypes harboring-13.910*T (South Asian*T, *n* = 2,065) have markedly lower diversity than the-13.910*C haplotypes (South Asian*C, *n* = 13,859), including a higher proportion of fixed variants (68.3% vs. 0.7%; *p*-value < 2.2e-16), depletion of SNPs at intermediate (25–75%) allele frequencies (0% vs. 21.8%; *p*-value < 2.2e-16) and lower entropy (0.005 vs. 0.2, *p*-value < 2.2e-16) (Figure S3.1-2). Comparing the present-day South Asians to the ancestral reference groups, we find South Asian*T haplotypes are nearly identical to the Steppe pastoralist*T haplotypes, with an average of only 0.19 differences over 177 SNPs in the core region (we note that-13.910*T allele is absent in Iranian farmers and *SAHG*-related populations). The average differences between South Asian*T haplotypes were significantly higher when compared to other groups: 41.79 with Iranian farmer*C, 38.58 with Steppe pastoralist*C, and 19.12 with SAHG*C haplotypes (Figure S3.3). To assess statistical significance of the pairwise differences among haplotypes, we randomly permuted the ancestry labels of the reference haplotypes (10,000 iterations) and recalculated the mean of the empirical null distribution. The expected mean in a set of random five haplotypes is 29.36, compared to the observed value of 0.19 (p-value < 1e-4). In contrast, the South Asian*C haplotypes showed no clear pattern of affinity to any particular ancestral population. The average differences were broadly similar across all three groups: 28.85 (Iranian farmer*C), 35.44 (SAHG*C), 30.91 (Steppe pastoralist*C), and 40.49 (Steppe pastoralist*T) (Figure S3.4).

A limitation of our analysis is the use of low-coverage ancient genomes imputed with contemporary reference data(1000 Genomes; Glimpse v1.0)^35^, which may bias results in complex regions like *LCT*^36^. To address this, we compared present-day South Asians and Europeans (GBR, 1000 Genomes, 394 SNPs in the core region). In Europeans,-13.910*T originated through Steppe pastoralist-related gene flow and was subsequently under positive selection^37^. South Asian*T haplotypes are nearly identical to GBR*T (average 0.33 differences) but highly divergent from GBR*C (60.54), while South Asian*C haplotypes show similar divergence from both GBR*T (55.30) and GBR*C (53.01) (Figure S3.4).

This pattern is also reflected in the extended haplotype—a 870 kb region in strong linkage disequilibrium (r² > 0.8) with-13.910:C>T variant (Figure S2.7). The extended haplotype exhibits a similar profile to the core haplotype: low diversity and high similarity to the Steppe pastoralist*T haplotypes (Figure S3.1,2,8). Moreover, the length of the haplotype is consistent with theoretical expectations for an admixture tract length for Steppe pastoralist related gene flow that occurred around 3,500 years ago as inferred using ancient DNA^31^ (Figure S4.1).

Together, these results suggest-13.910*T originated through pastoralist-related gene flow in South Asia, similar to Europeans.

### Ancestry Explains Patterns of-13.910*T Allele Frequency in most South Asians

To understand the role of ancestry in shaping lactase persistence in South Asia, we examined the relationship between ancestry and-13.910*T frequency (*n* = 7,391, excluding individuals with >10% sub-Saharan African-related ancestry). We used PCA loadings as a proxy for genome-wide ancestry (Figure 1B) and binned individuals (excluding endogamous groups that were projected later) based on PC1 and PC2 values, ensuring equal numbers of individuals in each bin (see Methods). For each bin, we computed the average allele frequency of-13.910*T and median PC loadings and tested their linear association (Methods, Supplementary Note 3). We find that there is a strong association between-13.910*T allele frequency and PC1 (r^2^ = 0.91, *p*-value < 1e-06) and PC2 (r^2^ = 0.87, *p*-value < 1e-06) (Figure S5.1 B). Projecting endogamous groups onto the regression model showed that most groups lie on or overlap the regression line (after accounting for uncertainty) (Figure S5.1 B). Notably, 17 groups that lack-13.910*T fall within the first PC bin (X = 0). The two pastoralist groups, Toda and Gujjar, are significant outliers (Z-score > 3), deviating markedly from the expected ancestry–allele frequency relationship (Figure S5.1B, Supplementary Note 5.1). We find qualitatively similar results when considering LP frequency (accounting for dominance) instead of allele frequency (Figure S5.2; Supplementary Note 5.1)

To more directly investigate the ancestry-allele frequency relationship, we used the genome-wide ancestry proportions of *SAHG*, Iranian farmers and Steppe pastoralist populations for individuals on the Indian cline (*n* = 4,946) (Supplementary Table 1.2; Supplementary Note 1). We find there is strong positive correlation between Steppe pastoralist-related ancestry and-13.910*T allele frequency (*r^2^* = 0.97, *p*-value < 1e-06) (Figure 3). In contrast, *SAHG*-related ancestry is negatively correlated with allele frequency (r^2^ =-0.96, p-value < 1e-06) (Figure 3). Iranian farmer-related ancestry also shows a significant positive association (*r^2^* = 0.69, p-value = 7.0e-04) (Figure 3), though the combined model with Steppe pastoralist-and Iranian farmer ancestries explains less variation than Steppe pastoralist-related ancestry alone (*r^2^* = 0.61, p-value < 0.05 based on permutation test; Figure S5.5, Supplementary Note 5.2). Again, the two South Asian pastoralist groups––Toda and Gujjar––are significant outliers (Z-score > 3), deviating markedly from the regression line (Figure 3; Figure S5.3, Supplementary Note 5.2). We find qualitatively similar results when considering LP frequency instead of allele frequency (Figure S5.4, Figure S5.6, Supplementary Note 5.2). Overall, these analyses suggest that Steppe pastoralist ancestry explains nearly all variation in-13.910*T allele frequency in most contemporary South Asians, with the notable exception of pastoralist groups––Toda and Gujjar.

**Figure 3.**
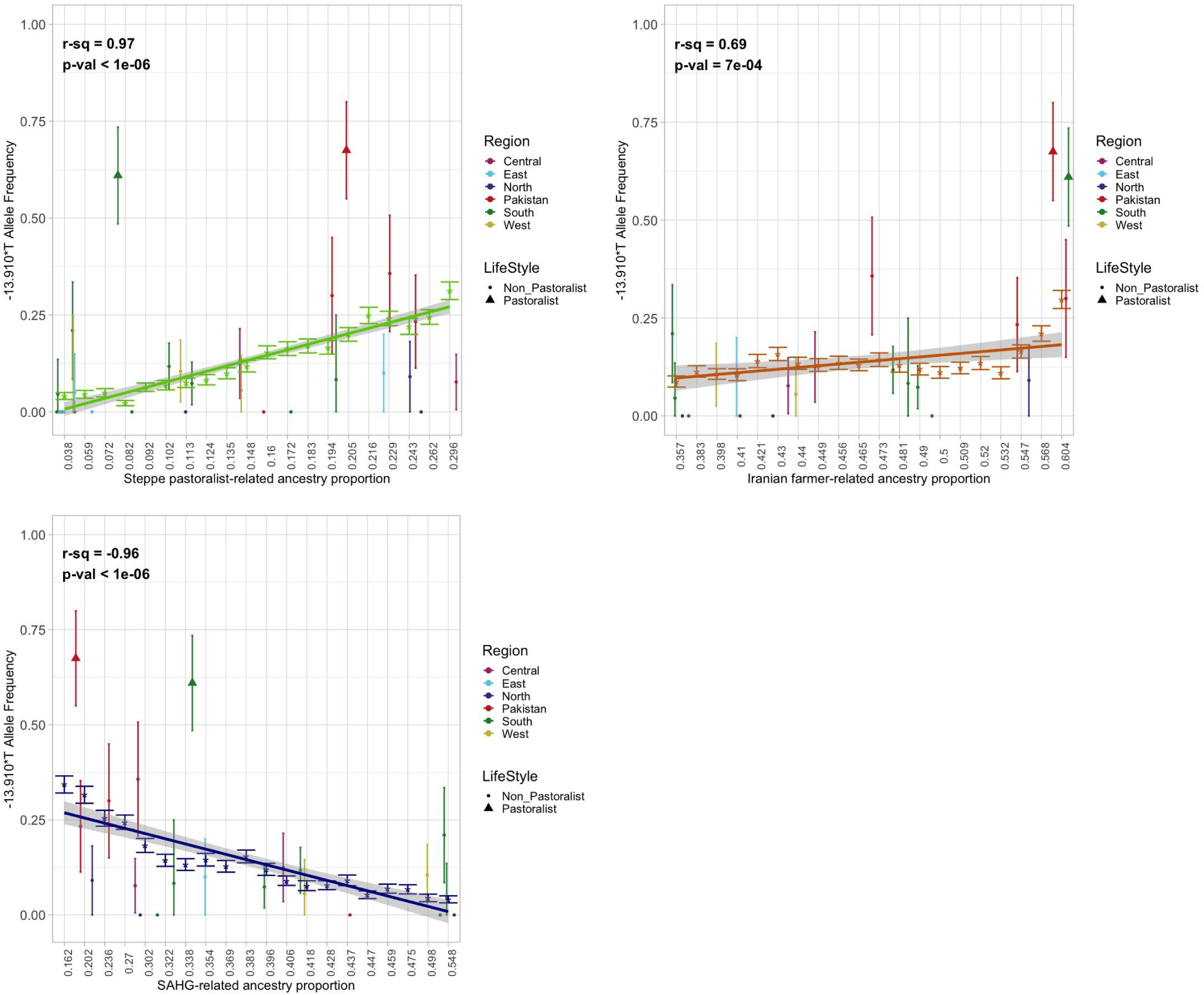
The correlation between ancestry proportions and allele frequency of-13.910:C>T in South Asians. All South Asian individuals were grouped into bins according to their Steppe pastoralist-related, Iranian farmer-related, and SAHG-related ancestry proportions estimated from qpAdm so an even number of individuals were in each bin. The median value of each bin is reported on the x-axis. Each bin’s average allele frequency was calculated and plotted along the y-axis. The regression line shows the relationship between each bin’s median ancestry proportion and corresponding mean allele frequency. The relationship between allele frequency and Steppe pastoralist-related ancestry is shown in light green, Iranian farmer-related ancestry in dark orange, and SAHG-related ancestry in navy blue. Endogamous groups are projected on each regression line according to their mean ancestry proportion and mean allele frequency. Groups are colored according to geographic region as in Figure 1 A and B, and pastoralist groups Toda and Gujjar are shown as solid triangles, with the rest of the non-pastoralist groups shown as dots.

### Can Genetic Drift Alone Explain the High LP Frequency in South Asian Pastoralists?

We performed simulations reflecting the key features of the demographic histories of Toda and Gujjar to evaluate whether drift alone could explain the elevated-13.910*T frequencies. Both Toda and Gujjar harbor Steppe pastoralist-related ancestry (7.8 and 21%, respectively).

Following the Steppe pastoralist gene flow that occurred 3,500 years ago, many South Asian groups experienced shifts toward endogamy leading to strong founder effects^31^. To measure the strength of the founder events in these pastoralist groups, we applied ASCEND^38^ that utilizes genome-wide allele sharing correlations to estimate the timing and intensity of founder events. We find significant evidence for a recent, strong founder event in Toda that occurred 532 ± 28 years ago, with an estimated founder intensity of 8.5% (for comparison, this estimate is approximately 5 times higher than in Ashkenazi Jews and around 2.7 times higher than in Finns^38^) (Figure S6.1, Supplementary Note 6). In contrast, the Gujjar population has nominal evidence for a founder event with an estimated date of 3223 ± 1134 years ago and intensity of 0.8% (Figure S6.1, Supplementary Note 6).

Using coalescent simulations^39^, we modeled changes in Steppe pastoralist–related ancestry over time as the proxy for the frequency of-13.910*T and calculated the probability that Steppe-derived haplotypes could reach or exceed the observed elevated frequencies (∼65% frequency) under a range of bottleneck scenarios (Figure S6.2). Using demographic parameters inferred for Toda and Gujjar, we find that the probability of reaching the observed frequency is effectively zero (Figure S6.3). Even under extreme bottlenecks (*N_bottleneck_* = 10–5000), the probability of exceeding 65% was less than 0.1 for 7.8% Steppe ancestry (Toda) and 0.06 for 21% Steppe ancestry (Gujjar). These results indicate that drift and founder events alone cannot explain the exceptionally high-13.910*T allele frequencies, suggesting that positive selection likely played a key role in the rise of LP in these pastoralist groups.

### Evidence of Selection at the LCT locus in South Asian Pastoralists

To investigate whether the *LCT* region shows evidence of ancestry-specific enrichment, we first performed local ancestry inference for haplotypes carrying-13.910*T allele. For each haplotype, we computed pairwise similarity scores based on the number of SNP mismatches relative to reference haplotypes representing the three major ancestral components—*SAHG*, Iranian farmers, and Steppe pastoralists-related groups (see Methods, Supplementary Note 3). Each segment was then assigned to the ancestry that showed the highest similarity. This approach allowed us to estimate the proportion of Steppe-, Iranian farmer-, and *SAHG*-derived ancestry surrounding the *LCT* locus for each population. Across populations, we observed a nearly perfect correlation between Steppe ancestry and the frequency of the −13.910*T allele (r² = 0.999), indicating that Steppe ancestry explains almost all variation in allele frequency across groups (Table S3.2, Figure S3.10).

We applied the Local Ancestry Deviation test to formally assess whether the local ancestry near *LCT* deviates from the genome-wide expectation^40^. This test compares the observed proportion of Steppe ancestry in the region of interest to the genome-wide average for each population, computing a *Z*-score to quantify the deviation (see Methods). Across most geographic regions and endogamous communities, the local Steppe ancestry near *LCT* is consistent with genome-wide ancestry proportions (|*Z*| < 3; Figure 4A, Tables S4.1–S4.2). However, the two pastoralist groups—Toda and Gujjar—show significant enrichment of Steppe-derived haplotypes carrying the −13.910*T allele (*Z* = 8.74 and 4.5, respectively, Table S4.2). This excess of Steppe ancestry at the *LCT* locus in these groups provides strong evidence for recent positive selection favoring the lactase persistence allele.

**Figure 4.**
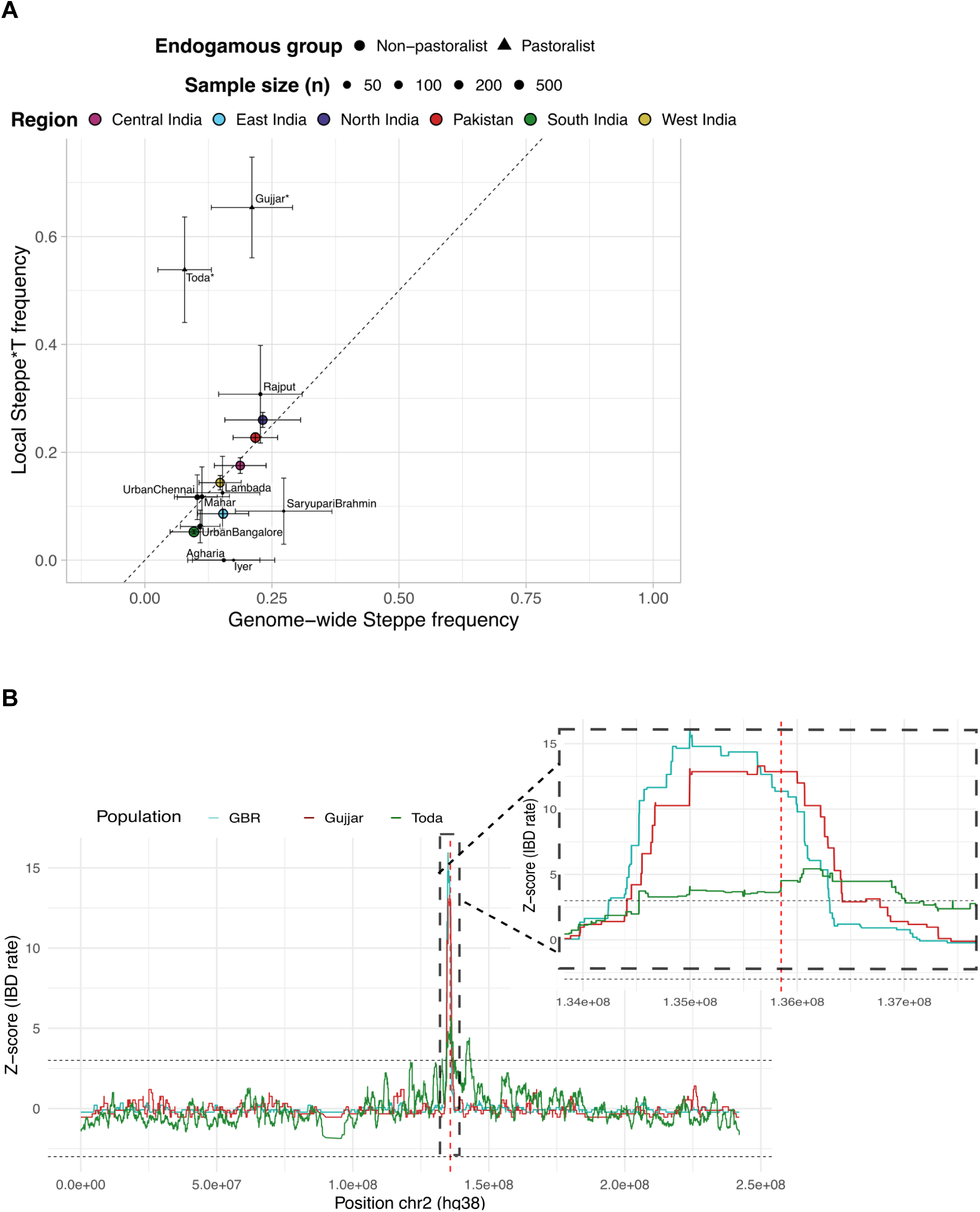
Evidence for Selection at-13.910:C>T locus for Pastoralist groups. **A.** Local Steppe*T ancestry and genome-wide Steppe pastoralist–related ancestry across regions and endogamous groups. Each dot represents the mean ancestry proportion for a group, scaled by sample size; triangles indicate pastoralist groups. Error bars show one standard deviation. These values were used in the local ancestry deviation test, stars indicate significant deviations from the genome-wide Steppe ancestry. **B.** IBD sharing rate Z-scores on chromosome 2 (main panel) and in the region surrounding the-13.910:C>T variant on chromosome 2 (inset), shown for GBR (blue), Gujjar (maroon), and Toda (green). The position of the-13.910:C>T variant is marked by a red vertical line (chr2:135,851,076; hg38).

We further examined the patterns of Identity-by-Descent (IBD) among Toda, Gujjar, and GBR. We applied hap-IBD (Zhou et al. 2020) to detect IBD segments along chromosome 2 and compared the IBD rates at the *LCT* locus relative to the background rates (Supplementary Note 6; Methods). IBD segments spanning-13.910:C>T variant were markedly elevated in all three populations—0.176 in Toda (*Z* = 4.5), 0.079 in Gujjar (*Z* = 12.9), and 0.071 in GBR (*Z* = 11.3) (Supplementary Table S3), unlike in non-pastoralist groups (|Z| < 3) (Figure 4B, Figure S7.1). Pairwise comparisons also revealed significant cross-population sharing near the *LCT* locus: GBR/Gujjar, GBR/Toda, and Gujjar/Toda pairs exceeded the chromosome-wide average by over 9–13 standard deviations (Table S7.1, Fig. S7.2). Together, these analyses reveal an unusually high rate of Steppe ancestry and elevated IBD sharing around-13.910:C>T—patterns not observed in non-pastoralist groups— consistent with recent positive selection acting on Steppe-derived haplotypes in pastoralist groups.

Finally, we estimated the selection coefficient (*s*) associated with −13.910*T in the Toda and Gujjar populations by comparing the observed allele frequencies with the expectations based on the deterministic selection model (see Methods). Assuming that the initial frequency of−13.910*T was proportional to the genome-wide Steppe pastoralist–related ancestry proportion—which is a conservative assumption, as it implies the allele was nearly fixed in the Steppe ancestors—the variant rapidly increased to its present frequency within the past 3,500 years. In the Toda, this corresponds to an estimated *s* = 0.047 (*s_d_* = 0.035, accounting for dominance of LP), and in the Gujjar, *s* = 0.033 (*s_d_* = 0.031). For comparison, present-day Europeans have an estimated *s =* 0.018^12,37^. This suggests that the strength of selection acting on this allele may have been higher in the South Asian pastoralist populations than in Northern Europeans, although this comparison does not account for demographic effects, particularly the strong bottleneck in the Toda population, which could influence allele frequency trajectories.

## Discussion

Our study provides the most comprehensive examination to date of lactase persistence in South Asia, integrating genome-wide data from around 8,000 present-day and ancient individuals across India, Pakistan, and Bangladesh. We find that the Eurasian variant (-13.910:C>T), linked to lactase persistence and under selection in Europe^3,4^, is also widespread across South Asia but with striking regional heterogeneity. Although dairy has deep cultural and dietary importance across the subcontinent, and milk consumption is common in both pastoralist and non-pastoralist groups, our findings reveal distinct trajectories of the-13.910:C>T variant across groups. The frequency of-13.910*T closely tracks Steppe pastoralist ancestry, suggesting that ancestry—not selection—explains most variation in LP in non-pastoralist South Asians. On the other hand, two pastoralist populations—the Toda from South India and the Gujjar from Pakistan—show markedly elevated frequencies, revealing a rare case of selection within the subcontinent.

The strong signal of selection in these two pastoralist groups remains striking. Both groups exhibit exceptionally high LP frequencies that cannot be explained by demographic history alone. Instead, strong enrichment of Steppe-derived haplotypes and long tracts of IBD surrounding the-13.910:C>T variant in these groups point to recent positive selection, with intensity of selection on the order of *s* ≈ 0.02–0.04 that is among the strongest known in recent human evolution^41^. Although the Toda of South India and the Gujjar of Pakistan occupy distinct ecological settings and likely experience similar episodes of nutritional stress (e.g., famine) and disease exposure as neighboring non-pastoralist populations, only these pastoralist groups show evidence of strong selection at the *LCT* locus. Future studies integrating genetic, archaeological, and anthropological data to quantify the relative consumption of raw versus fermented milk across South Asia will be crucial for elucidating the complex interplay between culture, diet, and evolution across subsistence groups.

The paradox of widespread dairy consumption but limited LP in South Asia raises important questions about gene–culture coevolution. It is possible that widespread milk fermentation practices that reduce lactose content, allowed populations to benefit from dairy without requiring genetic adaptation^42,43^. Indeed, Central Asian herders and Mongolians are mostly lactase nonpersistent, despite their significant dietary reliance on dairy products, suggesting that cultural or physiological adaptations may substitute for genetic changes^43,44^. In Europe, there is evidence that cheese and other dairy products were being consumed for nearly two millennia before-13.910*T allele was introduced through Steppe pastoralist-gene flow^13^. Analysis of UK Biobank further indicates that LP genotype was only weakly associated with milk consumption and does not exhibit consistent associations with improved fitness or health indicators^13^. Interestingly, a recent study proposes that *LCT* locus harbors several genes—*UBXN4, DARS1,* and *DARS1-AS1—*that influence immune function, not just lactose breakdown, raising the possibility that selection on-13.910*T could have been driven by both dairy consumption and immunological benefits^14^. Alternatively, physiological or microbiome-mediated mechanisms may contribute to adult milk tolerance, independent of the genetic adaptation^45–47^.

Finally, our work demonstrates the power of combining large-scale genomic surveys, local ancestry inference, and demographic modeling to disentangle the roles of gene flow, drift, and selection in shaping functional variation. The evolutionary history of-13.910*T in South Asia provides a striking example of how ancestral gene flow can introduce adaptive variation that is later shaped by local ecological pressures and cultural practices. More broadly, our findings reveal that the evolution of lactase persistence is not a single narrative of selection but a mosaic of demographic and cultural histories, each leaving a distinct genetic imprint on the human genome.

## Materials and Methods

### Dataset Description

We assembled a genome-wide dataset of 8,091 present-day and ancient genomes from India, Pakistan, and Bangladesh, including diverse timescales, geographic regions, ethnolinguistic, and subsistence groups (Supplementary Table 1.1).

### Present-day datasets

We first compiled 8,173 whole genome sequences from three recently published datasets: the Longitudinal Aging Study of India-Harmonized Diagnosis of Dementia (LASI-DAD)^23^, Genome Asia wave 1 (GAsP1)^24^ and Genome Asia wave 2 (GAsP2)^25^. We applied KING v2.3.0 (https://www.kingrelatedness.com/manual.shtml) using the “--ibdseg” option to identify and remove first-degree relatives (parent-offspring or siblings). We used the “--unrelated” tag to remove the minimum number of individuals necessary. After filtering, we retained 7,962 present-day individuals from South Asia.

#### LASI-DAD

This dataset consists of 2,679 individuals from 23 states and union territories in India, with a median sample size of 157 individuals per state. First degree relatives were removed, keeping the individual with higher coverage if a pair was identified, resulting in 2,620 individuals. Additionally, we removed 5 individuals who had missing phenotype information, resulting in a final dataset of 2,615 individuals. All genome sequences are mapped to the genome assembly GRCh38/hg38.

#### Genome Asia wave 1 (GAsP1)

This dataset consists of 688 individuals including 74 endogamous groups from South Asia. We excluded first degree relatives and groups with less than five individuals, in order to reliably infer allele frequencies. After filtering, the dataset comprised 541 individuals from 45 endogamous groups including 39 from India and 6 from Pakistan (Supplement Table S1.2). Since the dataset was initially mapped to the genome assembly GRCh37/hg19, we used *liftOver (*https://liftover.broadinstitute.org/) to convert the genomic positions to hg38.

#### Genome Asia wave 2 (GasP2)

This dataset consists of 4,806 individuals sampled from Bangladesh (Bangladesh Risk of Acute Vascular Events (BAN) (*n* = 495)), 5 groups from India (Diabetic Retinopathy Bangalore (BLR1) (*n* = 371), Retinoblastoma Bangalore (BLR2) (*n* = 9), Birbhum (BRB) (*n* = 1139), Tamil Nadu Coimbatore (COI) (*n* = 146), and Chennai (MAA) (*n* = 836)), and 2 groups from Pakistan (PROMIS Pakistan (PKN1) (*n* = 1006), and SAsianWGS Pakistan (PKN2) (*n* = 804)). All genome sequences are mapped to the genome assembly GRCh38/hg38.

### Ancient DNA

We used 181 individuals from the Allen Ancient DNA Resource (AADR)^26^ captured for the 1240K SNP panel. This includes 129 individuals from Swat Valley region in Pakistan from the Late Bronze Age to the Medieval period including 98 samples (Swat Protohistoric Grave Type - SPGT) from the Late Bronze Age/Iron Age (3000–800 BCE), 3 samples (Aligrama2_IA) from the Iron Age (970–550 BCE), 17 samples (Swat_H_Pakistan (*n* = 14), Butkara_H_Pakistan (*n* = 3)) from historical period (400 BCE–100 CE), and 8 samples (Swat_Medieval_Pakistan) from the Medieval period (1000–1650 CE) (Supplement Table 1.2). We also analyzed reference individuals from AADR, including Steppe pastoralist-related groups (Central_Steppe_MLBA (*n* = 35)) and Iranian farmer-related groups (Sarazm_EN (*n* = 2)) and Indus_Periphery_Pool (*n* = 11) that includes Indus_Periphery_West (*n* = 1)) (Supplement Table 1.2).

For inferring allele frequency and local ancestry inference, we used the imputed AADR^35^ dataset that was generated using GLIMPSE (v1.0.0) with the high coverage 1000 Genomes Project (1000G) phase 3 sequences^48^ as the reference panel, keeping only sites that passed all quality control filters from gnomAD (v2.1.1)^49^. Each individual was phased separately and only individuals with imputation quality score (IQS) of greater than 0.9 were retained, where IQS = average(*GP1* | *GT* = 1), where *GT* is the most likely genotype based on the imputed genotype posterior *GP* = (*GP0, GP1, GP2*) and ∑*GP* = 1. For our analyses, we only kept variants with medium to high quality as defined by the authors of the study^35^. Following imputation, we retained 10,545,635 variants, with 881,335 variants on chromosome 2.

### Phasing

We used SHAPEIT4^50^ (without reference data) with the HapMap recombination map^51^ to phase the genomes from present-day South Asian individuals. Each dataset (LASI-DAD, GAsP1, GAsP2) was phased independently to preserve as much genetic variation as possible. This approach has been shown to work reliably for South Asians yielding a low phase switch error rate of 1.13% in pedigrees^23^.

### Principal component analysis

To characterize the population structure in South Asia and its relationship to other world-wide populations, we combined our South Asia dataset with unrelated individuals from 1000G (The 1000 Genomes Project Consortium, 2015) including individuals from Europe (*n* = 526), Africa (*n* = 532), and East Asia (*n* = 512) and Human Genome Diversity Panel (HGDP) ^28^ including individuals from Europe (*n* = 155), Middle East (*n* = 161), Africa (*n* = 104) and East Asia (*n* = 223). For the 1000G Africans, we excluded ASW (Americans of African Ancestry in Southwest USA) and ASC (African Caribbean in Barbados) as they are recently admixed. We ran *smartpca*^52^ with 10,175 individuals and 887,967 SNPS that are common across all datasets, both with sub-Saharan African outgroups (Figure S1.1) and without African outgroups (9,368 individuals) (Figure 2B).

## ADMIXTURE

We ran *ADMIXTURE*^53^ using the same dataset as *smartpca* including South Asians, West Eurasians, East Asians, and sub-Saharan Africans. We varied the number of clusters (*K*) between 2 to 6 and performed cross validation by running the method ten times (--*cv=10*) (Supplementary Figure S1.3, S1.4, Supplementary Note 1).

### qpAdm analysis

To model the ancestry in South Asia, we used *qpAdm*^33,34^ which compares allele frequency correlations between a population of interest and a set of reference and outgroup populations. We excluded individuals with >1% sub-Saharan African-related component from ADMIXTURE analysis (based on the *K=2* analysis), which removed 156 Indian individuals and 329 Pakistani individuals (Supplementary Table 1.1). For individuals on the Indian cline, we applied the three-way model including Iranian farmer-related, Steppe pastoralist-related, and South Asian hunter-gather (SAHG)-related ancestries^31^. We used *Indus_Periphery_West* from AADR (*n* = 1; Supplementary Table S2.1) as a proxy for Iranian farmer-related ancestry, *Central_Steppe_MLBA* from AADR (n=35; Supplementary Table S2.1) as a proxy for Steppe pastoralist-related ancestry, and present-day Onge from GAsP1 (n=15) as a proxy for SAHG-related ancestry (Narashiman et al, 2019). For this model, we used the following outgroups: *Ethiopia_4500BP.SG, WEHG, EEHG, Anatolia_N, ESHG, Dai.DG, Iran_GanjDareh_N, Russia_Samara_EBA_Yamnaya, WSHG*. For individuals that fall outside the Indian cline and cluster toward the East Asia-related groups in PCA, primarily from Bangladesh (*n* = 522) and North-East India (*n* = 71), we also applied a four-way model with addition of Han Chinese (CHB) from 1000G as a proxy for East Asian-related ancestry. We considered that the model is a good fit to the data if all ancestry coefficients > 0 and the model’s p-value > 0.01.

### Measuring allele frequency of LP-associated variants

For each LP-associated variant, we estimated the allele frequency using vcftools “--freq” option for all present-day unrelated individuals (*n* = 7,962). We clustered individuals by endogamous group or geographic region of sampling (i.e. Pakistan, Bangladesh, or region of India). For each allele frequency estimate, we computed 95% confidence intervals by estimating the standard error (SE) using bootstrapping resampling with 1,000 replicates. We also calculated the standard error assuming a binomial distribution.

### Measuring LP Frequency

As-13.910:C>T confers a dominant phenotype, we estimated the LP frequency by weighting the heterozygous (CT) and homozygous (TT) genotypes carrying the alternate allele equally. Specifically, we calculated:

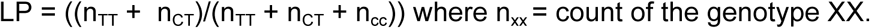

To this end, we determined the count of the heterozygous and homozygous individuals using plink1.9 “--recode-rlist”. For each LP frequency estimate, we computed 95% confidence intervals using bootstrapping resampling with 1,000 replicates.

For the ancient individuals, we estimated the LP frequency using read count and imputed data, after excluding individuals with less than 3 reads overlapping the 13.910:C>T variant. After filtering, we retained SPGT (*n* = 63), Aligrama2_IA (*n* = 2), Barikot_H_Pakistan (*n* = 2), Butkara_H_Pakistan (*n* = 2), Swat_H_Pakistan (*n* = 11), and Swat_Medieval_Pakistan (*n* = 2). We note other LP-associated variants were not captured in the 1240K AADR dataset. We analyzed the raw BAM files of each individual using IGV_2.9.4 ^54,55^ and counted the number of reads that supported the reference and alternate allele at-13.910:C>T variant. We compared these estimates with the frequencies based on the imputed genotypes, after retaining high quality variants^35^. We obtained qualitatively similar results for both methods (Supplementary Table 2.1).

### Measuring genetic diversity

To quantify genetic diversity, we computed per-SNP entropy for all SNPs in the core region or extended region using the formula

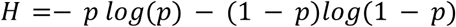

where *p* is the non-reference allele frequency at a given SNP. For each region, we then computed an average value. We stratified by the presence or absence of the derived allele, retaining-13.910*T (n = 2,065) or-13.910*C (n = 13,859). We applied the exact Wilcoxon rank-sum test to assess the statistical significance between diversity levels across haplotype groups.

### Demographic simulations for Toda and Gujjar

We conducted coalescent simulations using *msprime*^39^ to model the demographic history of Toda and Gujjar populations, incorporating gene flow from Steppe pastoralist-related groups followed by founder events (Figure S6.2). We simulated two ancestral populations — Pop1 and Pop2 — representing ancestral South Asians and ancestral Steppe pastoralists, which diverged 30,000 years ago. In the simulation, gene flow from the Steppe pastoralist-related population into SAHG was set to occur 3,500 years ago^31^, producing the contemporary South Asian population analyzed here. Both ancestral populations were assigned an initial effective size of 10,000. To explore uncertainty in founder event strength, we varied N_bottleneck_ = [5000, 1000, 500, 100, 90, 70, 50, 30, 10], alongside a constant-size model. In the ‘Bottleneck’ scenario, Ne was reduced to N_bottleneck_ starting 120 generations ago (t_start_ = 120) and lasting until 20 generations ago (t_end_ = 20), after which it returned to 10,000 (Figure S6.2). For each scenario, we simulated human chromosome 14 (106,880,170 bp) using recombination rates from the HapMap map ^51^. From the resulting population, we sampled 500 chromosomes and used tree sequences to map the locations, lengths, and frequencies of Steppe pastoralist-related ancestry segments, focusing on the proportion of regions inherited from Steppe pastoralist-related gene flow exceeding 65%. We conducted 100 independent simulations and bootstrap resampling to estimate means and standard errors of this proportion.

### Examining the relationship between ancestry and allele frequency

We examined the relationship between ancestry and allele frequency of-13.910*T frequency using PCA loadings, genome-wide ancestry proportions inferred using qpAdm and local ancestry inference.

#### Using PCA loadings

We used 7,391 individuals after excluding individuals with >10% sub-Saharan African ancestry-related components from ADMIXTURE (Figure S1.4 A, Figure S1.5). We used the SNP weights for PC1 and PC2 from Figure 2B as a proxy for genome-wide variation in ancestry in South Asians. We binned individuals (excluding endogamous groups) according to their PC1 or PC2 loadings, ensuring an even number of individuals in each bin (*n* ∼ 369). For each bin, the mean allele frequency of-13.910*T was calculated. Using the binned data, we performed a linear regression between the PCA loadings and-13.910*T allele frequency using linear model (*lm*) in R. We report the correlation coefficient and *p*-value for the regression model. For each endogamous group, we calculated the mean PC loading of the group and the mean allele frequency and projected these samples onto the regression line (Figure S5.1). We calculated the *Z*-score for each group to determine the deviation from the regression line, with values of |*Z*| > 3 indicating a significant deviation. We also examined the relationship between ancestry and LP frequency instead of-13.910*T allele frequency (Supplementary Note 5; Figure S5.2).

#### Using genome-wide ancestry proportions based on *qpAdm*

For individuals on the Indian cline (*n* = 4,946), we used the *qpAdm* ancestry coefficients for Iranian farmer-related, Steppe pastoralist-related, and SAHG-related ancestries (for individuals with *qpAdm p*-value > 0.01 and all ancestry coefficients > 0). For each ancestry component, individuals were binned to ensure an even number of individuals per bin (n = ∼247). The mean allele frequency of-13.910*T was calculated for each bin. Using a linear regression model in R, we estimated the correlation coefficient between the median ancestry coefficient for each bin and-13.910*T allele frequency. For each endogamous group, we calculated the mean ancestry proportion for each ancestral group and the mean allele frequency and projected these samples onto the regression line. We also conducted this analysis with individuals that fall off the cline (*n* = 477) but can be modeled using the three-way or 4-way model (including East Asian-related ancestry) in qpAdm fit the ancestry model (Supplementary Note 5; Figure S5.3). We also conducted the regression analysis with LP frequency as described in Supplementary Note 5 (Figure S5.4).

### Local ancestry inference at the core haplotype

To infer local ancestry at the core haplotype, we compared phased haplotypes in present-day South Asians to ancestral populations—SAHG, Iranian-related farmers, and Steppe pastoralists. For each South Asian*T haplotype, we computed pairwise similarity scores based on the number of SNP mismatches relative to reference haplotypes as follows:

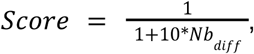

*Nb_diff_* is the number of SNP differences between the test and each reference haplotype. This yields scores that decrease sharply with more differences (e.g., 0 differences = 1.00, 1 = 0.09, 2 = 0.05, 3 = 0.03, etc.).

To estimate the probability of deriving ancestry from a particular ancestral population, we computed the average similarity score across individuals in an ancestral population 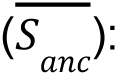

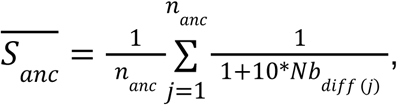

Where 𝑛𝑎𝑛𝑐 is the number of reference individuals in the population 𝑎𝑛𝑐.

We then normalized the averages estimates across all ancestral populations to infer the probability of deriving ancestry from each group:

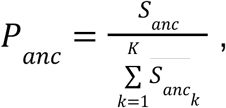

We note the probabilities for all ancestral populations should sum to 1 (Figure S4.1). We assigned the local ancestry to the ancestral population with the highest normalized score exceeding a confidence threshold of 0.80, otherwise, no ancestry was assigned.

#### Comparison with allele frequency

For individuals on the Indian cline (*n* = 4,830), we also examined the relationship between local ancestry at the core haplotype and-13.910*T allele frequency. Local ancestry assignments for-13.910*T haplotypes were obtained as described in *Local ancestry at the core haplotype*. To evaluate whether these measures differed significantly, Z-scores were computed as:

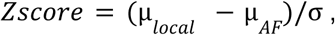

where µ_𝑙𝑜𝑐𝑎𝑙_ is the average local Steppe*T frequency and µ_𝐴𝐹_-13.910*T allele frequency; and the standard error is estimated as the 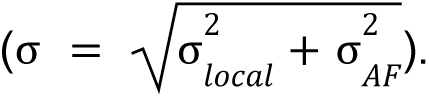 Standard errors for both estimates were derived from binomial sampling to account for stochastic variation arising from finite sample sizes.

### Local Ancestry Deviation (LAD) test

For each group, we computed *Z*-scores to compare the inferred local Steppe ancestry proportion with the expectation based on genome-wide Steppe ancestry proportion estimated using qpAdm (|Z| > 3 was considered significant). Z-scores were computed as:

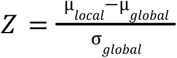

where µ_𝑙𝑜𝑐𝑎𝑙_ is the inferred local Steppe*T haplotype proportion, µ_𝑔𝑙𝑜𝑏𝑎𝑙_ is the mean genome-wide Steppe ancestry proportion per individual (based on *qpAdm*), and σ_𝑔𝑙𝑜𝑏𝑎𝑙_ is the corresponding standard deviation. Because the endogamous groups are highly homogeneous, the empirical distribution of genome-wide ancestry proportions does not adequately capture uncertainty and thus, we estimated standard errors in endogamous groups using a binomial approximation to account for sampling uncertainty.

### Patterns of Identity-by-Descent sharing

We identified IBD segments using *hap-IBD*^56^ with the following parameters: min-seed=0.5, max-gap=1000, min-extend=0.5, min-output=0.5 min-markers=100, min-mac=2 and nthreads=1. We used the deCODE recombination map^57^. We retained shared IBD segments that were at least 0.5 cM and 1Mb in length. We examined IBD sharing within and between the target populations: GBR, Gujjar and Toda. For within-population comparisons, average IBD sharing was computed across all possible pairs of individuals within that population. For between-population comparisons, average IBD sharing was computed across all possible pairs of individuals with one individual from each population. To compute average IBD sharing across chromosome 2, we considered 1,000 bp windows with a callability greater than 0.8 based on the strict callability mask from the 1000 Genomes Project^48^. This resulted in a total of 163,181,000 bp after filtering for chromosome 2. Local IBD sharing at the core haplotype region was compared to chromosome-wide IBD sharing using Z-scores.

### Estimations of selection coefficient

We estimated the strength of selection (*s*) acting on-13.910*T by comparing its observed frequency (*p*) change across generations to the expectations under deterministic selection models^58^. We assume the initial allele frequency *p_0_*corresponds to the genome-wide proportion of Steppe pastoralist-related ancestry, *t* represents the number of generations since the gene-flow and *p_t_* is the-13.910*T frequency at time *t* (present-day). Note, this approach implicitly assumes that the allele was fixed in the Steppe ancestral population—a conservative assumption, as lower initial frequencies would imply stronger selection (*higher s*).

First, we assumed additive effects (no dominance) and used the standard logistic model of directional selection, which yields a closed-form estimator for *s*:

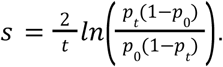

Accounting for the dominance (*h* = 1) of LP:

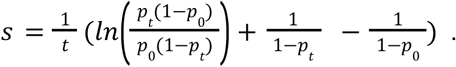

For the estimates in the Toda and Gujjar populations, we assumed *t* = 125 generations based on the inferred timing of Steppe pastoralist gene flow in ancient DNA^31^. For Toda and Gujjar, we set the initial frequencies as *p_0_*= 7.8% and 21%, respectively, with corresponding observed present-day frequencies of *p_t_* = 61% and 67.5%, respectively.

## Data and code availability

Whole genome sequences for present-day South Asians were downloaded from:

LASI-DAD data was accessed through the National Institute on Aging Genetics of Alzheimer’s Disease Data Storage Site (NIAGADS) under the accession NIAGADS: *ng00067*. These data can be obtained through the GCAD at the University of Pennsylvania: https://dss.niagads.org/documentation/data-application-and-submission/application-instructions/

GAsP1 data was accessed through the European Genome-Phenome Archive (EGA), accession number EGA: EGAS00001002921. Individual-level vcf files accessed via request forms are also available from https://browser.genomeasia100k.org.

GAsP2 data was accessed from the SRA under NCBI BioProject PRJNA476341 and individual-level VCF files were obtained directly from the authors upon request.

1000 Genomes Project and Human Diversity Genome Project. These data were obtained from the International Genome Sample Resource (IGSR): https://www.internationalgenome.org/data.

Ancient DNA data was downloaded from the Allen Ancient DNA Resource (version 54): https://dataverse.harvard.edu/dataset.xhtml?persistentId=doi:10.7910/DVN/FFIDCW.

Scripts and pipelines to reproduce the analysis are available through github: https://github.com/MoorjaniLab/LP_SouthAsia.

## Acknowledgements

We thank Monty Slatkin, Chris Martin, Sophie Joseph, Jeremy Choin, Bárbara Sousa da Mota, Nick Patterson, and members of the Moorjani lab for helpful discussions and comments on the manuscript. We thank Ali Akhbari and David Reich for sharing imputed AADR genotypes. We also thank Claire McEvoy for insights on dairy consumption in India from the LASI-DAD food survey. The Longitudinal Aging Study in India, Diagnostic Assessment of Dementia data (doi.org/10.25549/5hhx-s820) is sponsored by the National Institute on Aging (grant number R01AG051125, RF1AG055273, U01AG065958) and is conducted by the University of Southern California. JL was supported by National Institute on Aging, National Institutes of Health R01AG051125 and U01AG064948. KT was supported by CSIR-Bhatnagar Fellowship [GDA No. 24 (90807)], Council of Scientific and Industrial Research, Ministry of Science and Technology, Government of India. PM, MM and EK were supported by the Burroughs Wellcome Fund Careers at the Scientific Interface and NIH R35GM142978 awarded to PM.

