## Supplementary Information for "Revisiting the Evolution of Lactase Persistence: Insights from South Asian Genomes"

### Supplement Information

**<u>Table of Contents</u>**

**Supplementary Note 1. Data Description and Population Structure in South Asia 2**
1.1 Principal Components Analysis (PCA) 2
1.2 ADMIXTURE 4
1.3 Modeling Ancestry of South Asians 8
**Supplementary Note 2. Allele frequency Patterns of LP-Associated Variants in South Asians 11**
2.1 Allele frequency of LP-associated variants 11
2.2 Frequency of -13.910:C>T, accounting for dominance 15
2.3 Linkage Disequilibrium Patterns of LP-Associated Variants in South Asia 18
**Supplementary Note 3. Steppe origin of -13.910*T in South Asia 20**
3.1 Core Haplotype 22
Haplostrips 24
3.2 Extended Haplotype 27
**Supplementary Note 4. Local Ancestry Inference 30**
4.1 Method 30
4.2 Relationship between local ancestry and -13.910*T frequency 31
4.3 Local Ancestry Deviation test 33
**Supplementary Note 5. Relationship between Ancestry and frequency of LP-associated variants 36**
5.1 Relationship between ancestry (using PCA loadings) and allele frequency of -13.910:C>T 36
5.2 Relationship between genomewide ancestry (using qpAdm) and allele frequency of -13.910*T 39
5.3 Relationship between genome-wide ancestry and local ancestry at -13.910:C>T locus 45
**Supplementary Note 6. Simulations to Examine the Role of Demographic History in Shaping LP Prevalence in Toda and Gujjar 47**
6.1 Steppe Pastoralist-Related Gene Flow 47
6.2 History of Founder Events 47
6.3 Simulation Scenarios 48
Model Parameters 50
6.4 Distribution of Steppe Pastoralist-Related Ancestry in Simulations 50
**Supplementary Note 7. Detecting Signatures of Natural Selection Using Identity-By-Descent Segments 52**
7.1 IBD Sharing within a Population 52
7.2 IBD Sharing between Populations 55
**References 57**

### Supplementary Note 1. Data Description and Population Structure in South Asia

#### 1.1 Principal Components Analysis (PCA)

To understand the population structure in South Asia, we applied *smartpca*[1] to genetic data from unrelated individuals from our South Asian dataset (LASI-DAD, GAsP1 and GAsP2) merged with 1000G[2] and the Human Genome Diversity Panel (HGDP)[3], after removing first-degree relatives identified with *KING* (see Methods). Our analysis included 7,962 individuals from South Asia (Supplementary Table 1.1), 1,570 individuals from 1000G including Europeans (EUR, *n* = 526), Africans (AFR, *n* = 532), and East Asians (EAS, *n* = 512) and 643 individuals from HGDP including Europeans (*n* = 155), Middle Easterners (*n* = 161), Africans (*n* = 104) and East Asians (*n* = 223). As seen in previous studies, PC1 separates Africans and non-Africans, while South Asians fall along a cline of relatedness to West Eurasians (Europeans and Middle Easterners) and East Asians along the PC2[4–6] (Figure S1.1). Around 30 individuals from Pakistan exhibit a shift towards the African cluster on PC1, indicating some sharing with sub-Saharan African groups. This signal is likely attributable to some recent Middle Eastern ancestry in some Pakistani individuals, as suggested by earlier studies[5,7].

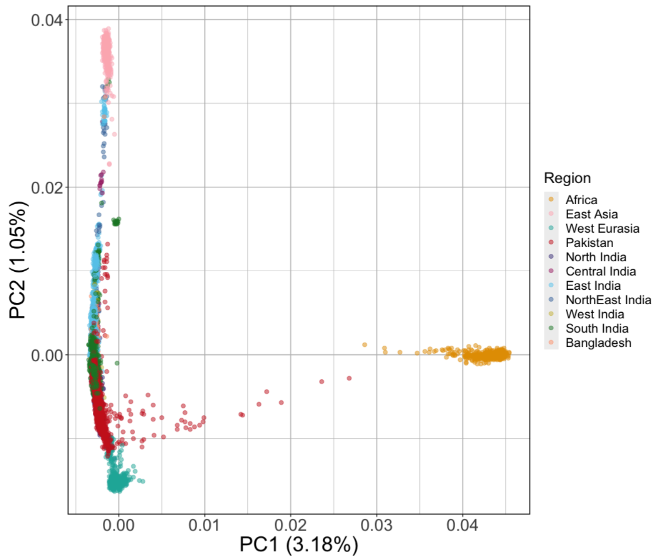


**Figure S1.1: Principal Components Analysis of South Asians from LASI-DAD, GAsP1, and GAsP2 with West Eurasians, East Asians, and Africans from 1000G and HGDP.** PC1 and PC2 are plotted against each other, with African populations shown in yellow, West Eurasian populations shown in teal, East Asian populations in pink. South Asian individuals from Pakistan, Bangladesh, and each geographic region of India are shown in different colors (based on the sampling locations shown in Figure 1A).

Previous studies have shown that individuals from endogamous communities in South Asia cluster closely with others from the same group and remain genetically distinct from other groups, reflecting a history of recent founder events[8–11]. To characterize the population structure among the endogamous groups in our dataset, we performed PCA with 541 individuals (45 groups) from GAsP1 and 842 West Eurasian- and 735 East Asian-related individuals from 1000G and HGDP. The overall population structure across all endogamous groups mirrors the population structure observed in Figure 1B. Notably, individuals within endogamous communities tend to be genetically homogeneous and generally cluster by sampling location (Figure S1.2 A). Individuals sampled from Pakistan and North India fall along one end of the Indian cline, while individuals from South India are at the other end. Groups from Bangladesh and East / North-East India cluster between the Indian cline and East Asian-related populations. Andamanese and Nicobarese Islanders form distinct clusters reflecting their population isolation from other groups. Importantly, clustering does not reflect mode of subsistence; for example, pastoralist groups such as the Toda and Gujjar cluster according to their geographic location, rather than lifestyle.

Finally, we examined how ancient South Asian individuals relate to other worldwide populations. To this end, we projected 129 ancient individuals (dated: ~1491 BCE to 1650 CE) sampled from the Swat Valley in Pakistan on the PCA described in Figure 1B. As expected from their geographic origin, ancient South Asians cluster with present-day individuals from Pakistan and North India along the Indian cline and show no significant clustering by sampling age (Figure S1.2 B).

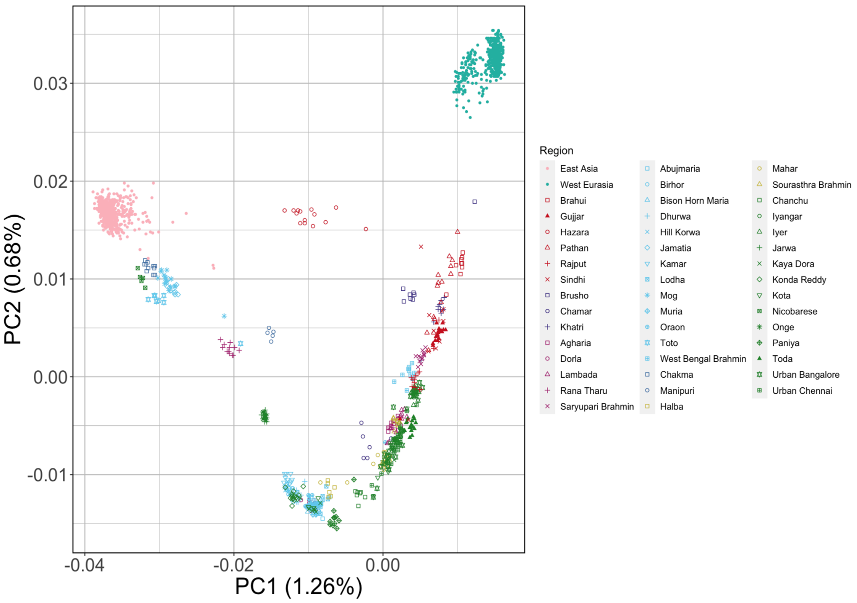

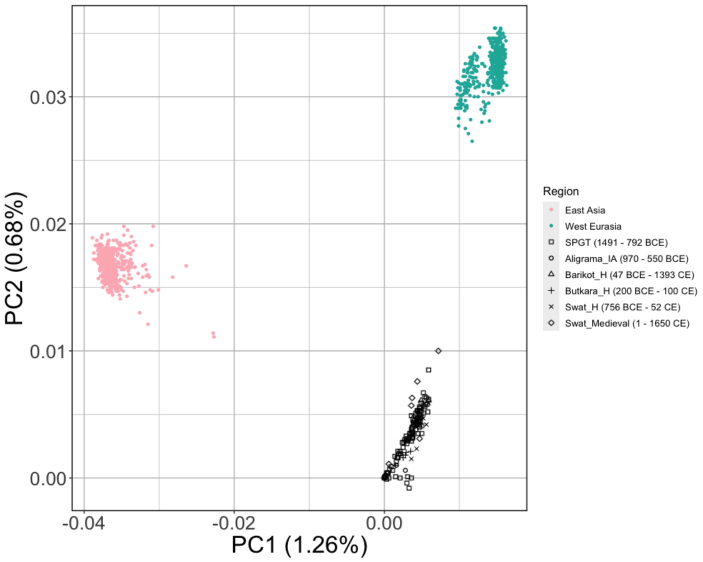


**Figure S1.2: Principal Components Analysis of GAsP1 endogamous groups and ancient South Asians with West Eurasians and East Asians.** PC1 and PC2 are plotted against each other, with West Eurasian populations shown in turquoise and East Asian populations in pink.

A. **Endogamous Groups** Each endogamous group is represented by a different symbol and colored by their geographic location: Pakistan, North India, Central India, East India, North-East India, West India, South India, or Bangladesh.
B. **Ancient South Asians** The ancient South Asian individuals (SPGT (~1491 - 800 BCE), Aligrama_IA (~970 - 550 BCE), Barikot_H (~47 BCE - 1393 CE), Butkara_H (~200 BCE - 100 CE), Swat_H (~400 BCE - 52 CE), and Swat_Medieval (~1 - 1650 CE)) are plotted in black, with different symbols for each group.

#### 1.2 ADMIXTURE

We performed unsupervised clustering using ADMIXTURE[12] using the same dataset described earlier (Figure S1.1) which includes 7,962 South Asians, 1,570 individuals from 1000G, and 643 individuals from HGDP. We varied the number of clusters (*K*) between 2 to 6 and performed cross validation with ten replicates (--*cv=10*) to evaluate model fit and identify the most appropriate value of *K*. The lowest cross validation error (=0.3402) was observed for *K*=5 (Figure S1.3).

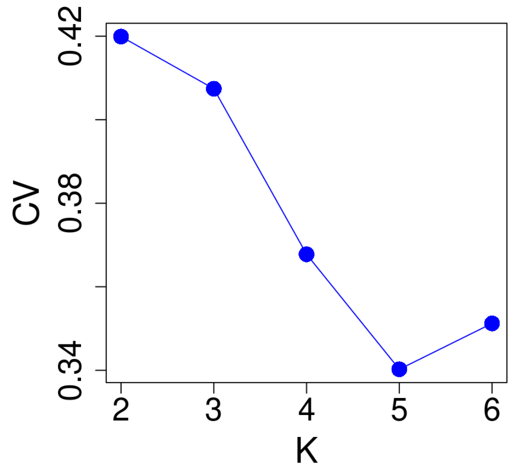


**Figure S1.3 Cross validation errors from ADMIXTURE analysis**. We performed ADMIXTURE analysis using the merged dataset of LASI-DAD, GAsP1, GAsP2, 1000G (AFR, EAS, and EUR) and HGDP (Africans, West Eurasians, and East Asians) and showed cross-validation error for the clusters (*K*) 2-6.

At *K*=2, we find individuals separated into two clusters, one consisting predominantly of individuals from Africa and the other containing non-Africans (Figure S1.4 A). A subset of individuals from Pakistan (*n* = 32) carry ≥10% ancestry from the African-related cluster, reaching as high as 62.5%. These are the same individuals noted to be drifted towards the African-cluster in the PCA in Figure S1.1 (Figure S1.5). We excluded these 32 individuals from downstream ancestry analysis. Within South Asia, individuals from Bangladesh, India, and Pakistan have variable amounts of East Asian- and West Eurasian-relatedness. At *K*=4 and *K*=5, we observe ancestry components which are predominantly seen in South Asian and East Asian-related individuals, likely reflecting the ancient Ancestral South Indians (AASI) ancestry in South Asia (Figure S1.4 C & D).

**A.** *K*=2

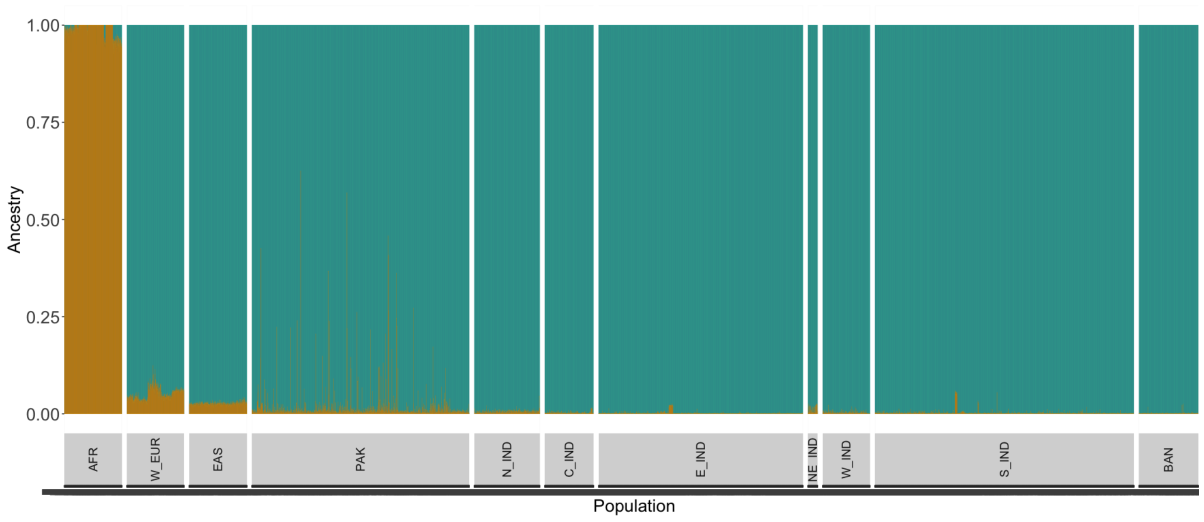

**B.** *K*=3

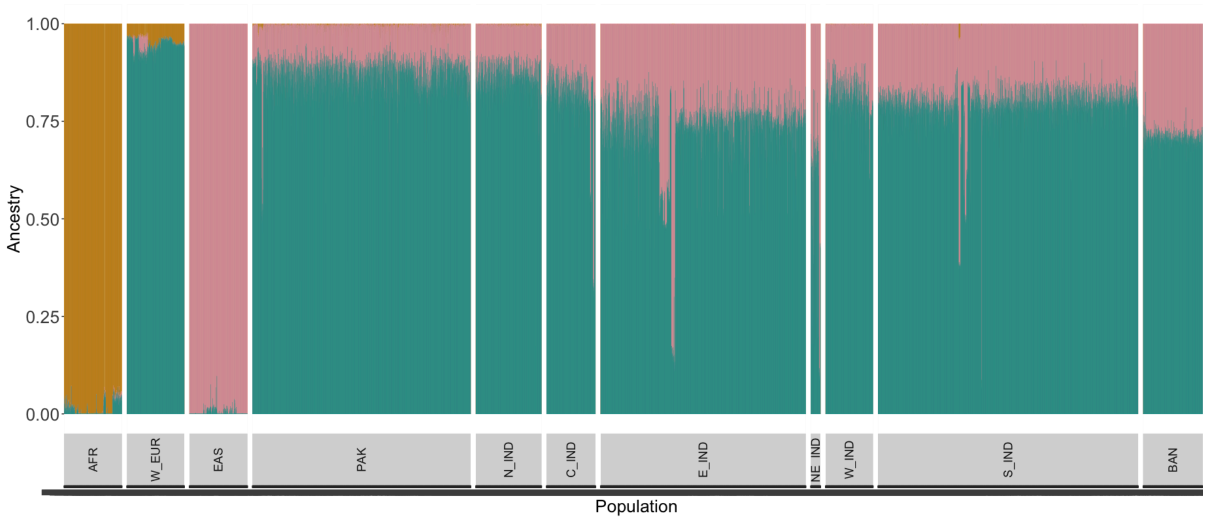
1.00
0.75
0.50
0.25
0.00
Ancestry
AFR
W_EUR
EAS
PAK
N_IND
C_IND
E_IND
NE_IND
W_IND
S_IND
BAN
Population

**C.** *K*=4

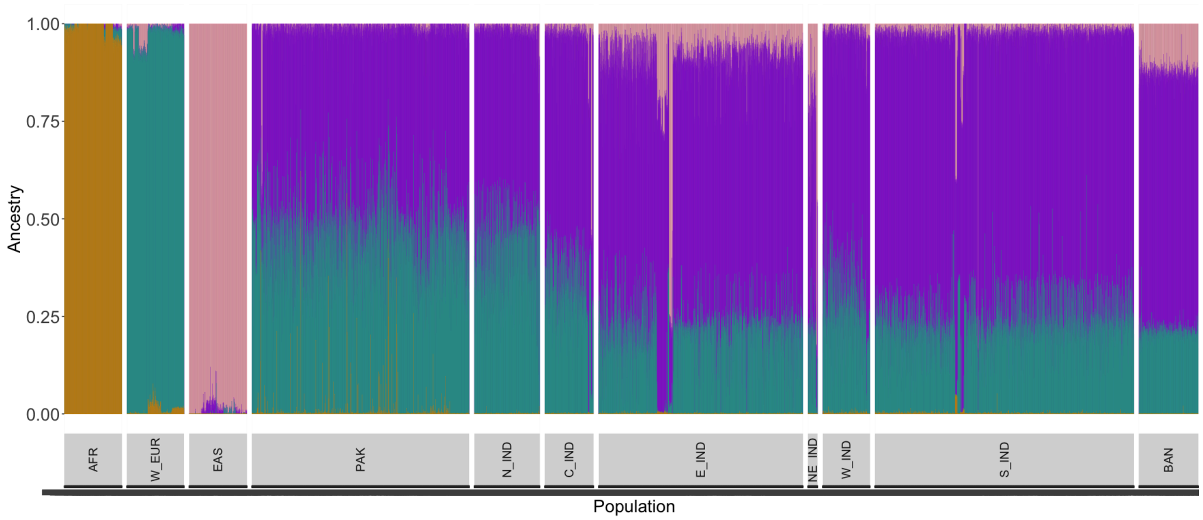
1.00
0.75
0.50
0.25
0.00
Ancestry
AFR
W_EUR
EAS
PAK
N_IND
C_IND
E_IND
NE_IND
W_IND
S_IND
BAN
Population

**D.** *K*=5

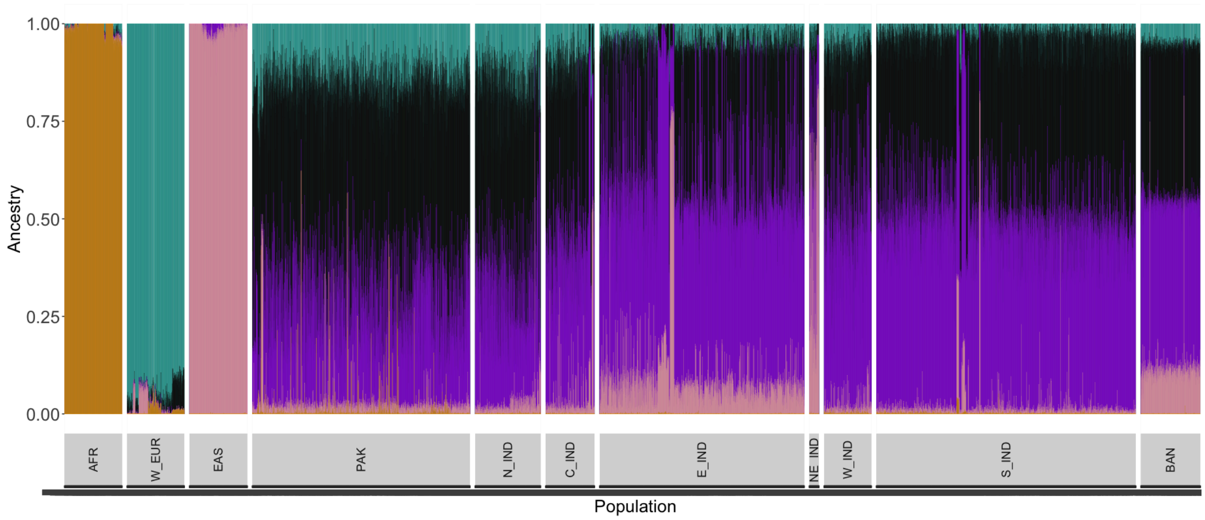


**Figure S1.4 ADMIXTURE analysis of LASI-DAD, GAsP1, and GAsP2 with 1000G West Eurasians, East Asians, and Africans.** We performed ADMIXTURE, by varying the number of clusters (*K*) from 2-5 (A-D). Each vertical line represents one individual and the different colors are the ancestry proportions from the different clusters inferred with ADMIXTURE. Individuals are grouped by their geographic region of sampling: Africa (AFR), West Eurasia (W_EUR), East Asia (EAS), Pakistan (PAK), North India (N_IND), Central India (C_IND), East India (E_IND), North-East India (NE_IND), West India (W_IND), South India (S_IND), and Bangladesh (BAN).

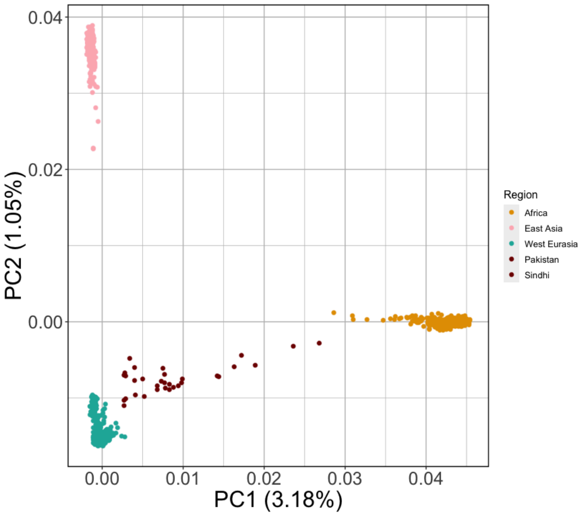


**Figure S1.5 South Asian individuals with >10% African ancestral component from ADMIXTURE shown in PCA of South Asians with West Eurasians, East Asians, and Africans.** Individuals with >10% African ancestral component when *K* = 2 from ADMIXTURE (Figure S1.4 A) are projected on the same PCA conducted in Figure S1.1 to understand how these individuals relate to Africans. The individuals are shown with the same colors associated with geographic regions and symbols for endogamous groups as Figure S1.1 and Figure 1B.

#### 1.3 Modeling Ancestry of South Asians

To model the ancestry in South Asia, we used *qpAdm*[13,14] which compares allele frequency correlations between the population of interest and a set of reference and outgroup populations. We excluded 32 individuals with >= 10% sub-Saharan African-related ancestry observed in ADMIXTURE (*K*=2) (Supplementary Table 1.1). We separately analyzed individuals that fall along the Indian cline ($n$ = 5,948) and outside the Indian cline ($n$ = 1,982). We defined the Indian cline based on PCA results (Figure 1B) by fitting a regression line that captures the diversity among South Asians and retaining mean +/- 2 standard deviations along PC1 and PC2.

We modeled individuals on the Indian cline as a mixture of three ancestral groups related to ancient Iranian farmers, Eurasian Steppe pastoralists, and South Asian hunter-gatherers (SAHG). We used *Indus_Periphery_West (I8726)* from AADR ($n$ = 1) as a proxy for Iranian farmer-related ancestry, *Central_Steppe_MLBA* from AADR ($n$ = 35) as a proxy for Steppe pastoralist-related ancestry, and present-day Andaman Islanders (Onge, $n$ = 15) from GAsP1 as a proxy for SAHG-related ancestry[4] (Supplementary Table S2.1). We used the following outgroups: *Ethiopia_4500BP.SG, WEHG, EEHG, Anatolia_N, ESHG, Dai.DG, Iran_GanjDareh_N, Russia_Samara_EBA_Yamnaya, WSHG*. We considered the model to be a 'good fit' if all three ancestry coefficients were non-zero and $p$-value > 0.01 in *qpAdm* analysis.

For individuals on the Indian cline, we find this model provides a good fit to data from 4,946 individuals (>83%). We infer the Iranian farmer-related ancestry ranges from 27.5-69.7%, Steppe pastoralist-related ancestry ranges from 0.3-46.4%, and SAHG-related ancestry ranges from 10.3-64.1%. Individuals from North India and Pakistan have higher mean Steppe pastoralist-related ancestry proportions (on average >20%) than individuals from South India (9.5%; $p$-value < 1e-20), as observed in previous studies[15]. Conversely, individuals from South India have higher mean SAHG-related ancestry (42.4%) than Pakistan and North India (28.9%; p-value < 1e-20) (Supplementary Table 1.2). The proportion of mean Iranian farmer-related ancestry remains does not show a clear geographic gradient (Figure S1.6, Supplementary Table 1.2).

Next we applied *qpAdm* to individuals that fall outside the Indian cline. We first applied the three-way model and found 580 individuals are a good fit. As many off-cline individuals are shifted towards East Asians in PCA (Figure 1B), particularly those from East India and Bangladesh, we also tried a four-way model with the addition of East Asian-related ancestry (Han Chinese from 1000G, CHB). This model fits >80% of the individuals from Bangladesh ($n$ = 495), and around 56% of the individuals from North-East India ($n$ = 71) (Supplementary Table 1.2). The remaining 836 individuals do not fit the three-way or four-way model.

We also separately modeled the ancestry composition of the 535 individuals from 44 endogamous communities (excluding Onge, as they are being used as SAHG proxies). We applied the three-way model for individuals on the Indian cline and four-way model for off-cline groups (Chakma, Jamatia, Manipuri, and Toto). We performed the analysis separately for each individual and reported average estimates for each group (retaining only individuals where the model is a good fit (see Methods)). On the whole, the three-way model provides a good fit for 295 individuals (~55%) across the 44 endogamous groups. Only 21 of the 44 groups fall on the Indian cline, and of these, 14 have >= 80% of the individuals fit the three-way model in their respective group. (Supplementary Table 1.2). In these 14 groups, the mean Iranian farmer-related ancestry ranged from 33.73–58.85%, mean SAHG-related ancestry ranged from 28.87–55.15% and mean Steppe pastoralist-related ancestry ranged from 7.91–27.31% (Supplementary Table 1.2; Figure S1.6). The three-way model did not provide a good fit for four of the groups from Pakistan (<= 50%), likely due to Middle Eastern gene flow and African-related ancestry (Figure S1.1, Figure S1.3). The four-way model with East Asian-related ancestry (Han Chinese from 1000G, CHB) provided a good fit for around 18-80% of the individuals belonging to the four off-cline groups.

Finally, we modeled the ancestry of ancient South Asians from the Swat Valley region in Pakistan (*SPGT*, *Aligrama2_IA*, *Swat_H_Pakistan*, *Butkara_H_SwatValley*, *Barikot_H*, and *Swat_Medieval Pakistan*) dated to ~1491 BCE to 1650 CE, using the same three-way model as applied to present-day South Asians on the Indian cline. This model fit data from ~87% of the individuals, with inferred ancestry proportions ranging from 1.7–24.9% SAHG-related, 36.6–86.1% Iranian farmer-related, and 0.8–50.7% Steppe pastoralist-related. These estimates fall within the range of estimates observed in present-day South Asians, and are comparable to those observed in present-day individuals from Pakistan and North India.

**A.**

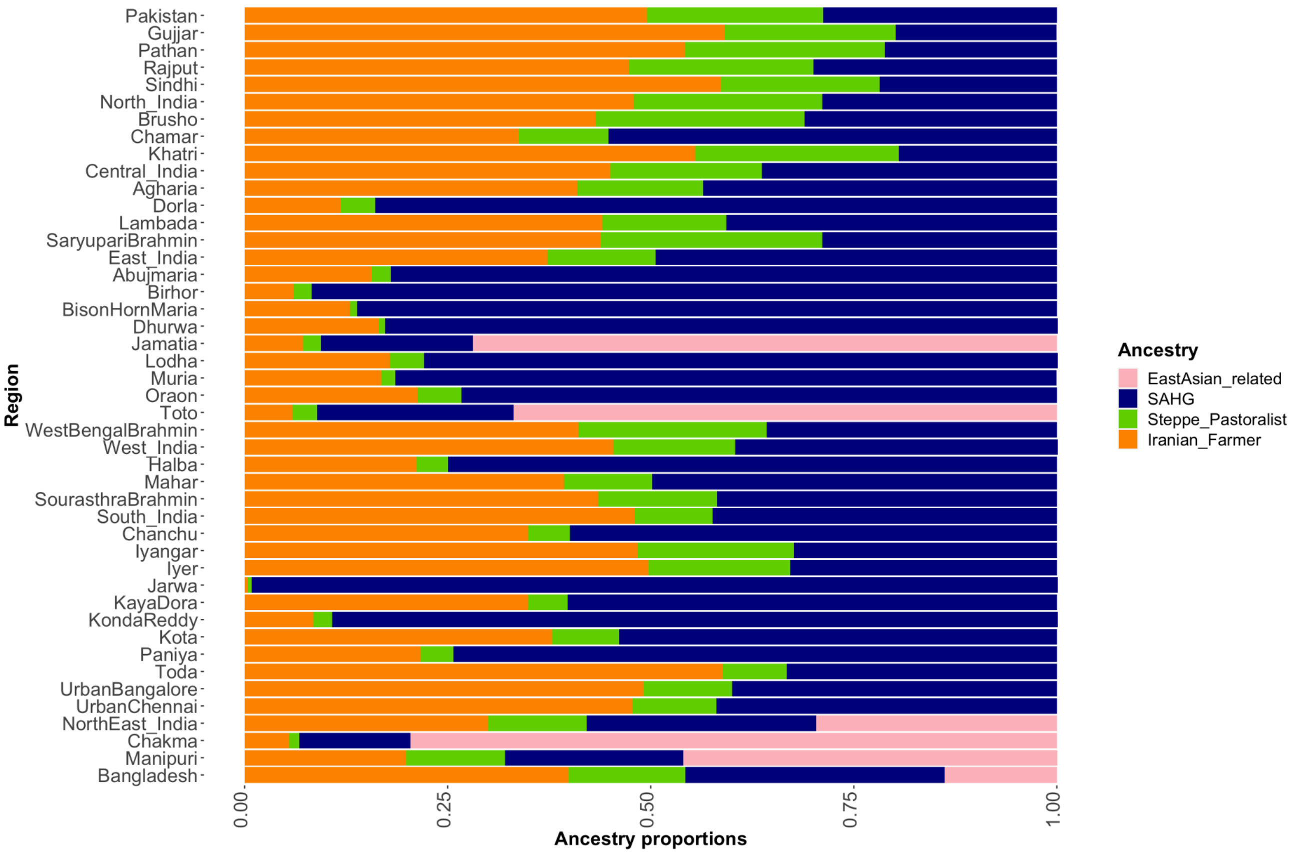

B.

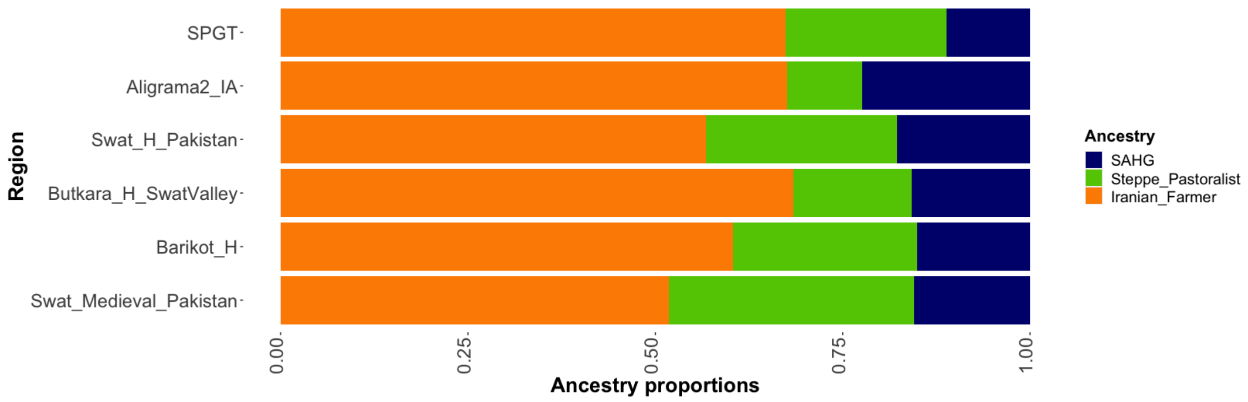


**Figure S1.6 Proportions of Iranian farmer-related, Steppe pastoralist-related, and South Asian hunter-gatherer-related, and East Asian-related ancestries in present day regions of South Asian endogamous groups.** Bars for geographic regions in South Asia and South Asian endogamous groups add to 1, illustrating the ancestry composition. The average proportion of Iranian farmer-related ancestry is shown in orange, Steppe pastoralist-related ancestry in green, SAHG-related ancestry in navy, and East Asian-related ancestry is shown in pink.

A. **Present-day South Asians** Populations are sorted on the y-axis by geography, starting with Pakistan, to North India, Central India, East India, West India, South India, North-East India, and Bangladesh, with endogamous groups listed in their geographic region.
B. **Ancient South Asians** Populations are sorted on the y-axis by time, with the oldest ancient South Asian group at the top (SPGT - ~1491 BCE), to the most recent group at the bottom (*Swat_Medieval_Pakistan* - ~1650 CE). These groups followed the three-way ancestry model (Steppe Pastoralist-related, Iranian Farmer-related, and SAHG-related ancestries).

### Supplementary Note 2. Allele frequency Patterns of LP-Associated Variants in South Asians

#### 2.1 Allele frequency of LP-associated variants

Previous studies have identified several variants associated to LP, including -13.910:C>T and -22.108:G>A in Europeans, -13.915:T>G in Middle Eastern pastoralist groups, and -13.907:C>G, -14.009:T>G, and -14.010:G>C in African pastoralists[16–21]. A study focusing on LP in India identified 7 candidate variants related to regional variation, including -13.779:G>C, -13.801:C>T, -13.879:G>A, -13.915:T>C, -14.011:C>T, -14.012:A>G, -14.026:T>C[22]. We examined the allele frequency of these variants in our South Asian dataset ($n$ = 7,962). We grouped individuals by country– India, Pakistan and Bangladesh. Given the dense sampling in India, we further grouped individuals by region of sampling (e.g., North, Central, East, North-East, West and South). We examined the frequencies separately in endogamous groups with at least 5 individuals in GAsP1. Below we show the distributions of each LP-associated variant in our dataset (note, -13.910:C>T is shown in Figure 2 in the main text).

A.

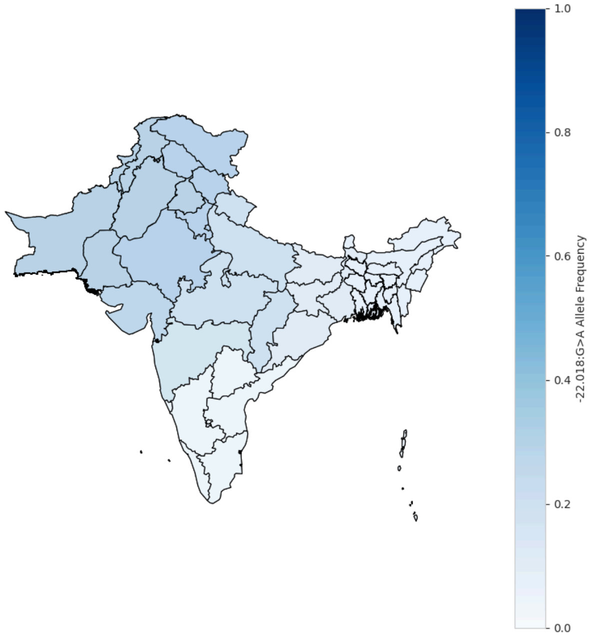


B.

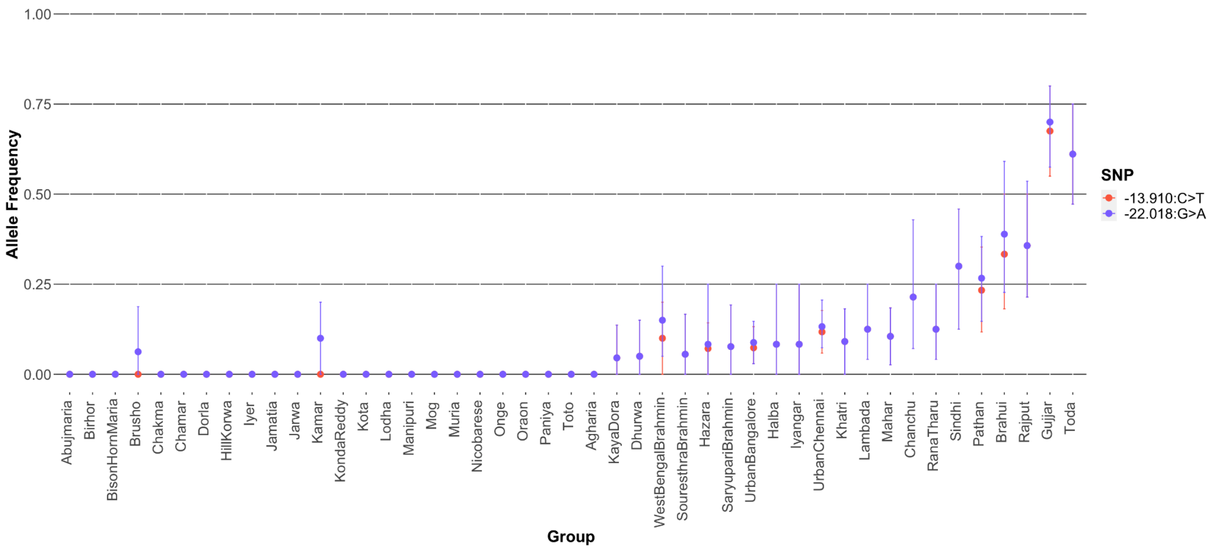


**Figure S2.1 Allele frequencies of European variants -13.910:C>T and -22.018:G>A**

A. **Allele frequencies across South Asia** The allele frequency for -22.018:G>A (blue scale) was calculated for Pakistan, India, and Bangladesh. The allele frequency is highest in Pakistan and North India, and decreases to near zero in South India.

B. **Allele frequency in endogamous groups** The allele frequencies for -13.910:C>T (red) and -22.018:G>A (blue) were calculated for each endogamous group from GAsP1 across India and Pakistan. The allele frequency of each endogamous group increased with geography, that is, groups in South India have low allele frequencies or are zero, and groups in North India and Pakistan have higher allele frequencies. The pastoralist groups Toda (South India) and Gujjar (Pakistan) are notable outliers for both alleles.

A.

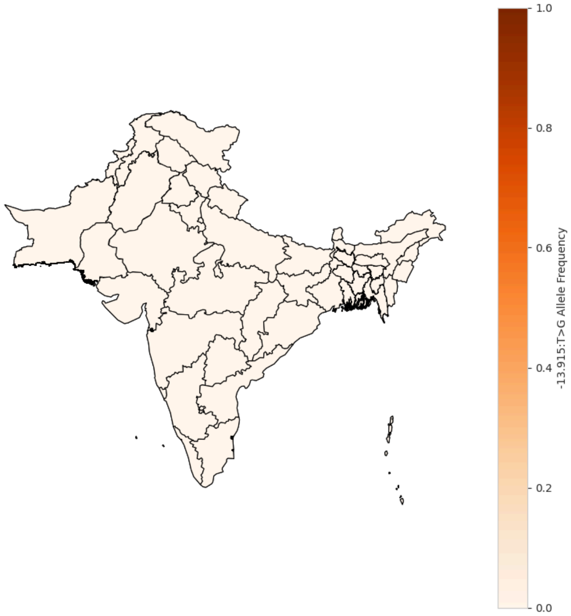


B.

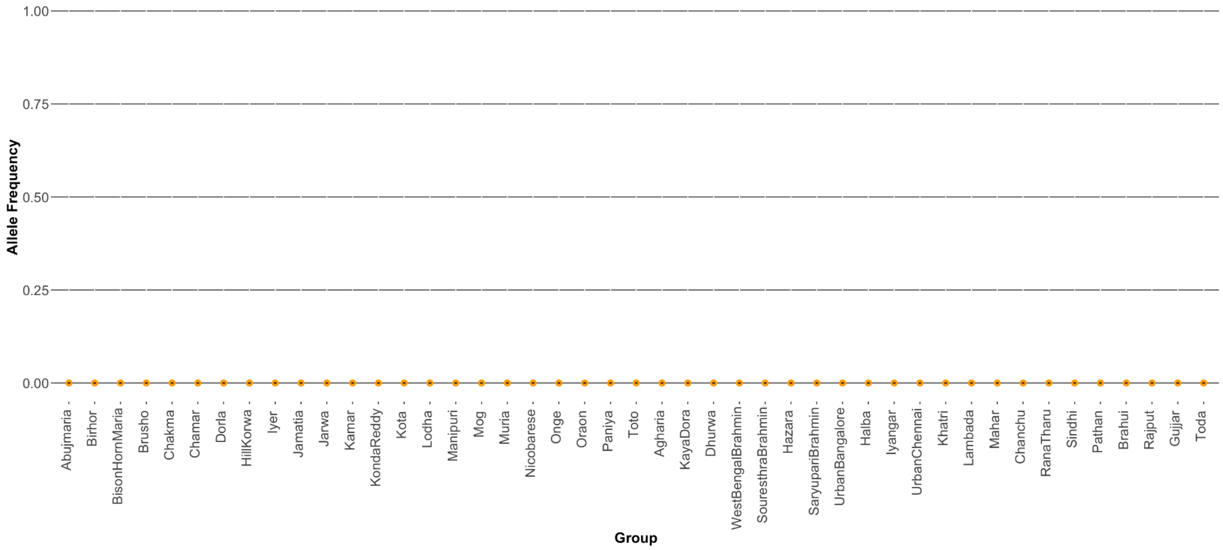


**Figure S2.2 Allele frequencies of Middle Eastern variant - 13.915:T>G**

A. **Allele frequency across South Asia** The allele frequency for -13.915:C>T (orange scale) was calculated for Pakistan, India, and Bangladesh. This allele is present at only 0.14% in South India and 0.06% in Pakistan.
B. **Allele frequency in endogamous groups** The allele frequency for -13.915:C>T (orange scale) was calculated for each endogamous group from GAsP1 across India and Pakistan. This allele is absent across all endogamous groups, including the pastoralist groups Toda and Gujjar.

**A.**

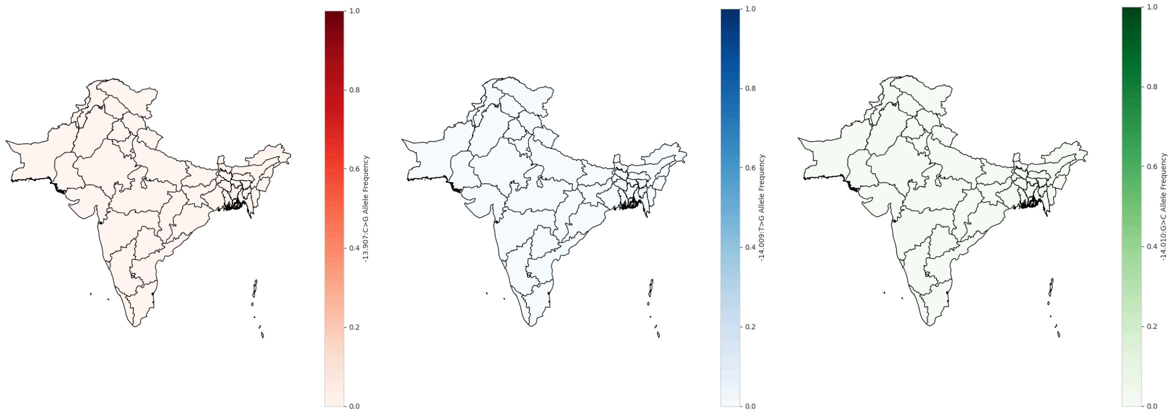


**B.**

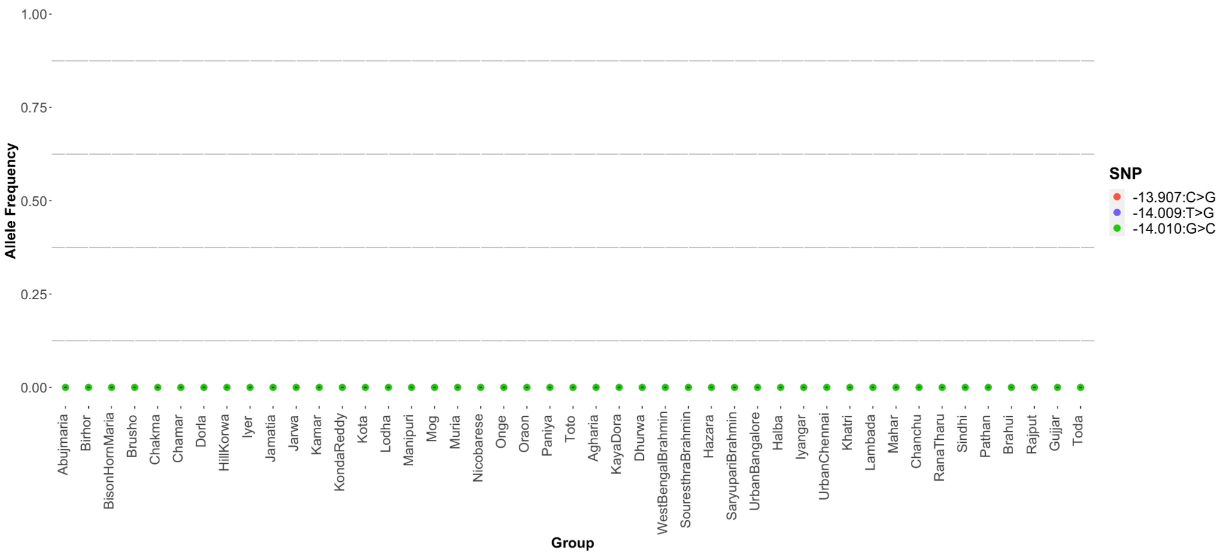


**Figure S2.3 Allele frequencies of African variants -13.907:C>G, -14.009:T>G, -14.010:G>C**

A. **Allele frequencies across South Asia** The allele frequencies for -13.907:C>G (red scale), -14.009:G>A (blue scale), and -14.010:G>C (green scale) were calculated for Pakistan, India, and Bangladesh. The allele frequency for both -13.907:G>C and -14.010:G>C are 0 across South Asia. Only -14.009:G>A is present at 0.05% in Pakistan.
B. **Allele frequency in endogamous groups** The allele frequencies for -13.907:C>G (red scale), -14.009:G>A (blue scale), and -14.010:G>C (green scale) were calculated for each endogamous group from GAsP1 across India and Pakistan. All three alleles were absent in all groups.

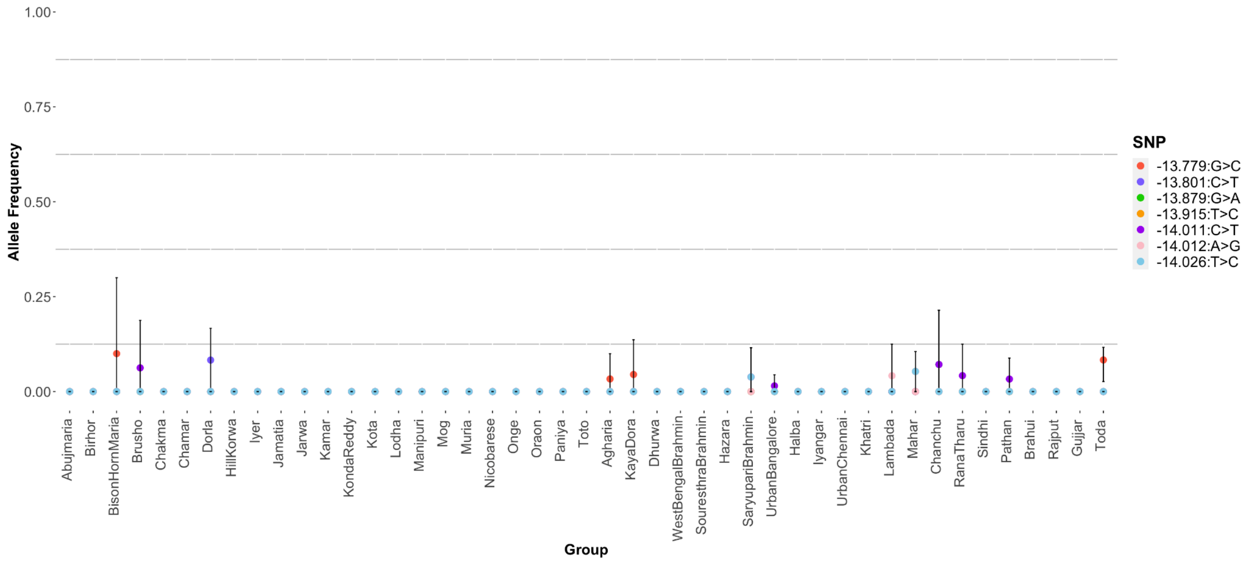


**Figure S2.4 Allele frequencies of candidate LP-associated variants from Gallego Romero et al 2012** The allele frequencies for the 7 putative novel variants identified in [22] were calculated for each of the endogamous groups from GAsP1. No allele shows a clear geographic association.

#### 2.2 Frequency of -13.910:C>T, accounting for dominance

As -13.910:C>T variant confers a dominant phenotype, we also estimated 'LP frequency' by weighting the heterozygous and homozygous genotypes equally (see Methods). Similar to what we observed in allele frequency patterns, we find the highest LP frequency in Pakistan (42.9%) and North India (44.5%), decreasing to South India (6.3%), East India (13.3%), and Bangladesh (11.2%) (Supplementary Table 2.1, Figure S2.5 A). The LP frequency among endogamous groups mirrors the patterns across geographic regions in South Asia, with the lowest LP frequencies (at or near zero) in East, North-East, and South India and high frequencies around 40% in Pakistan (Supplementary Table 2.1, Figure S2.5 B). Strikingly, two pastoralist groups—Toda and Gujjar— exhibit LP frequencies of 88.9% and 90%, respectively (Figure S2.5 B), similar to frequencies observed in Northern Europe[16,17].

Turning to ancient DNA samples from the Swat Valley, Pakistan, we used imputed data and retained only individuals that have at least 3 reads overlapping the -13.910:C>T SNP (Methods). We find that *T allele is absent in Iron or Bronze Ages individuals (*SPGT, n* = 63; *Aligrama2_IA*, *n* = 2), but the LP frequency rises to 18.18% in the historical period (*Swat_H_Pakistan, n* =11), and 50% by the medieval period (*Swat_Medieval_Pakistan, n* = 2) (Figure S2.5 C). This suggests that the LP variants were at low frequency in South Asia prior to the historical period. However, more high-quality ancient DNA data from across time and geography in South Asia will be essential to refine these temporal inferences and better understand the trajectory of LP in the region.

A.

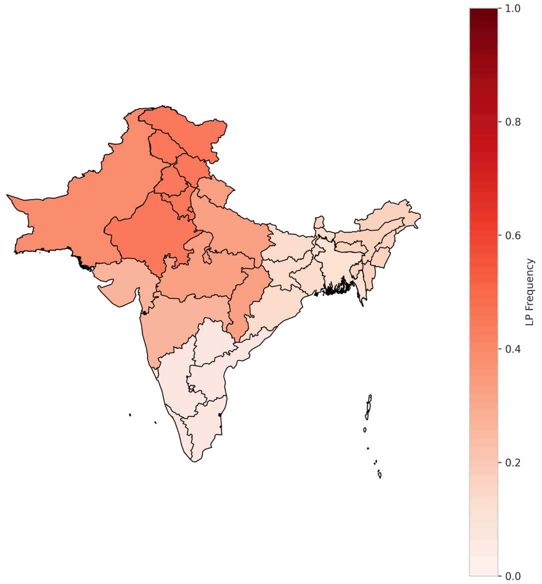

1.0
0.8
0.6
0.4
0.2
0.0
LP Frequency


B.

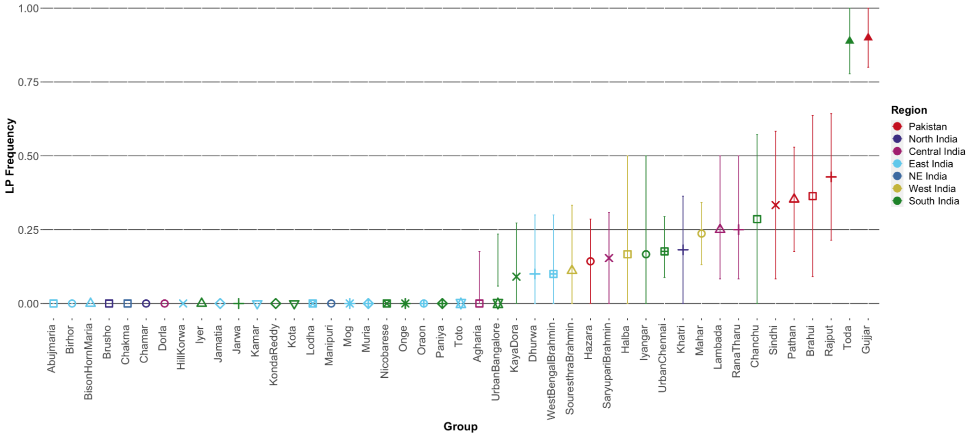

1.00
0.75
0.50
0.25
0.00
LP Frequency
Abujmaria
Birhor
BisonHornMaria
Brusho
Chakma
Chamar
Dorla
HillKorwa
Iyer
Jamatia
Jarwa
Kamar
KondaReddy
Kota
Lodha
Manipuri
Mog
Muria
Nicobarese
Onge
Oraon
Paniya
Toto
Agharia
UrbanBangalore
KayaDora
Dhurwa
WestBengalBrahmin
SouresthraBrahmin
Hazara
SaryuparBrahmin
Halba
Iyangar
UrbanChennai
Khatri
Mahar
Lambada
RanaTharu
Chanchu
Sindhi
Pathan
Brahui
Rajput
Toda
Gujjar
Group
Region
Pakistan
North India
Central India
East India
NE India
West India
South India


C.

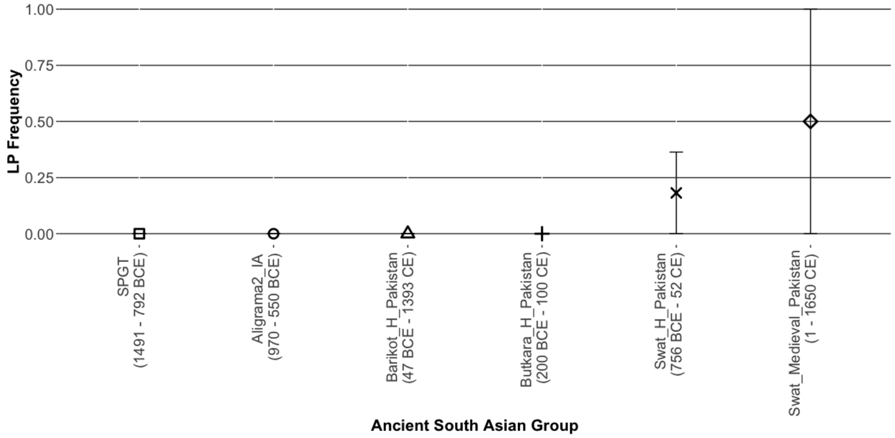

1.00
0.75
0.50
0.25
0.00
LP Frequency
SPGT (1491 - 792 BCE)
Aligrama2_IA (970 - 550 BCE)
Barikot_H_Pakistan (47 BCE - 1393 CE)
Butkara_H_Pakistan (200 BCE - 100 CE)
Swat_H_Pakistan (756 BCE - 52 CE)
Swat_Medieval_Pakistan (1 - 1650 CE)
Ancient South Asian Group

**Figure S2.5 LP frequency across space and time in South Asia**

A. **LP frequency across South Asia.** The LP frequency (red scale) was calculated for Pakistan, India, and Bangladesh. The LP frequency is highest in Pakistan and North India, and decreases in South India, East India, and Bangladesh.

B. **LP frequency in endogamous groups.** The LP frequency was calculated for each endogamous group from GAsP1 across India and Pakistan. The allele frequency of each endogamous group increased with geography, that is, groups in South India have low LP frequencies or are zero, and groups in North India and Pakistan have higher LP frequencies. The pastoralist groups Toda (South India) and Gujjar (Pakistan) are notable outliers. Groups are plotted by color and symbol as in Figure 1A and Figure 1B, representing geographic regions.

C. **LP frequency in ancient South Asians.** The LP frequency was calculated for ancient South Asians from Pakistan (~1491 BCE - 1650 CE) for individuals that had at least 3 reads at -13.910:C>T, considering individuals with at least one -13.910*T allele to be LP. We note LP is absent in samples from the Bronze and Iron Ages, but increases in frequency in the Historic and Medieval periods. The large range of error for Medieval samples can be explained by a small sample size of n=2.

**Distribution of LP-associated variants in 1000 Genomes dataset**

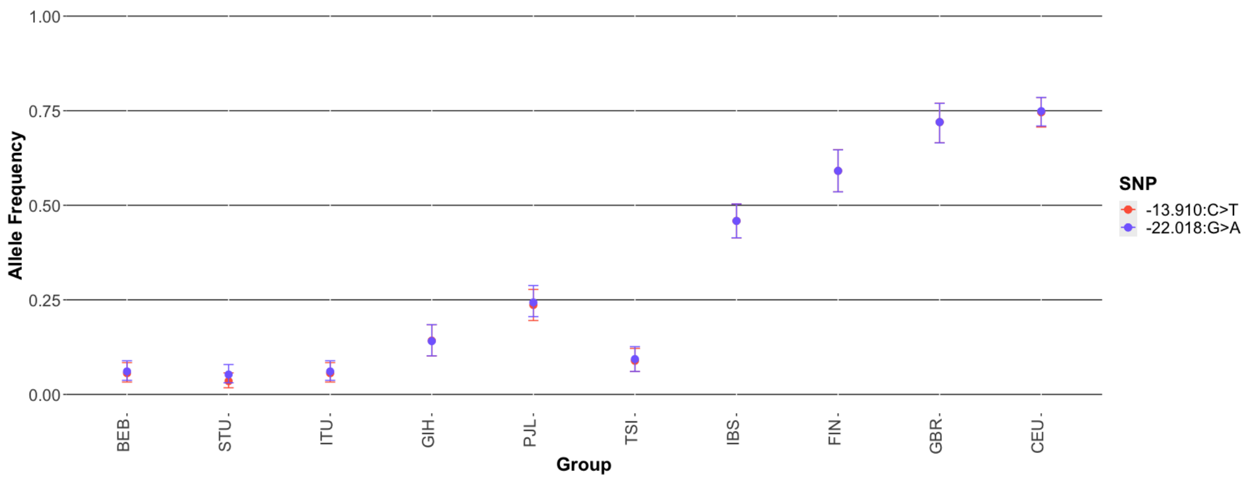


**Figure S2.6 Allele frequencies of European variants -13.910:C>T and -22.018:G>A in 1000G South Asians and Europeans** The allele frequencies for -13.910:C>T (red) and -22.018:G>A (blue) are calculated for South Asian groups (BEB, STU, ITU, GIH, PJL) and European groups (TSI, IBS, FIN, GBR, CEU) from 1000G. In South Asian groups, PJL has the highest allele frequency for both variants, and CEU and GBR have the highest allele frequencies for both variants in Europeans. We observe a geographic gradient, with lowest frequencies to the south and east of each region, and the highest frequencies to the north and west. All other variants that were identified in African, Middle Eastern and South Asians were at frequency of 0 or absent in 1000G groups.

#### 2.3 Linkage Disequilibrium Patterns of LP-Associated Variants in South Asia

Previous studies have found -22.018:G>A to be in complete LD ($r^2 = 1$) with -13.910:C>T in Europeans[16]. We applied PLINK 1.9[23] to calculate the $r^2$ between the -13.910:C>T and -22.018:G>A variants in South Asians. We find the two alleles are in high but incomplete LD ($1 > r^2 >= 0.79$) across India ($r^2 = 0.86$), Pakistan ($r^2 = 0.91$) and Bangladesh ($r^2 = 0.79$) (Supplementary Table 2.2). This trend holds across different regions in India (North $r^2 = 0.91$; Central $r^2 = 0.9$; West $r^2 = 0.91$: East: $r^2 = 0.76$; South $r^2 = 0.65$: North-East $r^2 = 1$). Among endogamous groups (*n* = 45), there are 24 groups where the allele frequency of both variants is 0. In the remaining groups, we find LD ranges between $r^2 = 0.57$-$0.91$ across groups where the variants are in high but not complete LD (n = 7) and are in complete LD in other groups (n = 14) (Supplementary Table 2.2). Interestingly, LD patterns differ across the two pastoralist groups, Gujjar exhibits incomplete LD ($r^2 = 0.89$), but Toda shows complete LD ($r^2 = 1$) across the two variants (Supplementary Table 2.2), which may reflect differences in the IBD patterns among the two groups (Supplementary Note 7). Overall, these results show that unlike in Europeans, the LP associated alleles -13.910:C>T and -22.018:G>A are in high LD in South Asians, though not complete.

To investigate the extent of LD around the -13.910:C>T variant (rs4988235, chr2:135,851,076), we calculated the $r^2$ between the -13.910:C>T variant and every SNP within a 10Mb window for present-day South Asians (*n* = 7,962). Using a threshold of $r^2 > 0.8$, we observe that the LD extends for nearly 1MB from chr2:135,080,336 to chr2:135,950,412 (hg38). Within this block, we see 9 SNPs with $r^2 > 0.8$, including -22.018:G>A (rs182549, chr2:135,859,184; $r^2 = 0.873$) (Figure S2.7, Table S2). We see no SNPs in complete LD ($r^2 = 1$) with -13.910:C>T (Table S2).

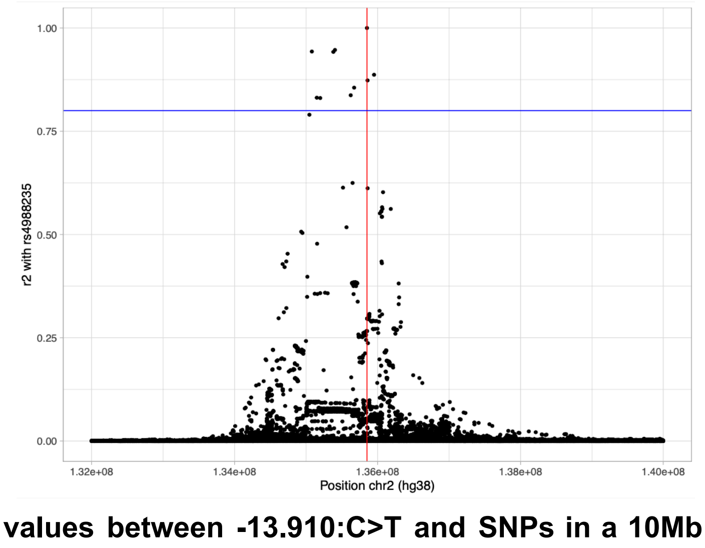


**Figure S2.7 The $r^2$ values between -13.910:C>T and SNPs in a 10Mb window.** The $r^2$ was calculated using plink for SNPs in a 10Mb window around the LP-associated variant -13.910:C>T. The $r^2$ values are plotted on the y-axis against the SNP position on chromosome 2 on the x-axis. The position of -13.910:C>T is indicated by the vertical red line. The horizontal blue line at $r^2 = 0.8$ indicates the $r^2$ threshold implemented to determine if two SNPs are in LD -

any SNP with $r^2$ >= 0.8 is considered to be in LD with -13.910:C>T. A total of 9 SNPs pass this threshold, spanning 0.87 Mb.

**Table S2 The $r^2$ values between -13.910:C>T and 9 neighboring SNPs located within a 10Mb window.** The ID for 9 SNPs in the 1Mb LD block around -13.910:C>T with $r^2$ > 0.8 are listed in column 1. Column 2 lists the positions of these SNPs on Chromosome 2. The $r^2$ values of the SNPs with -13.910:C>T are listed in column 3, with none of the SNPs indicating complete LD.

| **SNP ID** | **Chr 2 Position (hg39)** | **$r^2$ with -13.910:C>T** |
|---|---|---|
| rs7570971 | 135,080,336 | 0.943 |
| rs6730157 | 135,149,518 | 0.831 |
| rs1375131 | 135,197,227 | 0.831 |
| rs3940549 | 135,381,057 | 0.942 |
| rs6709525 | 135,403,522 | 0.946 |
| rs12465802 | 135,623,778 | 0.837 |
| rs62168795 | 135,671,796 | 0.856 |
| rs182549 | 135,859,184 | 0.873 |
| rs6754311 | 135,950,412 | 0.887 |

### Supplementary Note 3. Steppe origin of -13.910*T in South Asia

To investigate the origins of lactase persistence in South Asia, we compared haplotypes in present-day South Asians to those in ancestral reference groups—SAHG, Iranian-farmer and Steppe pastoralist-related populations. We focused on two genomic regions: (i) the *core* region (chr2:135,787,850–135,876,443 in hg38) that spans 88 Kb and includes the *LCT* and *MCM6* genes (annotated based on RefSeq database[24]); and (ii) the *extended* region (chr2:135,080,336-135,950,412 in hg38 that encompasses 870 kb and includes SNPs that are in high linkage disequilibrium (LD, $r^2 > 0.8$) with -13.910:C>T in present-day South Asians (Figure S2.7). For comparative analyses, we grouped individuals by endogamous group or geographic region. We further stratified individuals based on the presence or absence of the derived allele at -13.910:C>T. For this analysis, we used the following subsets of SNPs detailed in Table S3.1.

**Table S3.1. Description of datasets used in the analyses for this section.** This includes the number of variants in different genomic regions.

| Dataset | Data type | Number of variants in core region | Number of variants in extended region |
|---|---|---|---|
| GAsP1 + GAsP2 + LASI-DAD | WGS† | 394 | 3,178 |
| GAsP1 + GAsP2 + LASI-DAD + AADR | WGS† and Imputation | 177 | 1,273 |

†The WGS were all phased with SHAPEIT4[25] using HGDP[3] high-coverage data as reference panels and the HapMap recombination map[26].

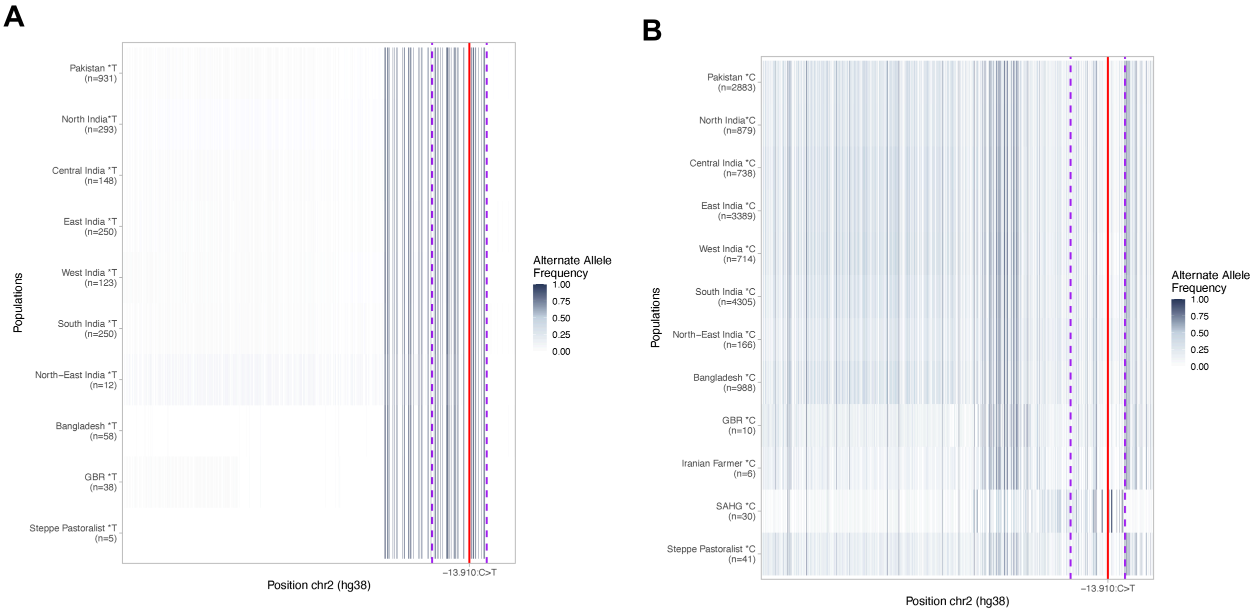


**Figure S3.1. Distribution of the frequency of the alternate allele in the extended region among (A) -13.910*T and (B) -13.910*C haplotypes.** Data are shown for GAsP1, GAsP2, and LASI-DAD individuals grouped by region and country of origin, as well as for GBR individuals and ancestral population proxies representing Steppe pastoralist-related, Iranian farmer-related, and SAHG-related groups. The red line indicates the position of the -13.910:C>T variant, and the dashed purple lines mark the boundaries of the core region.

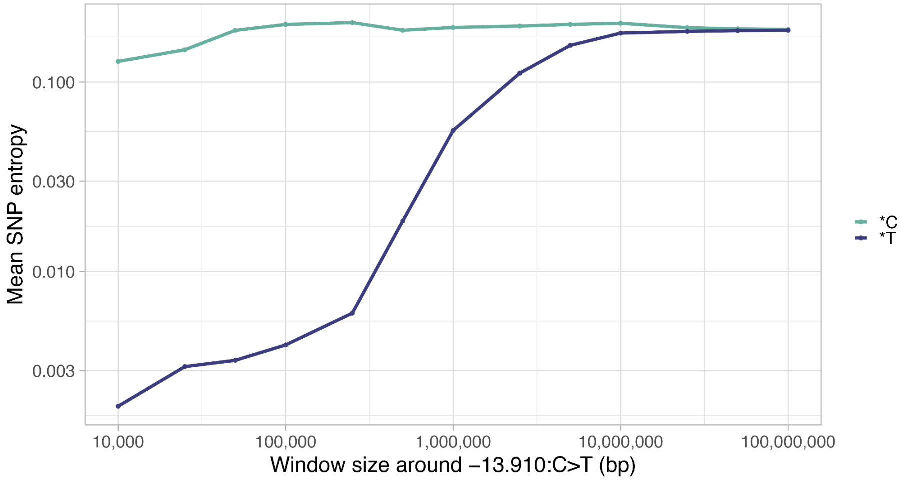


**Figure S3.2. Mean SNP entropy in increasing genomic distance from the -13.910:C>T variant.** Entropy was calculated for reference (-13.910*C; teall) and derived (-13.910*T; blue) haplotypes in windows ranging from 10 kb to 100 Mb, centered on the focal variant (−5 kb/+5 kb to −50 Mb/+50 Mb). The x-axis shows window size (bp, log scale), and the y-axis shows the mean SNP entropy across all SNPs within each window.

Then, we compared how diversity changes as the distance increases from the -13.910:C>T variant. To this end, we measured the entropy in windows of increasing distance from the focal variant. As shown earlier, there is markedly reduced diversity for -13.910*T compared to -13.910*C haplotypes (entropy = 0.002 vs 0.129 within the 10kb window). Interestingly, this depletion recovers slowly with increasing distance, exhibiting comparable diversity levels to -13.910*C haplotypes at approximately 5 Mb away from -13.910:C>T (Fig S3.2).

#### 3.1 Core Haplotype

We focused on the core region that spans 88 Kb and includes the *LCT* and *MCM6* genes and stratified individuals based on the presence or absence of the derived allele at -13.910:C>T. We find the present-day South Asian haplotypes harboring the -13.910*T allele (South Asian*T, $n$ = 2,065) have markedly lower diversity than the -13.910*C haplotypes (South Asian*C, $n$ = 13,859), including a higher proportion of fixed variants (68.3% vs. 0.7%; p-value < 2.2e-16), depletion of SNPs at intermediate (25–75%) allele frequencies (0% vs. 21.8%; p-value < 2.2e-16) and lower entropy (0.005 vs. 0.2, p-value < 2.2e-16) (Figure S3.1-2). Next, we compared the haplotypes in present-day South Asians with the ancestral reference populations, by counting the number of allelic differences between each test and reference haplotypes. We report the distribution separately for South Asian*T haplotypes (Figure S3.3) and for South Asian*C haplotypes (Figure S3.4). We also compared South Asian haplotypes to present-day Europeans, GBR (Table S3.5A-B).

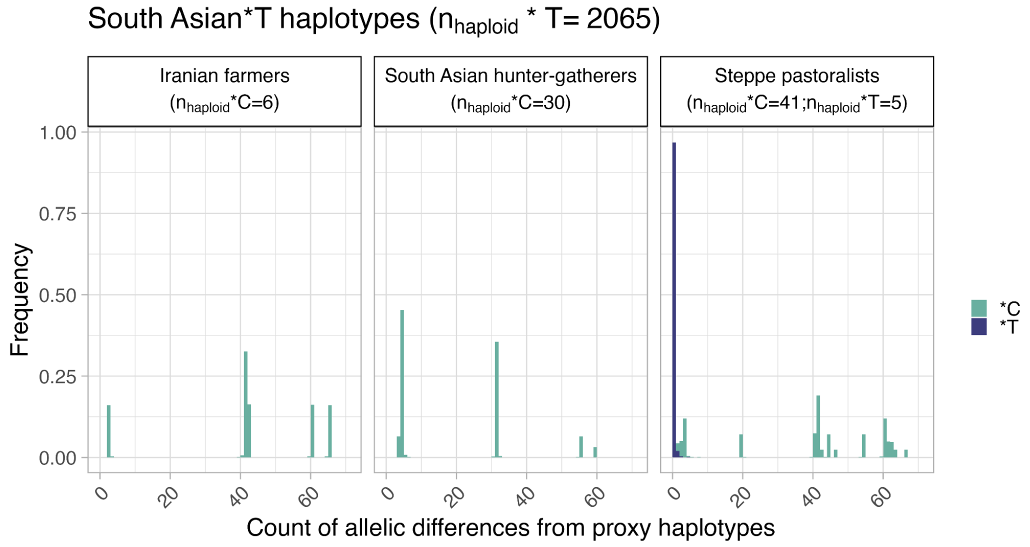


**Figure S3.3. Distribution of allelic differences between present-day South Asian*T haplotypes and ancestral sources.** The x-axis represents the number of allelic differences between South Asian*T haplotypes and an ancestral source haplotype. Ancestral source haplotypes carrying the T allele are shown in blue, and those carrying the C allele are shown in teal. For each pairwise comparison (e.g., South Asian*T vs. Steppe pastoralist*T, South Asian*T vs. Steppe pastoralist*C, etc.), frequencies are normalized so that the values sum to 1 within each group, allowing for direct comparison of distribution patterns across different ancestral sources. We note that -13.910*T allele is absent in Iranian farmers and SAHG-related populations.

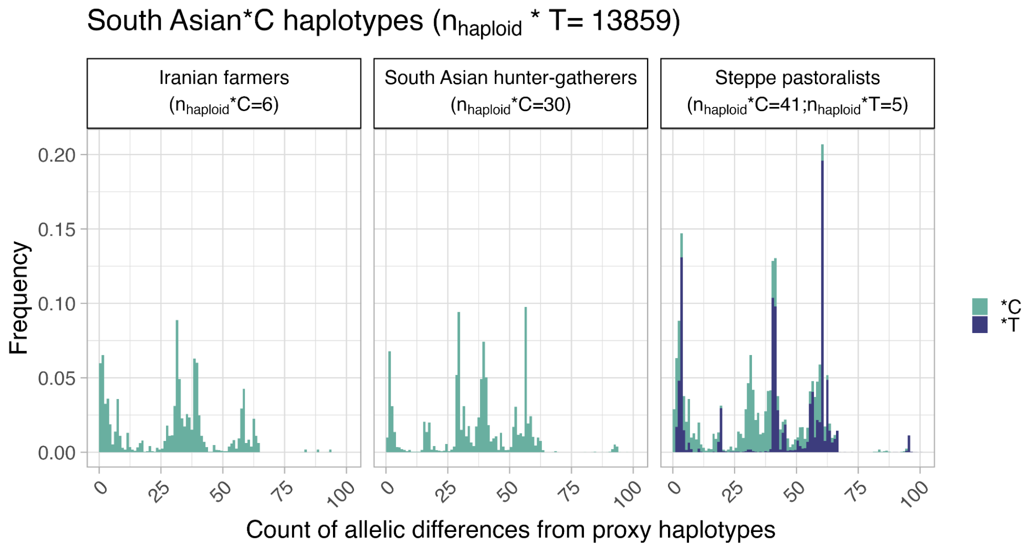


**Figure S3.4 Distribution of allelic differences between present-day South Asian*C haplotypes and ancestral sources.** The x-axis represents the number of allelic differences between South Asian*C haplotypes and an ancestral source haplotype. Ancestral source haplotypes carrying the T allele are shown in blue, and those carrying the C allele are shown in teal. For each pairwise comparison (e.g., South Asian*C vs. Steppe pastoralist*T, South Asian*C vs. Steppe pastoralist*C, etc.), frequencies are normalized so that the values sum to 1 within each group, allowing for direct comparison of distribution patterns across different ancestral sources.

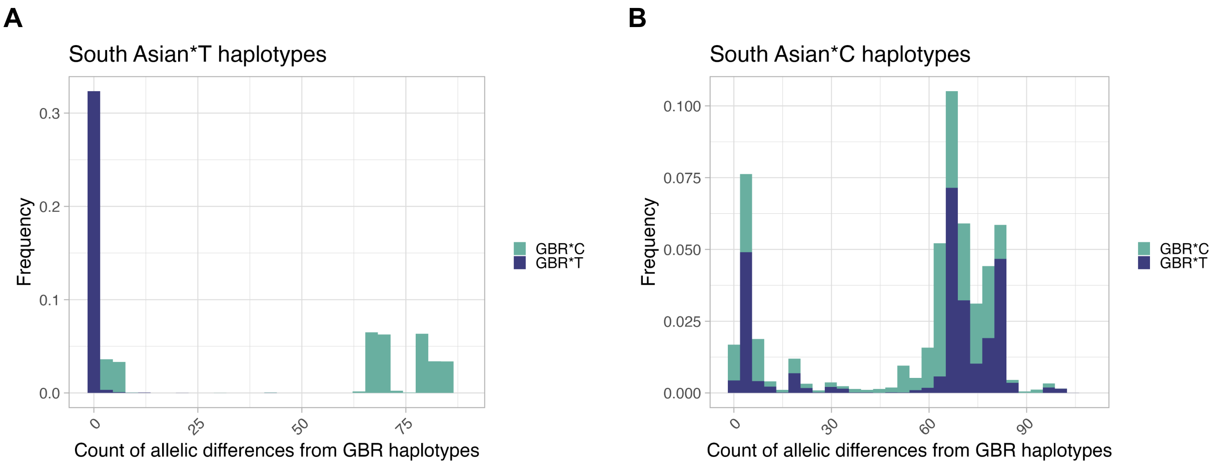

**Figure S3.5 Distribution of allelic differences between present-day South Asian haplotypes and GBR haplotypes.** Panel (A) shows the number of allelic differences between South Asian*T haplotypes and GBR*T (in blue) or GBR*C haplotypes (in teal), while panel (B) shows the same for South Asian*C haplotypes. For each comparison (e.g., South Asian*T vs. GBR*T, South Asian*T vs. GBR*C, etc.), frequencies are normalized so that the total counts sum to 1 within each pairwise comparison group, enabling direct comparison of the distribution shapes across different haplotype combinations.

#### *Haplostrips*

We visualized the allele sharing patterns in the core region using *Haplostrips*[27]. For North India, Central India, West India, East India, and South India, we randomly selected 10 individuals homozygous for the C allele (C|C) and 10 individuals homozygous for the T allele (T|T) at the −13.910:C>T variant. We also included 10 randomly selected GBR individuals for comparison. We ran *Haplostrips* with the option -c 0.00 to retain all variant sites in each population (Figure S3.6). Consistent with our pairwise comparisons (Fig S3.3-4), we find all the 141 haplotypes carrying -13.910*T alleles are identical (Figure S3.6A-S3.7A). In contrast, -13.910*C haplotypes did not show a clear clustering pattern, with substantially larger variation across haplotypes and only a small set (max 18 of 100 haplotypes) that are identical to each other (Figure S3.6B-S3.7B). The average number of difference between the selected South Asian*C haplotypes and GBR*T haplotypes is 40.94, comparable to results of the pairwise comparisons (Figure S3.4).

A

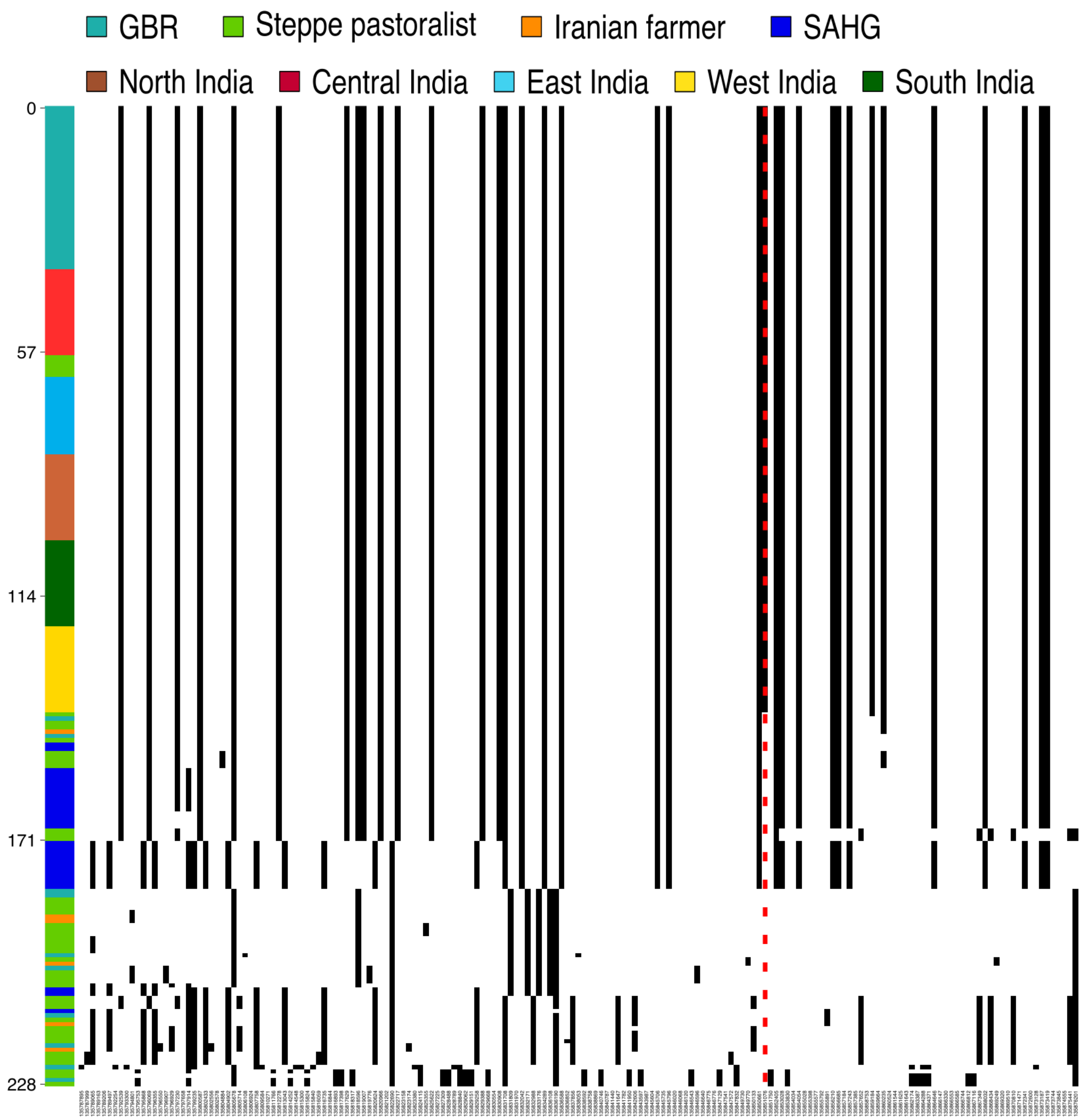
GBR
Steppe pastoralist
Iranian farmer
SAHG
North India
Central India
East India
West India
South India
0
57
114
171
228

**B**

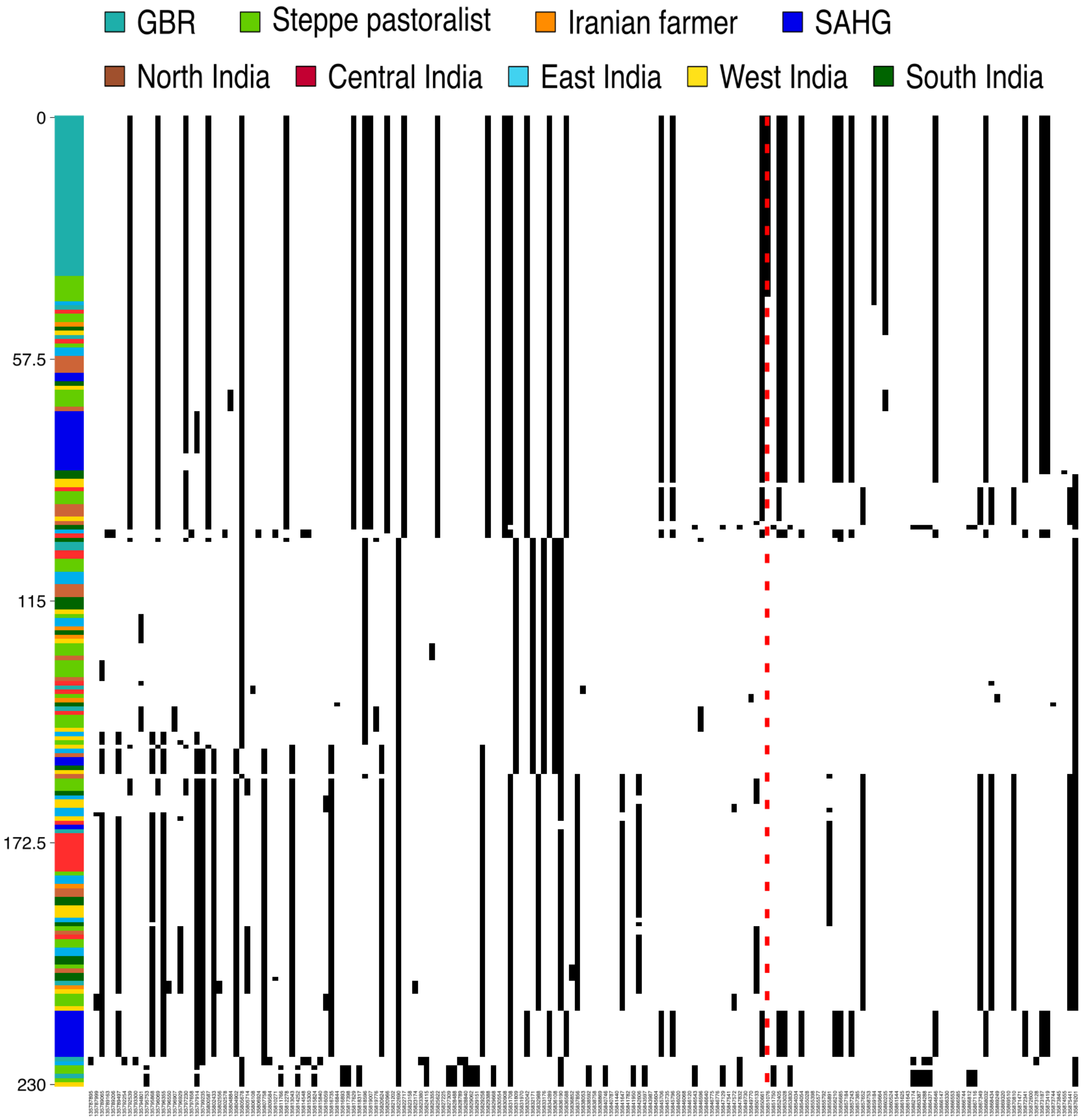


**Figure S3.6.** *Haplostrips* plot of the core region showing allele-sharing patterns for individuals from Pakistan, North India, Central India, West India, East India, and South India with (T|T) genotypes (A) and (C|C) genotypes (B), alongside GBR and Onge from our modern datasets, and individuals from the genetic groupings *Central_Steppe_MLBA* and *Indus_Periphery_Pool*, as well as from the city Sarazm in AADR. Only individuals with reads overlapping the −13,910:C>T variant are shown. The default color scheme was used, with the reference allele (REF) in white and the alternate allele (ALT) in black. The red dashed line marks the position of the −13,910:C>T variant (chr2:135,851,076).

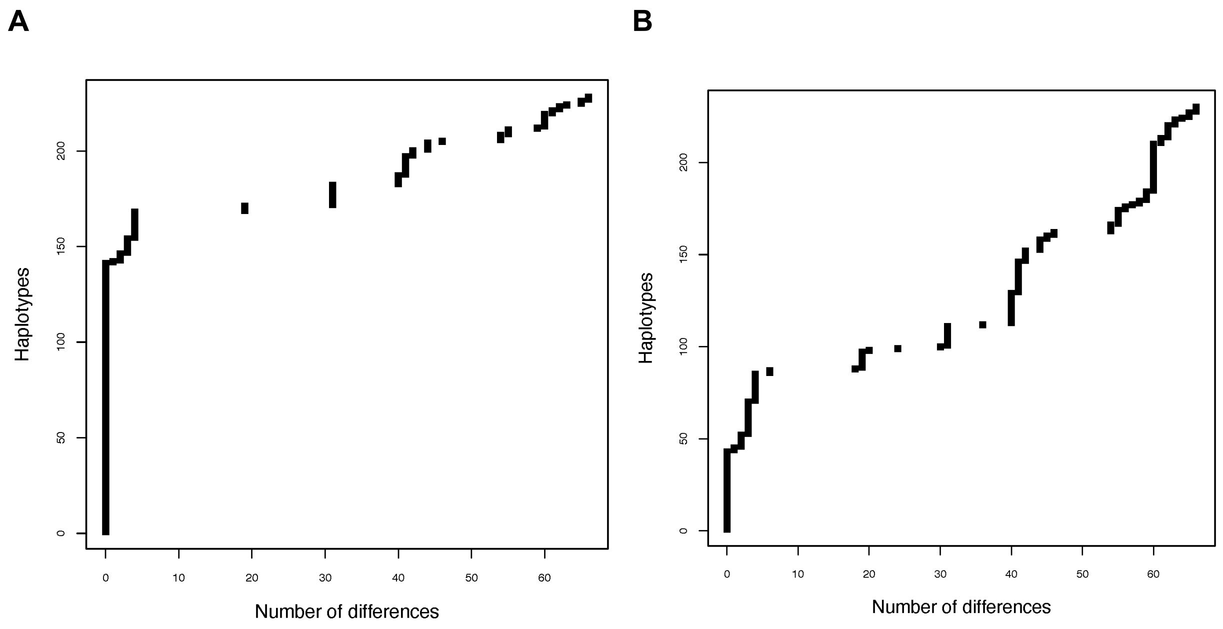


**Figure S3.7.** Number of differences between -13.910:C>T core haplotypes in individuals from Pakistan, North India, Central India, West India, East India, South India with (T|T) genotypes (A) and (C|C) genotypes (B), alongside GBR and Onge from our modern datasets and individuals from the genetic grouping *Central_Steppe_MLBA* and *Indus_Periphery_Pool*, and from the city of Sarazm from AADR.

#### 3.2 Extended Haplotype

For the extended region (870 kb around -13.910:C>T), we observed similar diversity patterns as the core region. Specifically, the -13.910*T haplotypes show similarly reduced diversity, including a higher proportion of fixed variants (56.9% vs. 0.6%; p-value < 2.2e-16), depletion of SNPs at intermediate (25–75%) allele frequencies (0% vs. 19.4%; p-value < 2.2e-16) and lower entropy (0.013 vs. 0.181, p < 2.2e-16). Compared to the ancestral reference populations, the South Asian*T haplotypes differs from the Steppe pastoralist*T haplotypes by an average of 6.6 variants (range: 0–557), compared to 236.2 differences with Steppe pastoralist*C, 227.3 with Iranian farmer*C, and 236.2 with SAHG*C. In contrast, South Asian*C haplotypes show much higher overall divergence: 294.9 (Steppe pastoralist*T), 302.5 (Steppe pastoralist*C), 283.0 (Iranian farmer*C), and 283.5 (SAHG*C). Together, these results point to a Steppe origin of the -13.910*T allele in South Asia, similar to Europeans[28].

**A**

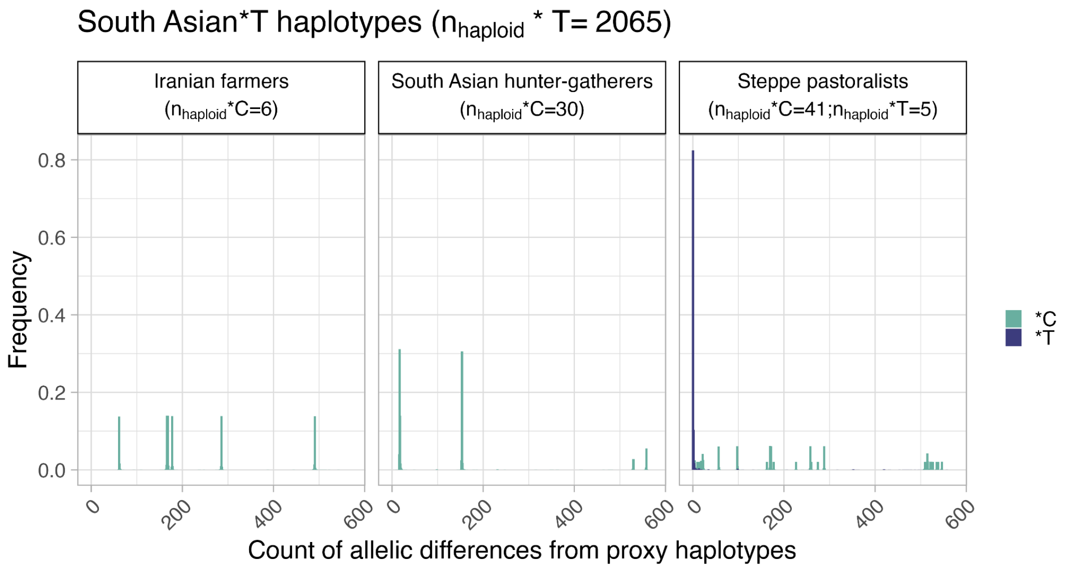


**B**

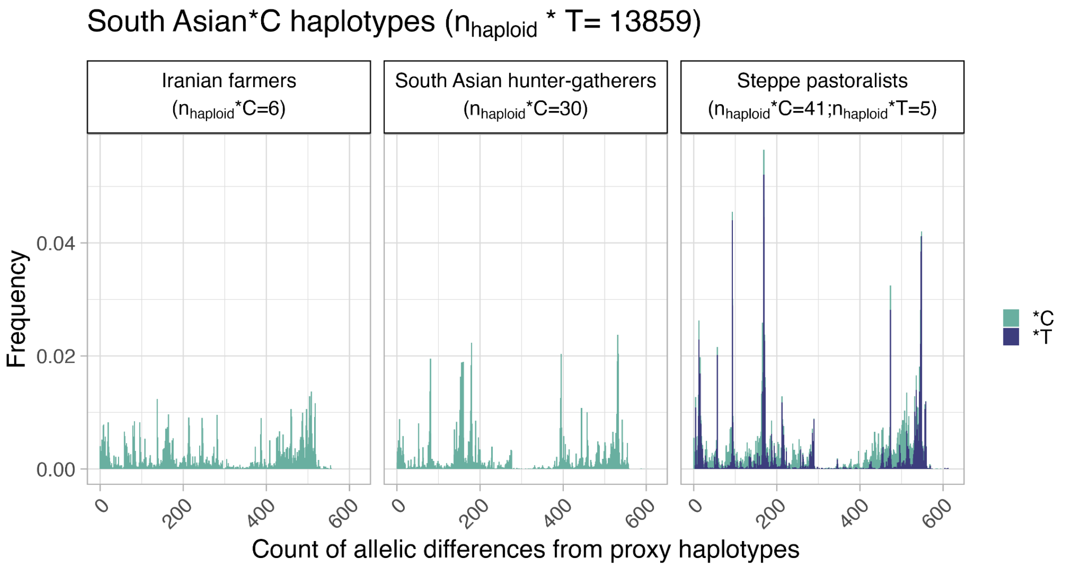


**Figure S3.8. Distribution of allelic differences between present-day South Asians and ancestral populations in the extended region. A.**-13.910*T haplotypes, **B.** -13.910*C haplotypes. The x-axis represents the number of allelic differences between South Asian*T haplotypes and each of the three ancestral source haplotype. Ancestral source haplotypes carrying the T allele are shown in blue, and those carrying the C allele are shown in teal. For each pairwise comparison (e.g., South Asian*T vs. Steppe pastoralist*T, South Asian*T vs. Steppe pastoralist*C, etc.), frequencies are normalized so that the values sum to 1 within each group, allowing for direct comparison of distribution patterns across different ancestral sources.

Finally, we compared the observed -13.910*T haplotype length (extended region of 870 kb based on LD patterns ($r^2$ > 0.8) in the *LCT* region (Figure S2.7)) to the theoretical expectation for Steppe gene flow in South Asians that occurred around 3,500 years ago or 125 generations ago[4]. Using the SMC' model[29], admixture tract lengths can be modeled as an exponential distribution as $Exp[2N(1-m)(1-e^{-T/2N})]$ where $N$ = population size, $m$ = admixture proportion and $T$ = timing of admixture in generations. We assume $N$ = 10,000, $T$ = 125 generations and $m$ ranges between [0.05, 0.1, 0.2, 0.3, 0.4] to mimic parameters in real data[4] (Supplementary Table S1.2). To convert the genetic distance (in Morgans) to physical distance (in base pairs), we used the deCODE pedigree-based genetic map[30] which corresponds to a recombination rate of 0.58 cM/Mb in the *LCT* region. We find that, for a range of admixture proportions tested, the observed length of -13.910*T haplotype (extended region) appears to be consistent with the expected admixture tract length for Steppe pastoralist gene flow in South Asia (Fig S3.9).

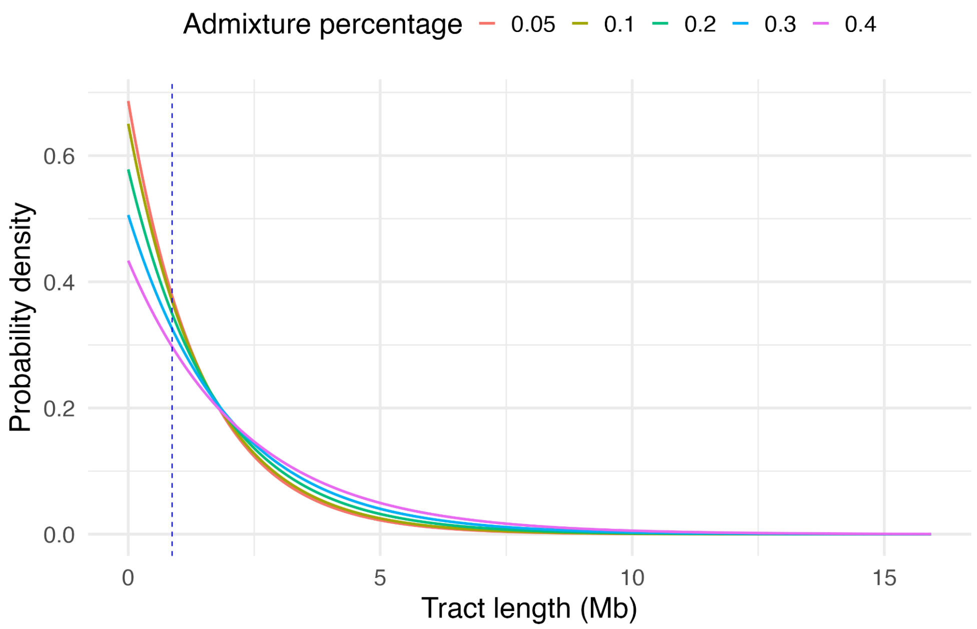


**Figure S3.9. Expected Steppe pastoralist haplotype lengths in South Asians.** Distribution for varying admixture percentage, computed using the SMC' estimator, with a population size ($N$) =10,000, a time of admixture ($T$) of 125 generations and a recombination rate of 0.58 cM/Mb. The dashed vertical line represents the length of the extended haplotype of 870 kb.

### Supplementary Note 4. Local Ancestry Inference

#### 4.1 Method

We developed a local ancestry inference method to identify the ancestral origins of the core haplotypes. The approach is based on quantifying haplotype similarity between present-day South Asian haplotypes and reference haplotypes from ancestral populations, under the assumption that haplotypes with fewer differences are more likely to share common ancestry. For each phased South Asian haplotype, we computed the number of SNP mismatches relative to each reference haplotype ($Nb_{diff}$) and transformed this into a similarity score that declines steeply with increasing differences:

$$Score = \frac{1}{1+10^{*}Nb_{diff}}.$$

The average similarity score across individuals within each ancestral population ($S_{anc}$) was then computed and normalized across all reference populations to obtain ancestry probabilities ($P_{anc}$). Local ancestry was assigned to the ancestral population with the highest normalized probability exceeding a confidence threshold of 0.8, ensuring only high-confidence assignments. This method was applied to haplotypes carrying the -13.910*T allele to evaluate their ancestral origins. The resulting probability distributions are summarized in Figure S3.9.

Among the 2,065 South Asian*T haplotypes, we assigned Steppe pastoralist–related ancestry to 1,998. The rest of the 67 haplotypes were not assigned to any ancestry, as the probability for all three ancestral populations was less than 0.8. We refer to the -13.910*T present-day South Asians haplotypes inferred to have Steppe pastoralist–related at this locus as 'local Steppe*T' haplotypes.

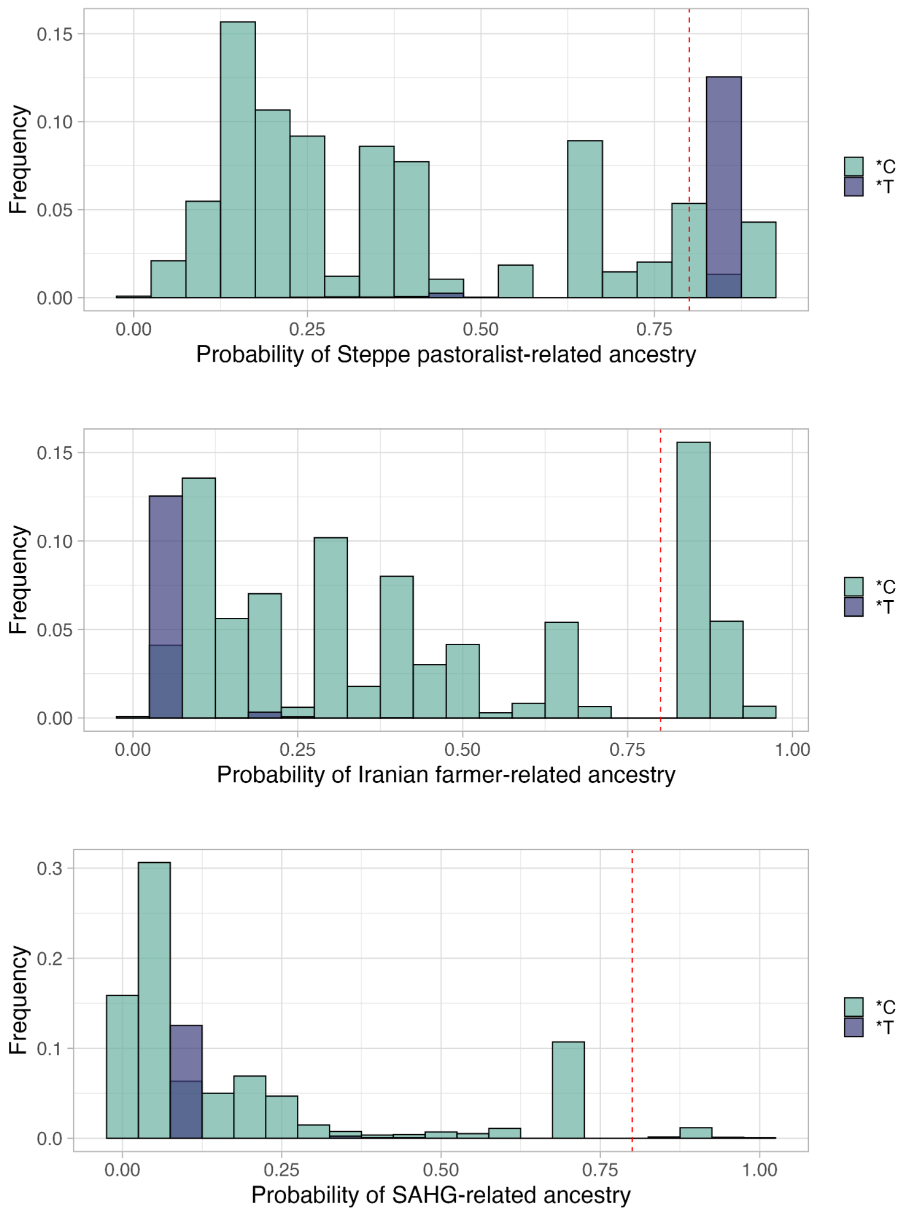


**Figure S4.1. Probability of ancestry for each of the South Asian core haplotypes.** Colored by the -13.910:C>T allele (T in teal and C in green), shown for each of the three ancestries: Steppe pastoralist-, Iranian farmer-, and SAHG-related. The red dotted line represents our threshold of 0.8.

#### 4.2 Relationship between local ancestry and -13.910*T frequency

We compared the local ancestry in the core region (local Steppe*T frequencies) to the allele frequency of -13.910*T (Table S4.1). We grouped individuals by geographic region or country. We find strong correlation between the haplotype frequencies of local Steppe*T and the allele frequency of -13.910*T (Pearson's r = 0.999, p-value = 5.97 × $10^{-7}$, Figure S4.7). We performed a similar analysis focusing on the endogamous communities. For each group with more than 10 individuals, we compared the inferred frequency of local Steppe*T haplotypes with the -13.910*T allele frequency. The two measures were also highly correlated across groups (Pearson's r = 0.998, p-value = 5.657e-12) (Table S4.2, Figure S4.2). In concordance, the chi-square tests of homogeneity showed no significant deviation between local Steppe*T and -13.910*T frequencies across geographic regions or endogamous groups (p-value > 0.05 for all comparisons) (Fig S4.7).

To formally test for deviations, we compute a *Z*-score = $(\mu_{local} - \mu_{AF})/\sigma$, where $\mu_{local}$ is the average local Steppe*T frequency and $\mu_{AF}$ is the average -13.910*T allele frequency; and the standard error is estimated as ($\sigma = \sqrt{\sigma^2_{local} + \sigma^2_{AF}}$). To account for the sampling noise, $\sigma$ for local Steppe*T and -13.910*T allele frequency was estimated using a binomial model. A *Z*-score > 3 implies significant deviation from neutral expectation. Across all regions and endogamous groups analyzed, we found no significant deviation between -13.910*T frequency and the local Steppe*T ancestry proportions (all |Z| < 3; Tables S4.1–S4.2). These results indicate that the observed distribution of the –13.910*T allele is statistically consistent with the Steppe pastoralist ancestry proportion in both regional and community-level analyses.

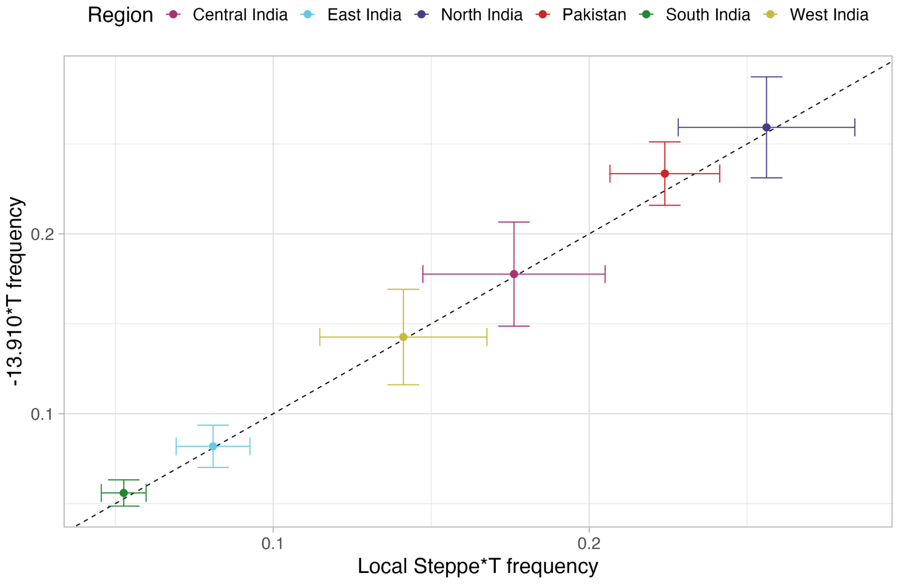


**Figure S4.2 Relationship between local Steppe*T haplotype frequency and -13.910*T allele frequency across regions.** Points are colored by region. The dashed black line represents equality between the two frequencies (x = y). Error bars represent +/-2 standard errors of the mean frequency, calculated assuming a binomial distribution.

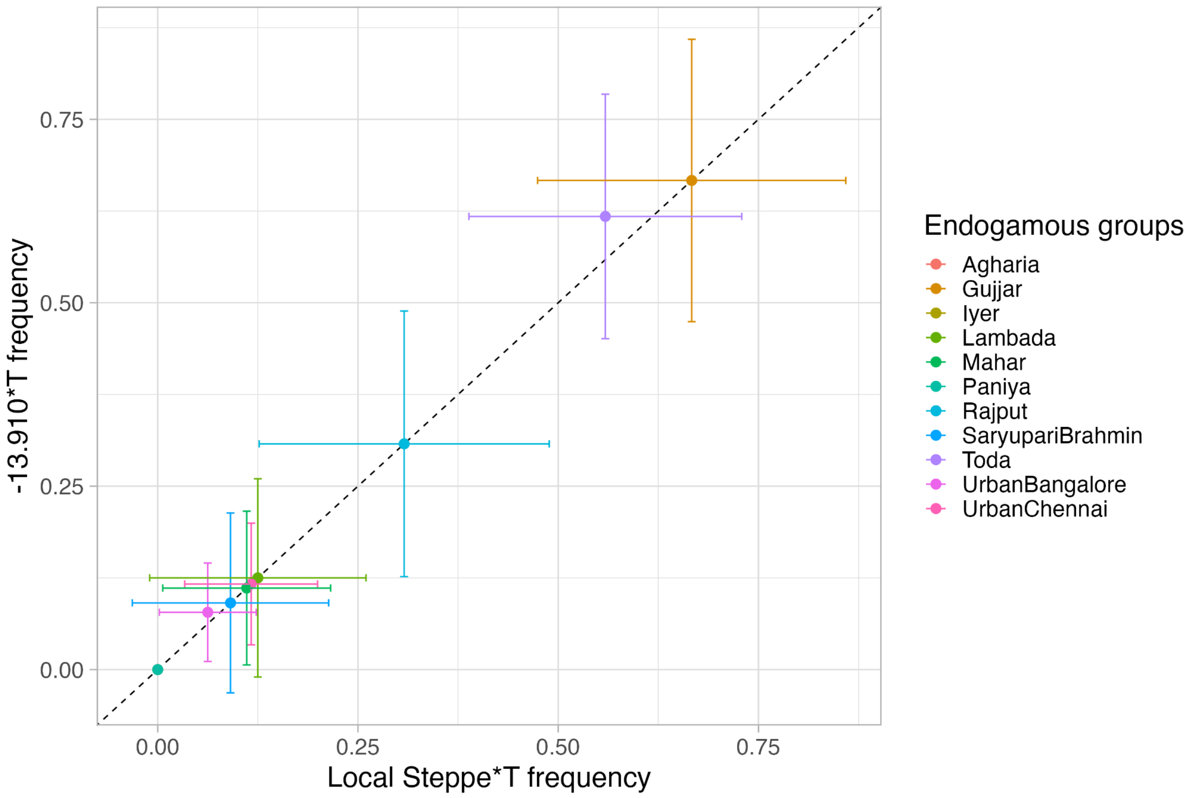


**Figure S4.3 Relationship between local Steppe*T haplotype frequency and -13.910*T allele frequency across endogamous groups.** Points are colored by group. The dashed black line represents equality between the two frequencies (x = y). Error bars represent +/-2 standard errors of the mean frequency, calculated assuming a binomial distribution.

#### 4.3 Local Ancestry Deviation test

Focusing on the individuals on the cline, that fit the three way model ($n$ = 4,946), we applied the Local Ancestry Deviation (LAD) test[31] that measures deviations in local ancestry proportion from the genome-wide ancestry proportion. The LAD test is similar to a Z-score and was computed as follows:

$$Z = (\mu_{local} - \mu_{global})/\sigma_{global}$$

where $\mu_{local}$ is the inferred local Steppe*T haplotype proportion, $\mu_{global}$ is the mean genome-wide Steppe ancestry proportion per individual (based on *qpAdm*), and $\sigma_{global}$ is the corresponding standard deviation. Because the endogamous groups are highly homogeneous, the empirical distribution of genome-wide ancestry proportions does not adequately capture uncertainty and thus, we estimated standard errors using a binomial approximation to account for sampling uncertainty. We applied this test to both individuals grouped by region and separately per endogamous group.

For all regions analyzed, we find no significant deviation between the genome-wide and local Steppe*T ancestry proportion (|Z| < 3, Figure 4A, Table S4.1). In most endogamous groups, we find the local Steppe*T frequency is similar to the genome-wide Steppe ancestry proportion (|Z| < 3, Figure 4A). Only two groups, Toda and Gujjar, show a significant deviation from the expectation ($Z$ = 8.74 and 5.54 respectively, Table S54.2). This enrichment suggests that other evolutionary processes (beyond gene flow)— such as strong genetic drift or natural selection— may have contributed to the frequency of -13.910*T in Toda and Gujjar.

**Table S4.1 Local Steppe*T haplotype frequencies, -13.910*T allele frequencies, and genome-wide Steppe ancestry across South Asian regions.** For each region/country, we report the frequency of haplotypes assigned to Steppe ancestry among -13.910*T carriers (local Steppe*T), the overall -13.910*T allele frequency, the genome-wide proportion of Steppe pastoralist–related ancestry and the different z-scores comparing these values.

| **Region** | ***n*** | $\mu_{AF}$ +/- $\sigma_{AF}$ | $\mu_{local}$ +/- $\sigma_{local}$ | $\mu_{global}$ +/- $\sigma_{global}$ | **Z-score** $\mu_{AF}$ vs $\mu_{local}$ | **Z-score** $\mu_{local}$ vs $\mu_{global}$ |
|---|---|---|---|---|---|---|
| North | 500 | 0.263 +/- 0.0139 | 0.26 +/- 0.0140 | 0.232 +/- 0.0746 | 0.153 | 0.378 |
| Pakistan | 1212 | 0.237 +/- 0.0086 | 0.227 +/- 0.00851 | 0.217 +/- 0.0438 | 0.783 | 0.227 |
| Central | 345 | 0.177 +/- 0.0145 | 0.175 +/- 0.0145 | 0.188 +/- 0.0509 | 0.071 | -0.240 |
| West | 348 | 0.145 +/ 0.0134 | 0.144 +/- 0.0133 | 0.148 +/- 0.0414 | 0.076 | -0.105 |
| East | 604 | 0.0869 +/ - 0.0081 | 0.086 +/- 0.0081 | 0.154 +/- 0.0503 | 0.0724 | -1.35 |
| South | 1937 | 0.056 +/ - 0.0037 | 0.0524 +/- 0.00358 | 0.0965 +/- 0.0469 | 0.653 | -0.939 |

**Table S4.2. Local Steppe*T haplotype frequencies, -13.910*T allele frequencies, and genome-wide Steppe ancestry across endogamous groups.** For each endogamous groups, we report the frequency of haplotypes assigned to Steppe ancestry among -13.910*T carriers (local Steppe*T), the overall -13.910*T allele frequency, and the genome-wide proportion of Steppe pastoralist–related ancestry and the different z-scores comparing these values.

| **Endogamous group** | ***n*** | $\mu_{AF}$ +/- $\sigma_{AF}$ | $\mu_{local}$ +/- $\sigma_{local}$ | $\mu_{global}$ +/- $\sigma_{global}$ | **Z-score** $\mu_{AF}$ vs $\mu_{local}$ | **Z-score*** $\mu_{local}$ vs $\mu_{global}$ |
|---|---|---|---|---|---|---|
| Toda | 13 | 0.6185+/- 0.095 | 0.538 +/- 0.098 | 0.078 +/- 0.053 | 0.563 | **8.74*** |
| Gujjar | 13 | 0.654 +/- 0.093 | 0.654 +/- 0.093 | 0.211 +/- 0.080 | 0 | **5.54*** |
| Iyer | 11 | 0.000 +/- 0.000 | 0.000 +/- 0.000 | 0.174 +/- 0.081 | NA | -2.16 |
| Rajput | 13 | 0.308 +/- 0.091 | 0.308 +/- 0.091 | 0.227 +/- 0.082 | 0 | 0.979 |
| Lambada | 12 | 0.125 +/- 0.068 | 0.125 +/- 0.068 | 0.153 +/- 0.073 | 0 | -0.377 |
| Urban Chennai | 30 | 0.117 +/- 0.041 | 0.117 +/- 0.041 | 0.103 +/- 0.039 | 0 | 0.346 |
| Mahar | 17 | 0.118 +/- 0.055 | 0.118 +/- 0.055 | 0.112 +/- 0.054 | 0 | 0.1 |
| Urban Bangalore | 32 | 0.078 +/- 0.034 | 0.063 +/- 0.030 | 0.109 +/- 0.039 | 0.346 | -1.191 |
| Agharia | 13 | 0.000 +/- 0.000 | 0.000 +/- 0.000 | 0.155 +/- 0.071 | NA | -2.186 |
| Saryupari Brahmin | 11 | 0.091 +/- 0.061 | 0.091 +/- 0.061 | 0.273 +/- 0.095 | 0 | -1.918 |

* indicates Z-score > 3

### Supplementary Note 5. Relationship between Ancestry and frequency of LP-associated variants

In this note, we examine how the allele frequency of -13.910*T is influenced by differences in ancestry across South Asia. Because -13.910:C>T confers a dominant phenotype, we also examine the frequency of lactase persistence (referred to as “LP frequency”) by weighting the heterozygous and homozygous genotypes equally (see Methods).

#### 5.1 Relationship between ancestry (using PCA loadings) and allele frequency of -13.910:C>T

We first explored the relationship between ancestry (as captured by PC loadings in Figure 1B) and the allele frequency of -13.910:C>T for 7,391 individuals (excluding individuals with >10% African-related ancestry component from ADMIXTURE. We also exclude endogamous groups that we project later). We binned individuals according to PC1 and PC2 values so the same number of individuals were in each bin (see Methods). Then we examined the linear relationship between median PC value per bin and the mean -13.910*T allele frequency. We find that PC1 is significantly correlated with -13.910*T frequency ($r^2$ = 0.91, p-value < 1e-06) as is PC2 ($r^2$ = 0.87, p-value < 1e-06) (Figure S5.1 B). We observe qualitatively similar results for the association between ancestry and LP frequency, accounting for the dominant mode of inheritance of -13.910:C>T (Figure S5.2)

**A.**

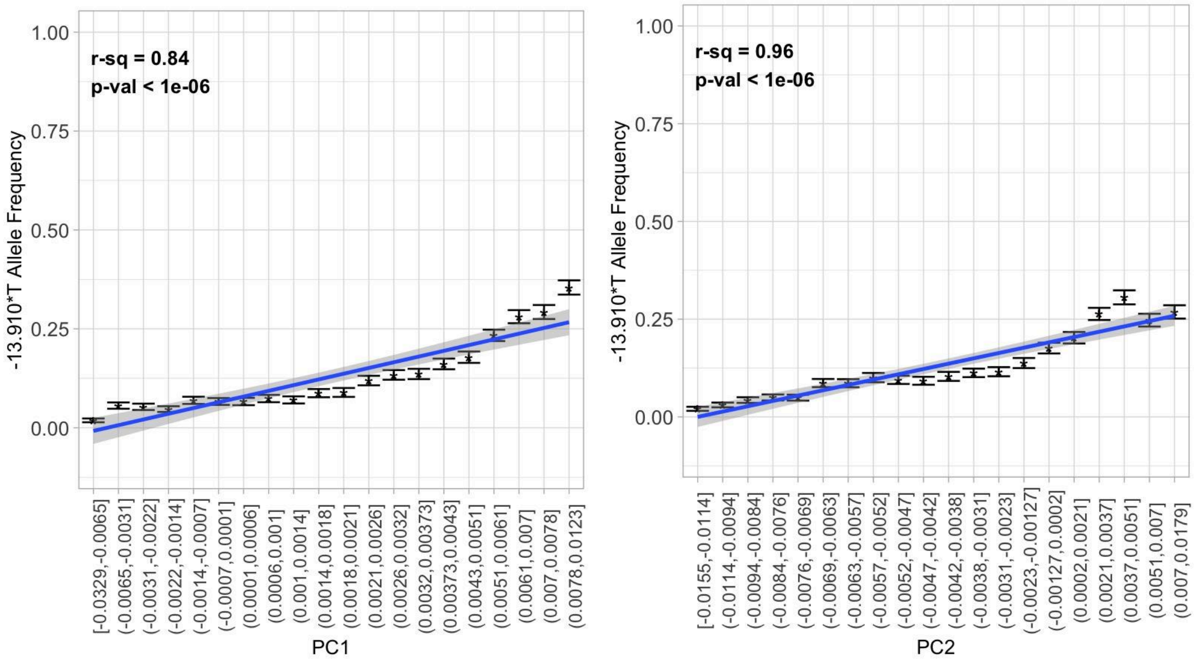

**B.**

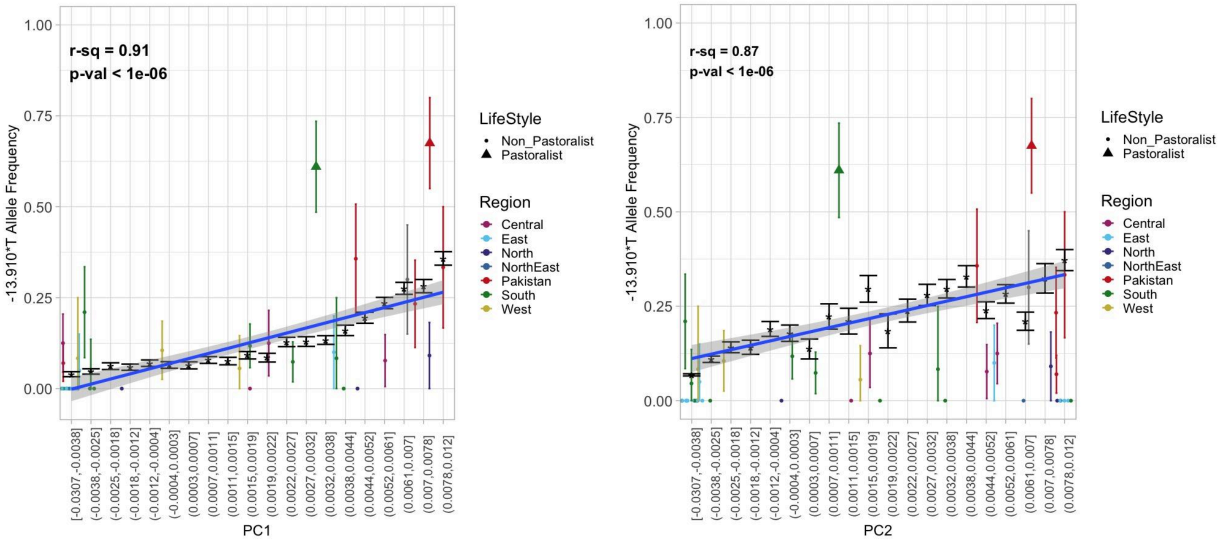


**Figure S5.1 The correlation between PC1 and PC2 and average -13.910:C>T allele frequency estimates for South Asians**

A. **All South Asian individuals** All South Asian individuals - India, Pakistan, and Bangladesh - were grouped into bins according to their PC1 and PC2 loadings. Each bin's average allele frequency was calculated and plotted along the y-axis. The regression line shows the relationship between each bin's median PC value and corresponding mean allele frequency.
B. **All South Asian individuals with endogamous groups projected** Individuals from endogamous groups across South Asia were excluded from initial binning and regression line calculations. The average allele frequency for each endogamous group was plotted against their respective bin on the X axis given their average PC value. The non-pastoralist endogamous groups are shown as dots, and pastoralist groups shown as triangles, with each group colored according to their geographic region. The error bars for each group reflect the 95% CI from bootstrapping.

**A.**

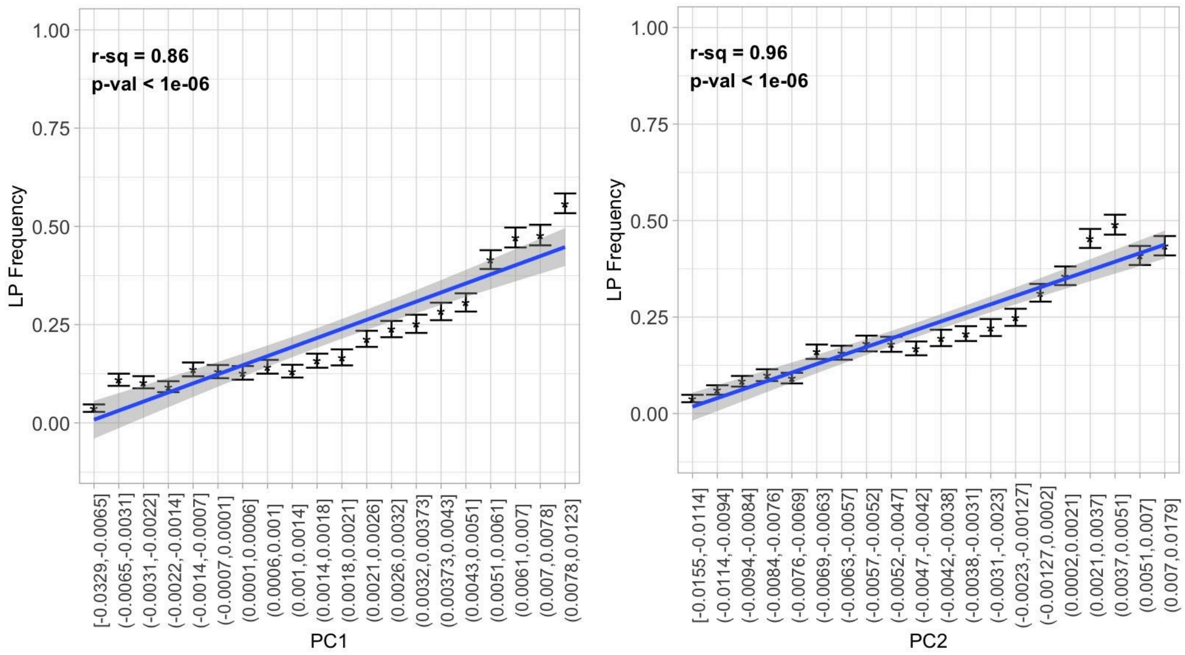


**B.**

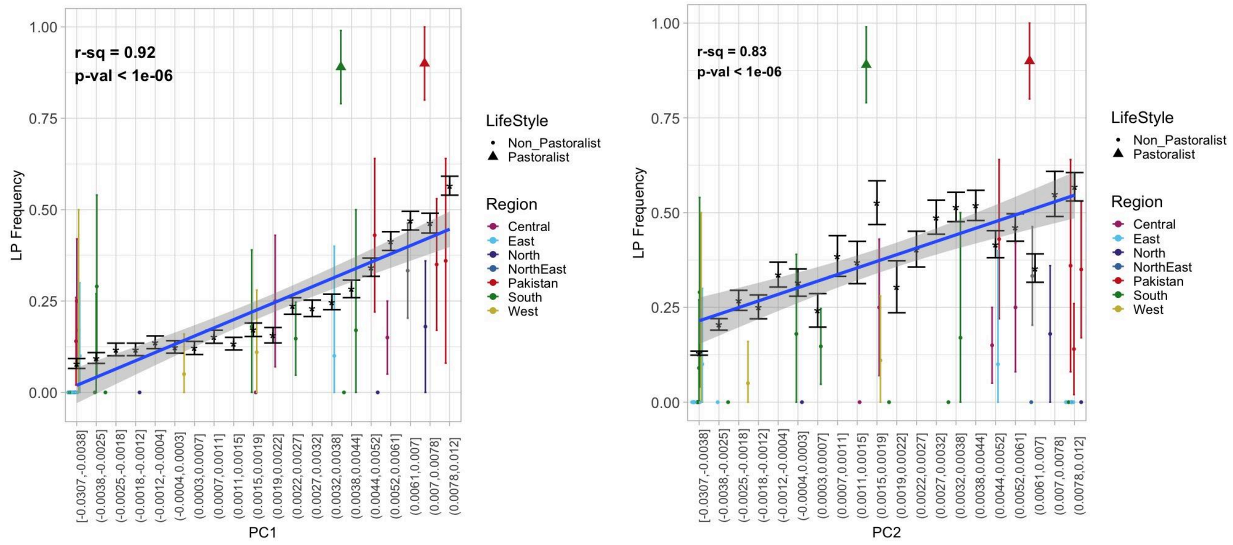


**Figure S5.2 The correlation between PC weights and LP estimates in South Asians.**

A. **All South Asian individuals** All South Asian individuals - India, Pakistan, and Bangladesh - were grouped into bins according to their PC1 and PC2 loadings. Each bin's average LP was calculated and plotted along the y-axis. The regression line shows the relationship between each bin's median PC value and corresponding mean LP.
B. **All South Asian individuals with endogamous groups projected** Individuals from endogamous groups across South Asia were excluded from initial binning and regression line calculations. The average LP for each endogamous group was plotted against their respective bin on the X axis given their average PC value. The non-pastoralist endogamous groups are shown as dots, and pastoralist groups shown as

triangles, with each group colored according to their geographic region. The error bars for each group reflect the 95% CI from bootstrapping.

#### 5.2 Relationship between genomewide ancestry (using qpAdm) and allele frequency of -13.910*T

To more directly study the association between ancestry and the frequency of -13.910*T in South Asia, we used the estimates of ancestry proportions based on qpAdm. We binned individuals according to their ancestry coefficient (for each ancestry group), ensuring an even number of individuals across bins. Within each bin, we calculated the mean allele frequency and examined the linear relationship with the median ancestry coefficient. We used the estimates based on:

a) Individuals on the Indian cline ($n$ = 4,946): We used the ancestry estimates based on the three-way model with SAHG-, Iranian farmer- and Steppe pastoralist-related ancestries.

b) Individuals outside the Indian cline ($n$ = 477): We used the ancestry estimates for the three-way and four-way model with SAHG-, Iranian farmer-, Steppe pastoralist- and East Asian-related ancestries.

For individuals that fit the three-way model, we observed a strong positive relationship between the proportion of Steppe pastoralist-related ancestry for a given individual and -13.910*T frequency ($r^2$ = 0.97, p-value < 1e-06) (Figure 3). Turning to Iranian farmer-related ancestry, we find weaker correlation with allele frequency, but still significant ($r^2$ = 0.69, p-value = 7.0e-04) (Figure 3). We observed an inverse relationship between SAHG-related ancestry and allele frequency ($r^2$ = -0.96, p-value < 1e-06) (Figure 3), consistent with our previous observations that Steppe pastoralist-related ancestry and SAHG-related ancestry are inversely related in South Asians (Figure 1C). We obtain qualitatively similar results when using LP frequency instead of allele frequency in the regression analysis (Figure S5.4 A).

For individuals outside the Indian cline, we used qpAdm results for individuals that fit with the three-way model or four-way model (see Methods). We find again the strongest correlation between -13.910*T allele frequency and Steppe pastoralist-related ancestry ($r^2$ = 0.97, p-value < 1e-06), a weaker, but still significant correlation with Iranian farmer-related ancestry ($r^2$ = 0.68, p-value = 9.3e-04). Moreover, we observe an inverse relationship with SAHG-related ancestry ($r^2$ = -0.96, p-value < 1e-06) and no significant association with East Asian-related ancestry ($r^2$ = -0.18 , p-value = 0.44) (Figure S5.3). When considering LP frequency instead of allele frequency, the results are qualitatively similar (Figure S5.4 B).

Since both Steppe pastoralist- and Iranian farmer-related ancestries appear to be significantly correlated with allele frequency, we evaluated their joint impact on -13.910*T allele frequency. We examined the correlation using the combined Steppe and Iranian farmer-related ancestries, rather than each separately, and found the correlation to be weaker than with Steppe ancestry alone ($r^2$ = 0.61, p-value < 1e-6). To assess the specific contribution of Iranian farmer-related ancestry, we performed a permutation test by shuffling its proportions and re-estimating the correlation. In all cases, we observe that the correlation in the combined ancestry model remained lower than with Steppe pastoralist-related ancestry alone (Figure S5.5). These results hold true when considering LP frequency instead of allele frequency (Figure S5.6). These results indicate that -13.910:C>T allele frequency is best explained by Steppe pastoralist-related ancestry, consistent with local ancestry analysis (Supplementary Note 4).

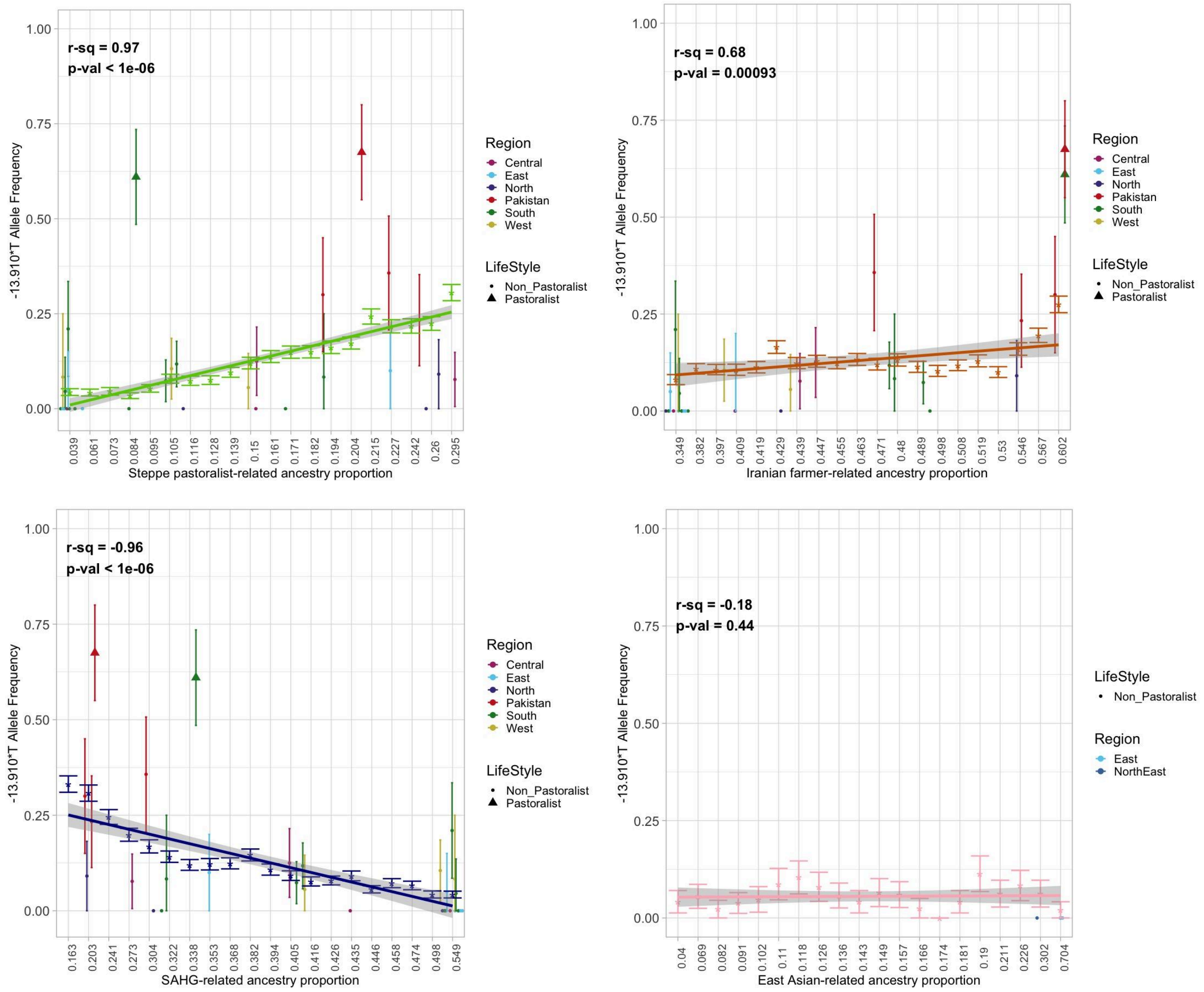


**Figure S5.3 The correlation between ancestry proportions and allele frequency of -13.910:C>T in South Asians** All South Asian individuals from North-East India and Bangladesh were grouped into bins according to their Steppe pastoralist-related, Iranian farmer-related, SAHG-related, or East Asian-related ancestry proportions estimated from qpAdm so an even number of individuals were in each bin. Each bin's average allele frequency was calculated and plotted along the y-axis. The regression line shows the relationship between each bin's median ancestry proportion and corresponding mean allele frequency. The relationship between allele frequency and Steppe pastoralist-related ancestry is shown in light green, Iranian farmer-related ancestry in dark orange, SAHG-related ancestry in navy blue, and East Asian-related ancestry in pink.

**A.**

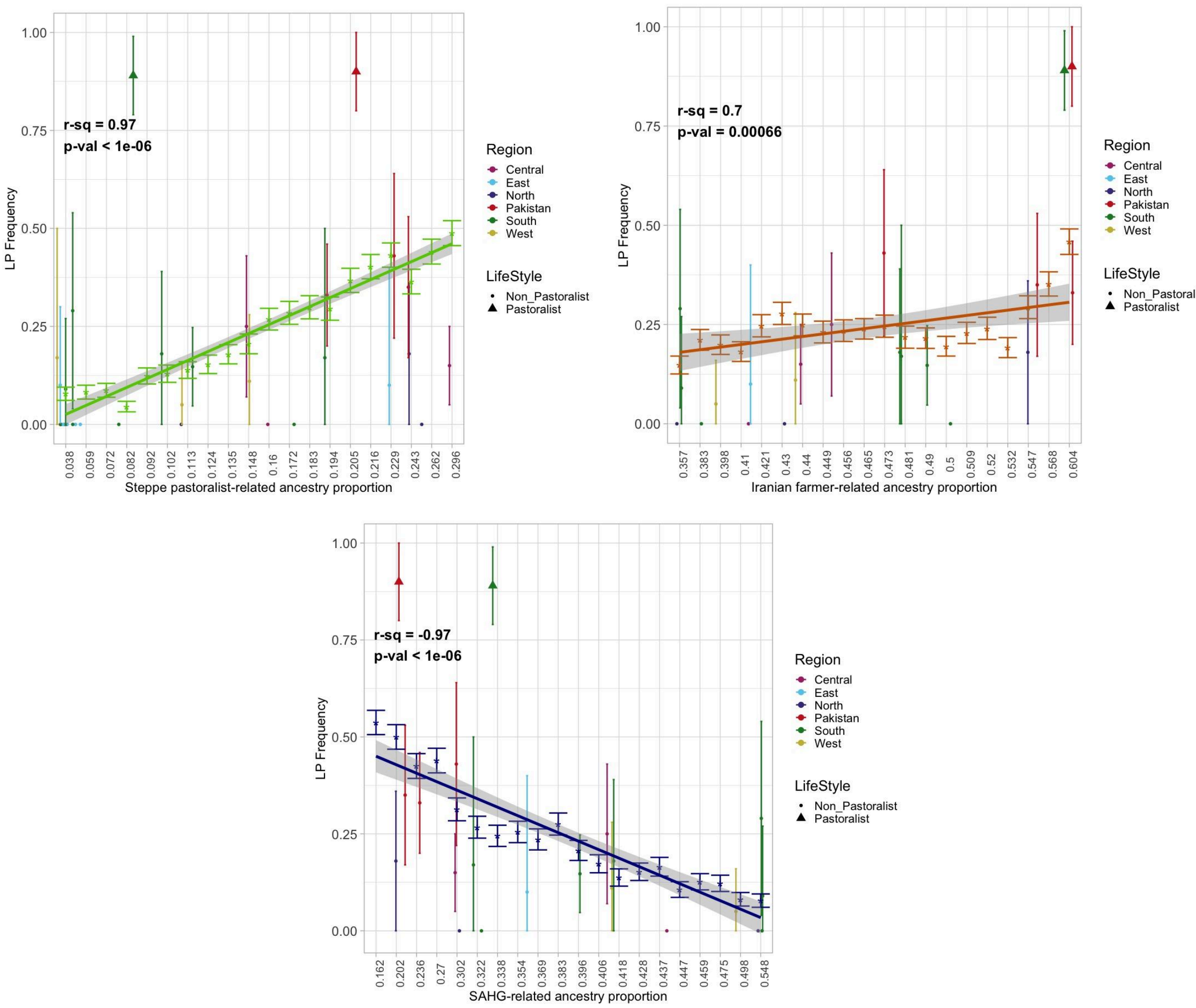
r-sq = 0.97
p-val < 1e-06
LP Frequency
Steppe pastoralist-related ancestry proportion
Region
Central
East
North
Pakistan
South
West
LifeStyle
Non_Pastoralist
Pastoralist
r-sq = 0.7
p-val = 0.00066
Iranian farmer-related ancestry proportion
r-sq = -0.97
p-val < 1e-06
SAHG-related ancestry proportion

**B.**

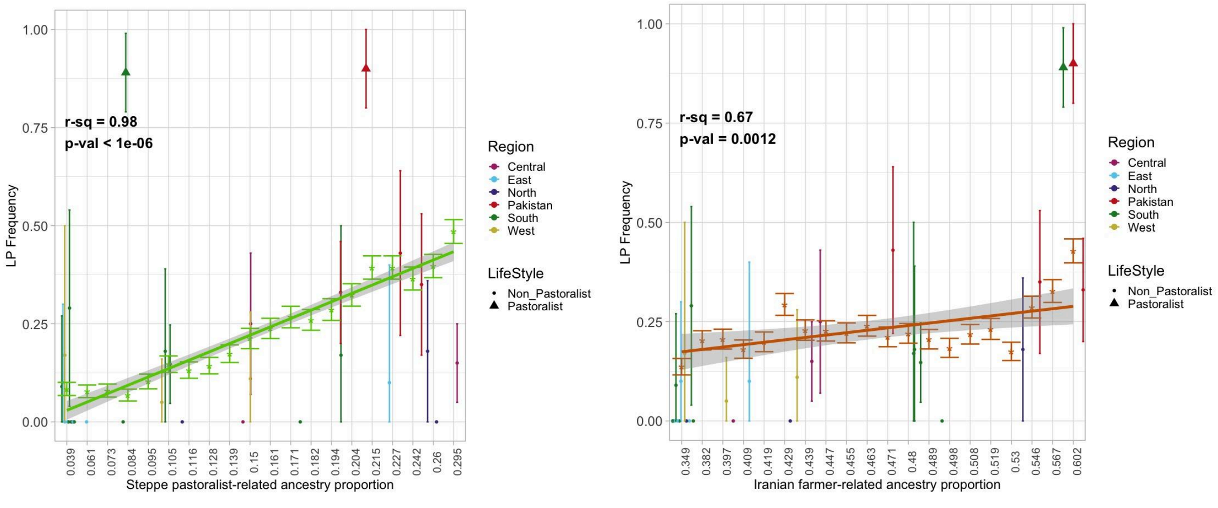
r-sq = 0.98
p-val < 1e-06
LP Frequency
Steppe pastoralist-related ancestry proportion
Region
Central
East
North
Pakistan
South
West
LifeStyle
Non_Pastoralist
Pastoralist
r-sq = 0.67
p-val = 0.0012
Iranian farmer-related ancestry proportion

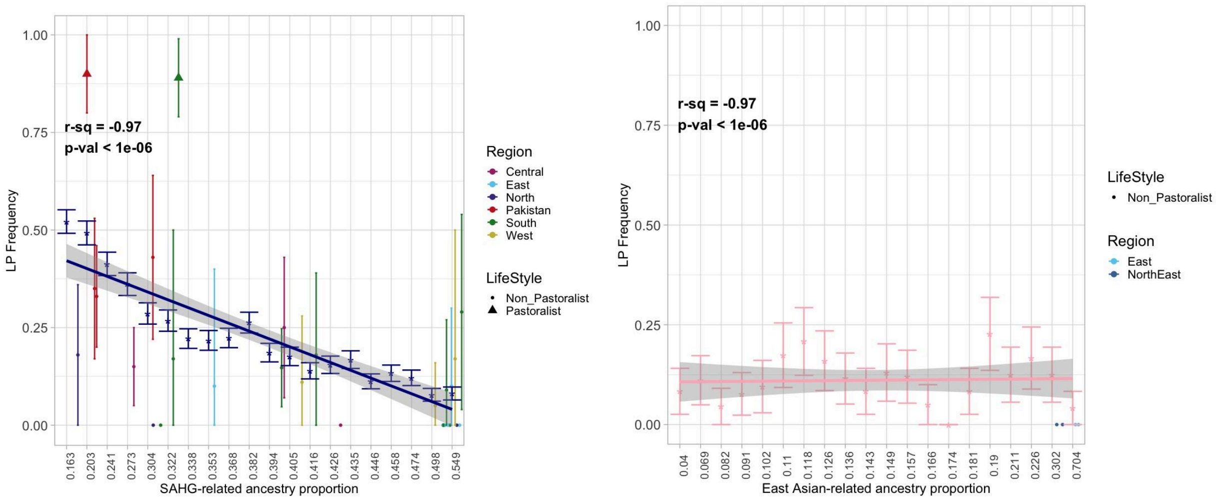


**Figure S5.4 The correlation between ancestry proportions and LP frequency of -13.910:C>T in South Asians**

A. **Ancestry Proportions based on the three-way model (Steppe pastoralist-, Iranian farmer-, and SAHG-related ancestries).** All South Asian individuals - India, and Pakistan- were grouped into bins according to their Steppe pastoralist-related, Iranian farmer-related, or SAHG-related ancestry proportions estimated from qpAdm so an even number of individuals were in each bin. Each bin's average LP was calculated and plotted along the y-axis. The regression line shows the relationship between each bin's median ancestry proportion and corresponding mean LP. The estimates from endogamous groups were projected on the plot, with pastoralist groups shown in triangles and non-pastoralists shown in dots, with each group colored by geographic region. The relationship between LP and Steppe pastoralist-related ancestry is shown in light green, Iranian farmer-related ancestry in dark orange, and SAHG-related ancestry in navy blue.
B. **Ancestry Proportions based on four-way model (Steppe pastoralist-, Iranian farmer-, SAHG-, and East Asian-related ancestries).** All South Asian individuals from North-East India and Bangladesh were grouped into bins according to their Steppe pastoralist-related, Iranian farmer-related, SAHG-related, or East Asian-related ancestry proportions estimated from qpAdm so an even number of individuals were in each bin. Each bin's average LP was calculated and plotted along the y-axis. The regression line shows the relationship between each bin's median ancestry proportion and corresponding mean LP. The relationship between LP and Steppe pastoralist-related ancestry is shown in light green, Iranian farmer-related ancestry in dark orange, SAHG-related ancestry in navy blue, and East Asian-related ancestry in pink.

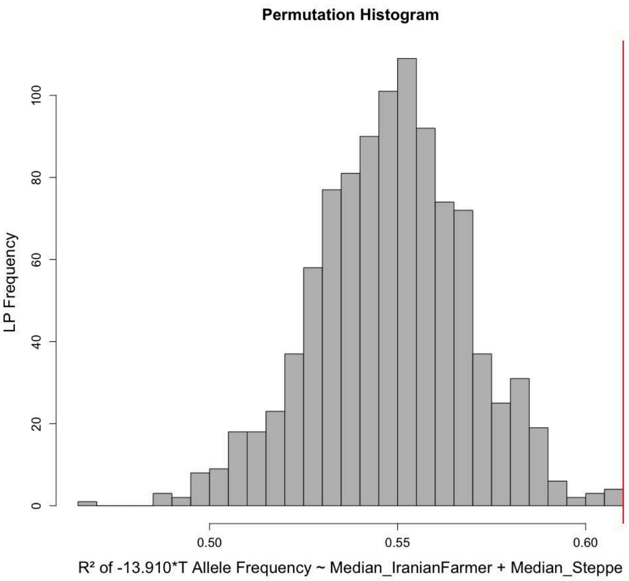


**Figure S5.5 Permutation test to evaluate the regression model including both Steppe pastoralist and Iranian farmer ancestries vs. the Steppe pastoralist ancestry only** We tested whether Iranian farmer combined with Steppe pastoralist ancestry was a better predictor of allele frequency than Steppe ancestry alone. We calculated the correlation coefficient between allele frequency and ancestry for both of these models. We permuted allele frequencies across populations (n_perm = 1000). The histogram shows the distribution of the correlation coefficients of the null hypothesis of no association. The red line represents $r^2 = 0.61$, the real correlation coefficient of the combined ancestry model. In all permutations, the correlation of the combined model was weaker than with Steppe pastoralist ancestry alone.

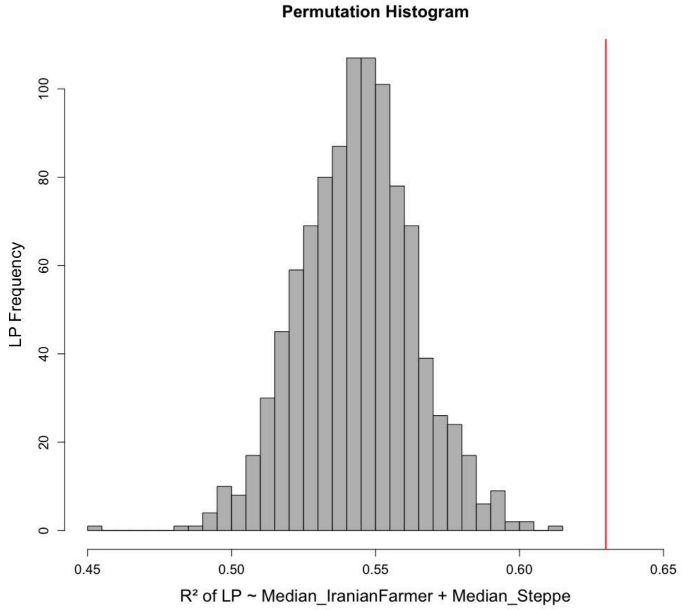


**Figure S5.6 Permutation test to evaluate the regression model including both Steppe pastoralist and Iranian farmer ancestries vs. the Steppe pastoralist ancestry only.** We tested whether Iranian farmer combined with Steppe pastoralist ancestry was a better predictor of LP frequency than Steppe ancestry alone. We calculated the correlation coefficient between LP frequency and ancestry for both of these models. We permuted LP frequencies across populations (n_perm = 1000). The histogram shows the distribution of the correlation coefficients of the null hypothesis of no association. The red line represents $r^2 = 0.63$, the real correlation coefficient of the combined ancestry model. In all permutations, the correlation of the combined model was weaker than with Steppe pastoralist ancestry alone.

### 5.3 Relationship between genome-wide ancestry and local ancestry at -13.910:C>T locus

We next examined the relationship between genome-wide ancestry proportions and local ancestry in the core region. As done earlier, individuals were binned according to their genome-wide ancestry proportions (for each geographic or endogamous), ensuring equal sample sizes across bins. Within each bin, we calculated the mean local ancestry proportion at the core haplotype (local Steppe*T) and evaluated its linear relationship with the median genome-wide ancestry coefficient.

For individuals on the Indian cline, we observed a strong positive correlation between Steppe pastoralist-related genome-wide and local Steppe*T ancestries ($r^2 = 0.97$, p-value < 7.24e-13) (Figure S5.7A). The correlation between Iranian farmer-related genome-wide ancestry and local Steppe*T was weaker but still significant ($r^2 = 0.69$, p-value = 0.000746) (Figure S5.7B). In contrast, SAHG-related genome-wide ancestry showed a negative correlation with local Steppe*T ancestry ($r^2 = -0.96$, p-value < 2.61e-11) (Figure S5.7C). These patterns are consistent with similar correlation seen between genome-wide Steppe pastoralist ancestry and allele frequency.

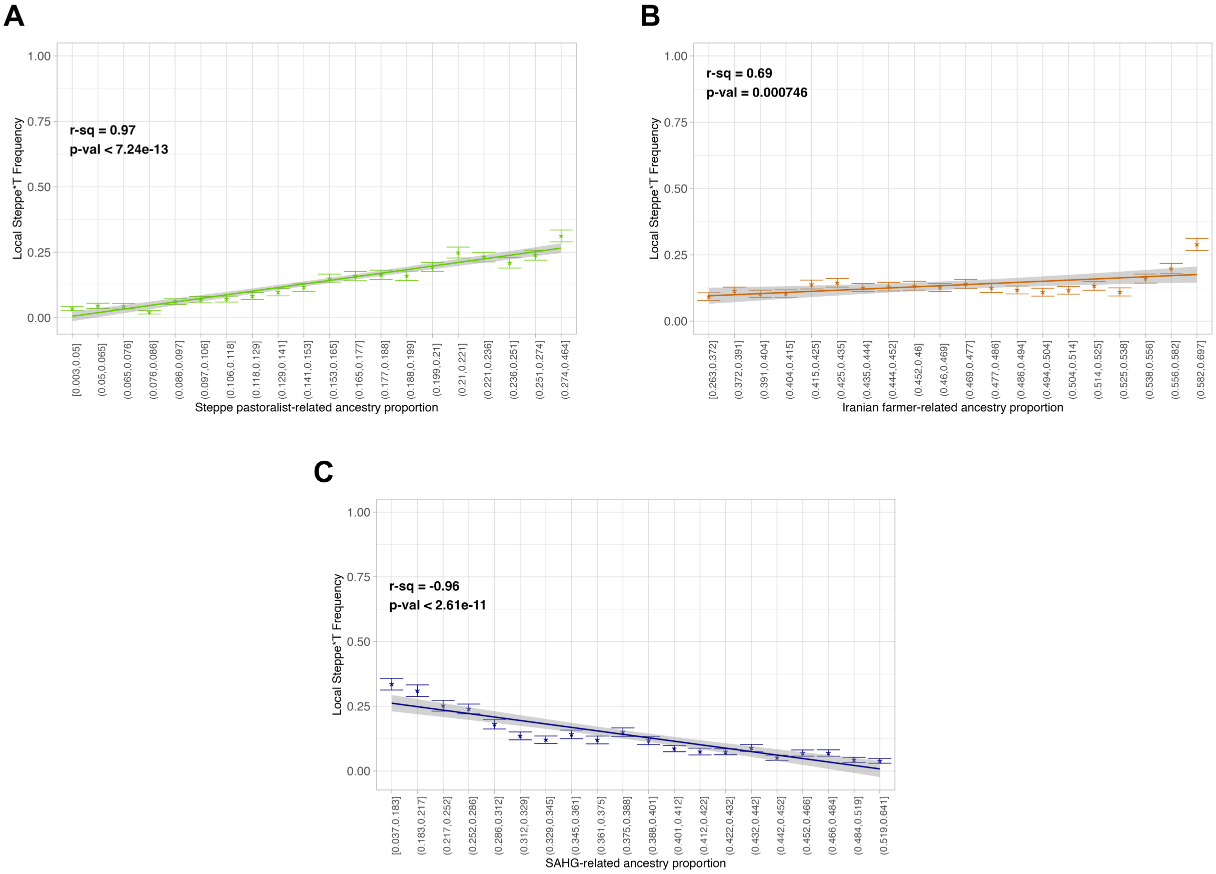

**Figure S5.7 The correlation between genome-wide ancestry proportions and local Steppe*T frequency in South Asians.** South Asian individuals were grouped into bins according to their Steppe pastoralist-related (green), Iranian farmer-related (orange), or SAHG-related (blue) ancestry proportions estimated from qpAdm so an even number of individuals were in each bin. Each bin's average local Steppe*T frequency was calculated and plotted along the y-axis. The regression line shows the relationship between each bin's median genome-wide ancestry proportion and corresponding local Steppe*T frequency.

### Supplementary Note 6. Simulations to Examine the Role of Demographic History in Shaping LP Prevalence in Toda and Gujjar

To understand how the demographic history of Toda and Gujjar has shaped lactase persistence, we performed simulations to examine if drift alone can explain the observed high frequency of -13.910:C>T variant in these groups. Specifically, we modelled key aspects of the history of each population and estimated the likelihood of observing the current -13.910*T allele frequency in the absence of selection.

#### 6.1 Steppe Pastoralist-Related Gene Flow

Both Toda and Gujjar have ancestry from three ancestral sources related to South Asian hunter-gatherers (SAHG), ancient Iranian farmers and ancient Eurasian Steppe pastoralists (see Supplementary Note 1). Using qpAdm, we estimated the ancestry composition of Toda as 7.8% +/- 2.1% Steppe pastoralist-related, 58.9 +/- 2.8% Iranian farmer-related, and 33.2 +/- 2.0% SAHG-related ancestries. For the Gujjar, the estimates were 21.1 +/- 3.1% Steppe pastoralist-related, 59.1 +/- 5.5% Iranian farmer-related, and 19.8 +/- 6.0% SAHG-related ancestries (Supplementary Table 1.2). Investigation of the haplotype structure around *LCT* locus suggests that all individuals carrying -13.910*T allele harbor Steppe pastoralist-related ancestry at this locus (see Supplement Note 4). Previous studies have shown that the Steppe pastoralist-related gene flow occurred around 3,500 years ago in South Asia[4].

#### 6.2 History of Founder Events

Many South Asian groups have experienced strong founder events in their recent past[8–11]). To measure the strength of founder events in South Asia, we applied ASCEND[8] that uses the correlation in allele sharing patterns across the genome to infer the timing and the intensity of founder events. The founder intensity is a composite parameter that is a function of the duration of the bottleneck and the population size during the bottleneck[8]:

$$\text{Founder intensity } (I_f) = [\text{Duration of bottleneck } (D_f)]/[2*\text{Population size } (N_{bottleneck})]$$

We applied ASCEND to Toda and Gujjar using data for a set of 1 million randomly selected SNPs. We used 15 random individuals from the GAsP1 dataset as outgroups. This analysis revealed significant evidence of founder events in both groups. For Toda, the inferred intensity and timing is 8.5 +/- 0.2% and 19.1 +/- 0.8 generations (532 ± 28 years ago, assuming human generation time of 28 years[32]). For Gujjar, we infer an intensity and timing of 0.8 +/- 0.2% and 115.1 +/- 40.5 generations (3223 ± 1134 years). Assuming a duration of 100 generations, the founder intensity translates to an estimated effective population size during the bottleneck of 588.2 for Toda and 6250 for Gujjar.

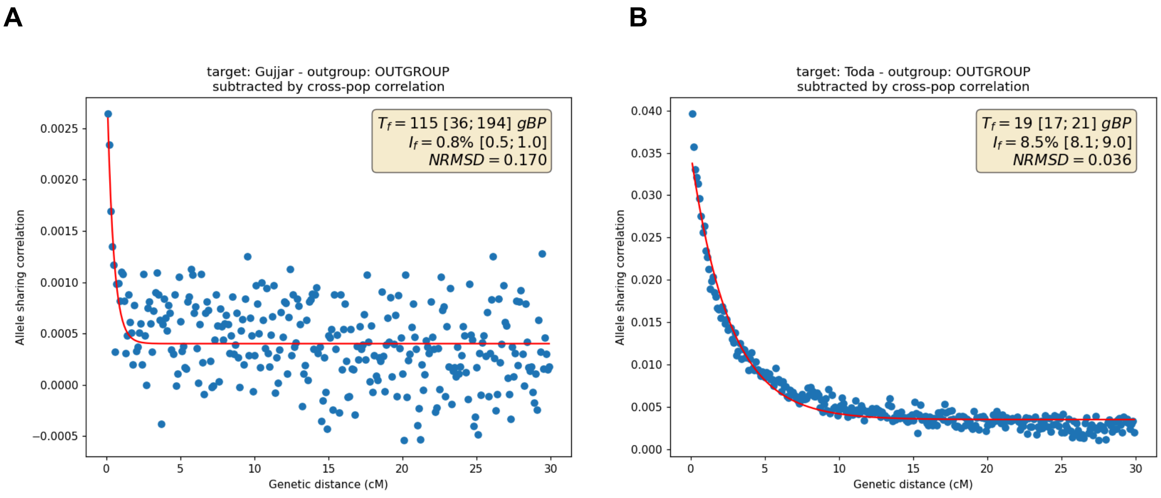


**Figure S6.1: Fit of ASCEND for Gujjar and Toda.** We show the model fits inferred using ASCEND for (A) Gujjar and (B) Toda, plotting allele sharing metric against the genetic distance. The inferred value for the date of the bottleneck ($T_f$), the founder intensity ($I_f$), their 95 %CI, and the Normalized Root Mean Square Deviation (NRMSD) are indicated for each plot.

#### 6.3 Simulation Scenarios

We conducted coalescent simulations using msprime[33] to model demographic scenarios based on the inferred history of Toda and Gujjar populations, incorporating gene flow from Steppe pastoralist-related groups followed by founder events (Figure S6.2). Specifically, we assume there are two populations— Pop1 and Pop2 which broadly resemble ancestral South Asians (SAHG) and ancestral Steppe pastoralists. These groups diverged ~30,000 years ago[34]. Around 125 generations ago, there was gene flow from Steppe pastoralist-related population into SAHG forming the contemporary population (South Asia) sampled in our analysis (equivalent to 3,500 years ago, assuming 28 years per generation[4]). The initial population size of both ancestral groups was 10,000. In the past 120 generations, both populations had founder events. To account for the uncertainty in the strength of the bottleneck, we explored a range of effective population sizes during the bottleneck: $N_{bottleneck}$ = [5000, 1000, 500, 100, 90, 70, 50, 30, 10], as well as a constant population scenario. For simplicity, we do not include Iranian farmer-related gene flow so that we can track the haplotypes inherited from Steppe pastoralist population without confounding.

We consider two demographic scenarios: ‘Constant’ and ‘Bottleneck’ (Figure S6.2). In the ‘Constant’ model, the effective population size is maintained at 10,000 throughout time. In the ‘Bottleneck’ scenario, the South Asian population experiences a bottleneck beginning 120 generations before present ($t_{start}$ = 120) and lasting for 100 generations until 20 generations before present ($t_{end}$ = 20) (Figure S6.2). During this period, we varied the effective population size ($N_{bottleneck}$ = 10–5000), after which the population size returned to $N_{South\ Asia}$ (= 10,000). For each simulation, we generated a genome sequence corresponding to human chromosome 14 (106,880,170 bp in length) using a recombination rate derived from the HapMap recombination map for chromosome 14. We sampled 500 chromosomes from the 'South Asia' population and used tree sequences from msprime to identify the exact locations, lengths, and frequencies of Steppe pastoralist-related ancestry regions in each individual. To ensure robustness, we conducted 100 independent simulations and performed bootstrap resampling (with replacement) to estimate the mean and standard error of each summary statistic.

The *msprime* command for the '*Constant*' demographic model:

```
demography_simple = msprime.Demography() #initializing an msprime "demography" object
# This is the "trunk" population that we merge other populations into
demography_simple.add_population(
    name="SAHG",
    description="Ancestral South Asians",
    initial_size=N_SAHG,
    initially_active=True,
);

demography_simple.add_population(
    name="SouthAsia",
    description="SouthAsian",
    initial_size=N_SAsia,
    initially_active=True,
)
demography_simple.add_population(
    name="Steppe",
    description="Steppe",
    initial_size=N_Steppe,
    initially_active=True,
)

# Split Steppe and South Asians
demography_simple.add_population_split(
    time=T_europa_asia, derived=["SouthAsia"], ancestral="SAHG"
)

demography_simple.add_population_split(
    time=T_europa_asia, derived=["Steppe"], ancestral="SAHG"
)

# Add Steppe pastoralist-related geneflow for one generation (pulse model)
demography_simple.add_migration_rate_change(time = T_Steppe_into_SAS, rate = admixtureproportion,
dest = "Steppe", source = "SouthAsia")
demography_simple.add_migration_rate_change(time = T_Steppe_into_SAS+1, rate = 0, dest =
"Steppe", source = "SouthAsia")

demography_simple.sort_events()
```

The *msprime* command added for the '*Bottleneck*' demographic model:

```
# Add Bottleneck in Asian population
demography_simple.add_population_parameters_change(time=tstart, population="SouthAsia",
initial_size=N_SAsia)

demography_simple.add_population_parameters_change(time=tend, population="SouthAsia",
initial_size=N_bottleneck)
```

#### Model Parameters

### Times provided in years, then converted in generations.
gen_time = 28.0 #generation time
### Population sizes
N_SAsia = 10_000
N_Steppe=10_000
N_SAHG=10_000
### Split Times - converting years BP into generations BP
T_europa_asia = 30_000 / gen_time
### Steppe pastoralist-related ancestry admixture times
T_Steppe_into_SAS = 3_500 / gen_time

Bottleneck parameters:
$N_{bottleneck}$: 5000, 1000, 500, 100, 90, 70, 50, 30, 10
tstart: 120
tend: 20
*Admixture parameters:*
*$a_S$: 7.8%, 21%*

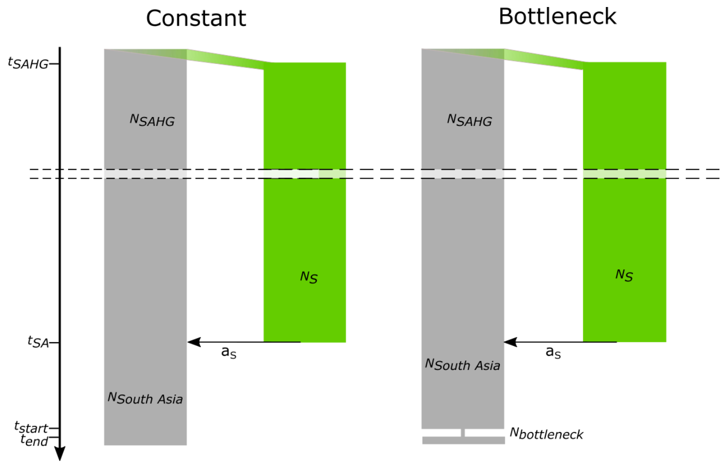


**Figure S6.2. Two scenarios simulated using msprime.** With effective population size of the ancestral South Asians ($N_{SAHG}$), effective population size of South Asia ($N_{South\ Asia}$), effective population size of ancestral Steppe pastoralist-related population ($N_S$) and effective population size during the bottleneck ($N_{bottleneck}$). We define $t_{SAHG}$ as the split time between Steppe pastoralist-related and other Eurasians lineages and $t_{SA}$ as the time of Steppe pastoralist-related admixture and $a_S$ as the admixture proportion.

#### 6.4 Distribution of Steppe Pastoralist-Related Ancestry in Simulations

We simulated data under two demographic scenarios: ‘Constant’ and ‘Bottleneck’ with a range of population sizes ($N_{bottleneck}$ = 10–5000) during the bottleneck. We examined the change in Steppe pastoralist-related ancestry proportion as a proxy for -13.910*T frequency. We assume the initial Steppe-pastoralist related gene flow ($p_{initial}$) was similar to the observed genome-wide estimate in present-day Toda (7.8%) and Gujjar (21%) and the gene flow occurred 3,500 years ago[4] (Fig S6.1). We analyzed the frequency of Steppe pastoralist-related haplotypes across simulated chromosomes, including all Steppe haplotypes regardless of length or frequency in the ancestral population. For each simulation, we quantified the probability of Steppe pastoralist haplotypes reaching or exceeding the observed -13.910*T frequency ($p_{obs}$ = 65%).

Under the constant population size model, the probability to exceed $p_{obs}$ is 0 across 100 simulations, if $p_{initial}$ = 7.8% (Toda) or 21% (Gujjar). Even under an extreme bottleneck ($N_{bottleneck}$ = 10), the probability to exceed $p_{obs}$ is less than 0.1, ranging between 0.024 for $p_{initial}$ = 7.8% (Toda) to 0.089 for $p_{initial}$ = 21% (Gujjar). As the population size during the bottleneck ($N_{bottleneck}$) increases, $p_{obs}$ decreases from 1% ($N_{bottleneck}$ = 30–70, Figure S6.3) to less than $10^{-5}$ ($N_{bottleneck}$ = 100–1000, Figure S6.2). For comparison, the inferred $N_{bottleneck}$ for Toda ($N_{bottleneck}$ = 588.2) and Gujjar ($N_{bottleneck}$ = 6250) assuming a bottleneck duration of 100 generations, the probability to exceed $p_{obs}$ is 0. These results suggest that, given the inferred demographic histories of the Toda and Gujjar populations, genetic drift alone is insufficient to account for the observed high frequencies of the -13.910*T allele.

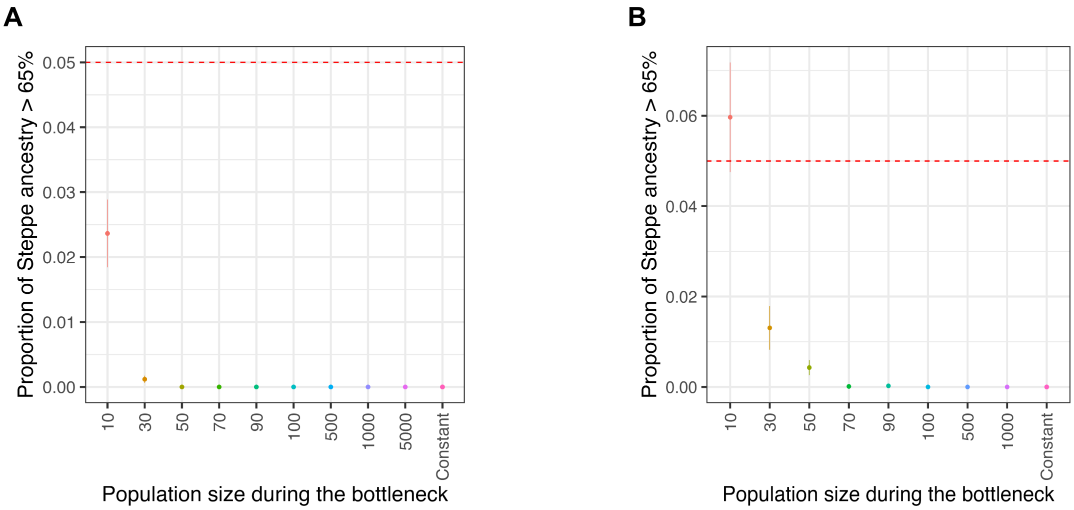


**Figure S6.3: Proportion of the regions inherited from Steppe pastoralist-related gene flow exceeding 65%, under different strengths of bottleneck (x-axis).** (A) For a Steppe pastoralist-related gene flow of 7.8% and (B) a Steppe pastoralist-related gene flow of 21%. Population size before and after the bottleneck N=10,000. The dashed red line corresponds to 5%. Error bars are generated from bootstrap resampling.

### Supplementary Note 7. Detecting Signatures of Natural Selection Using Identity-By-Descent Segments

In this section, we examine Identify-by-Descent (IBD) sharing at the *LCT* locus among endogamous groups from the GAsP1 dataset. We use the insight that a beneficial allele– and surrounding variants (haplotype)—rapidly increase in frequency in the population. Because this occurs faster than recombination can break down the allelic correlation in the region, individuals with the selected allele share long, nearly identical haplotypes, that are IBD, at the selected locus. By comparing the rate of IBD sharing across the genome, we can thus identify selected regions as they contain elevated rates of local IBD sharing, setting them apart from the average background relatedness.

For this analysis, we used phased data from South Asian endogamous groups and GBR that are part of the GAsP1 dataset (Supplementary Note 1). We identified IBD segments within each population separately using *hap-IBD*[35] and used the deCODE recombination map[36]. We then examined IBD sharing between populations which present an excess of IBD segments at the *LCT* locus: GBR/Gujjar, GBR/Toda and Gujjar/Toda. We describe IBD sharing patterns at the *LCT* locus and across chromosome 2, for endogamous groups with more than 10 representatives.

#### 7.1 IBD Sharing within a Population

Across South Asian endogamous groups and GBR, we examined patterns of IBD sharing at the *LCT* locus and compared to the average along chromosome 2 (Table S7.1). Most populations in South Asia exhibit low or background levels of IBD sharing at the *LCT* locus ($|Z| < 3$, Figure S7.1A), consistent with neutral expectations. However, two pastoralist groups—Gujjar and Toda—show markedly elevated rates around the -13.910:C>T variant similar to GBR (Figure 4B). Specifically, we identified 81, 62, and 111 shared IBD segments in GBR, Gujjar, and Toda individuals, corresponding to IBD rates of 0.071, 0.079, and 0.176, respectively. In all three populations, the rate of IBD sharing at the *LCT* locus significantly exceeds the chromosome 2 average rate of 0.001 (GBR), 0.003 (Gujjar) and 0.049 (Toda) respectively ($Z = 11.3$ for GBR, $Z = 12.9$ for Gujjar and $Z = 4.5$ for Toda, Supplementary Table S3). The lower Z-score in Toda reflects the presence of large number of IBD segments throughout chromosome 2, likely due to a history stronger founder events in this population (on average, Toda individuals have an IBD rate of 0.049, compared to 0.001 in GBR and 0.003 in Gujjar) (Figure S6.1B, see Supplementary Note S6). Together, these results indicate elevated IBD-sharing at the LCT locus in Toda and Gujjar—a pattern that is absent in non-pastoralist South Asians—consistent with strong selection in pastoralist groups.

**A**

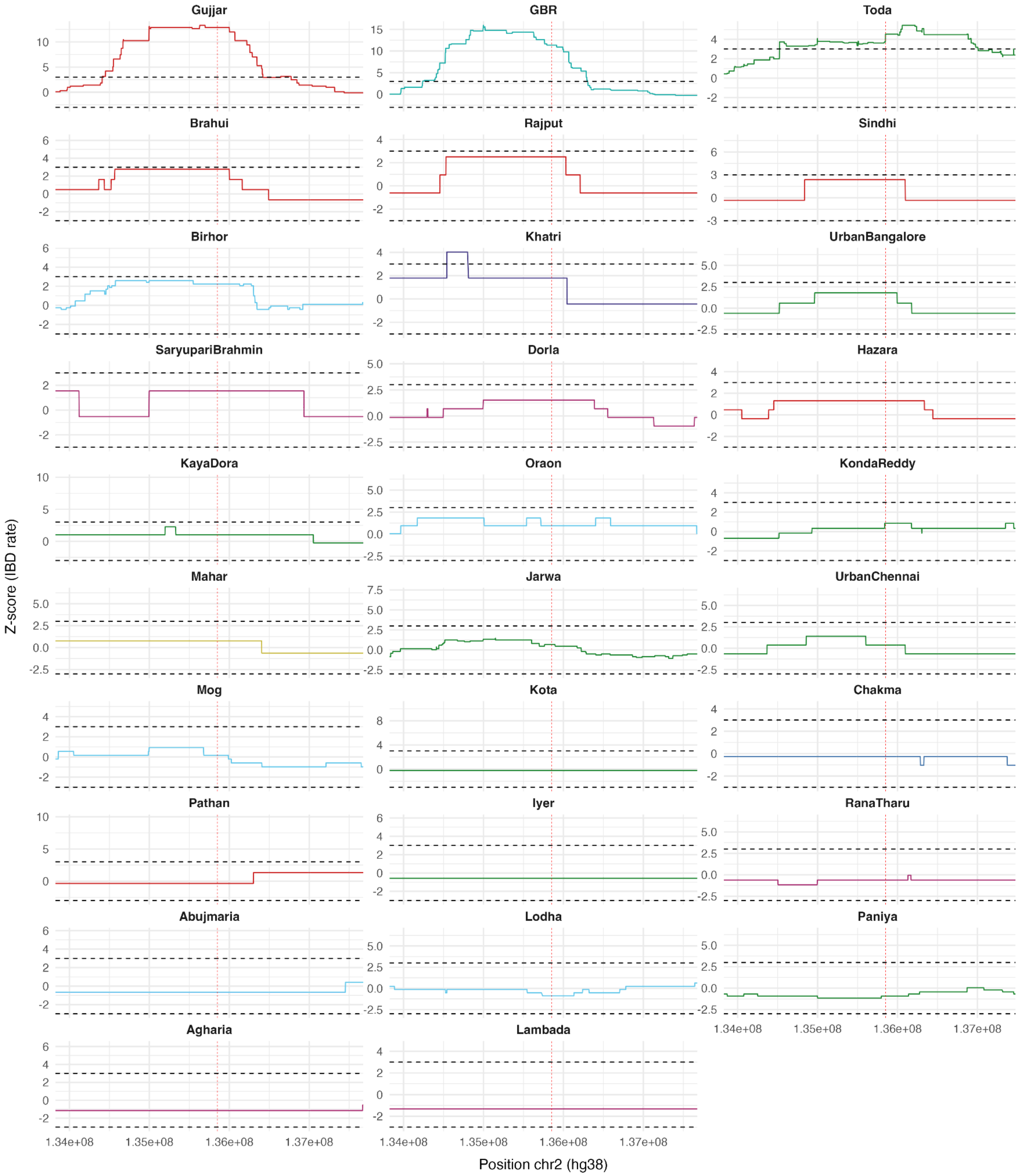

Gujjar
GBR
Toda
Brahui
Rajput
Sindhi
Birhor
Khatri
UrbanBangalore
SaryupariBrahmin
Dorla
Hazara
KayaDora
Oraon
KondaReddy
Mahar
Jarwa
UrbanChennai
Mog
Kota
Chakma
Pathan
Iyer
RanaTharu
Abujmaria
Lodha
Paniya
Agharia
Lambada
Z-score (IBD rate)
Position chr2 (hg38)
1.34e+08
1.35e+08
1.36e+08
1.37e+08

**B**

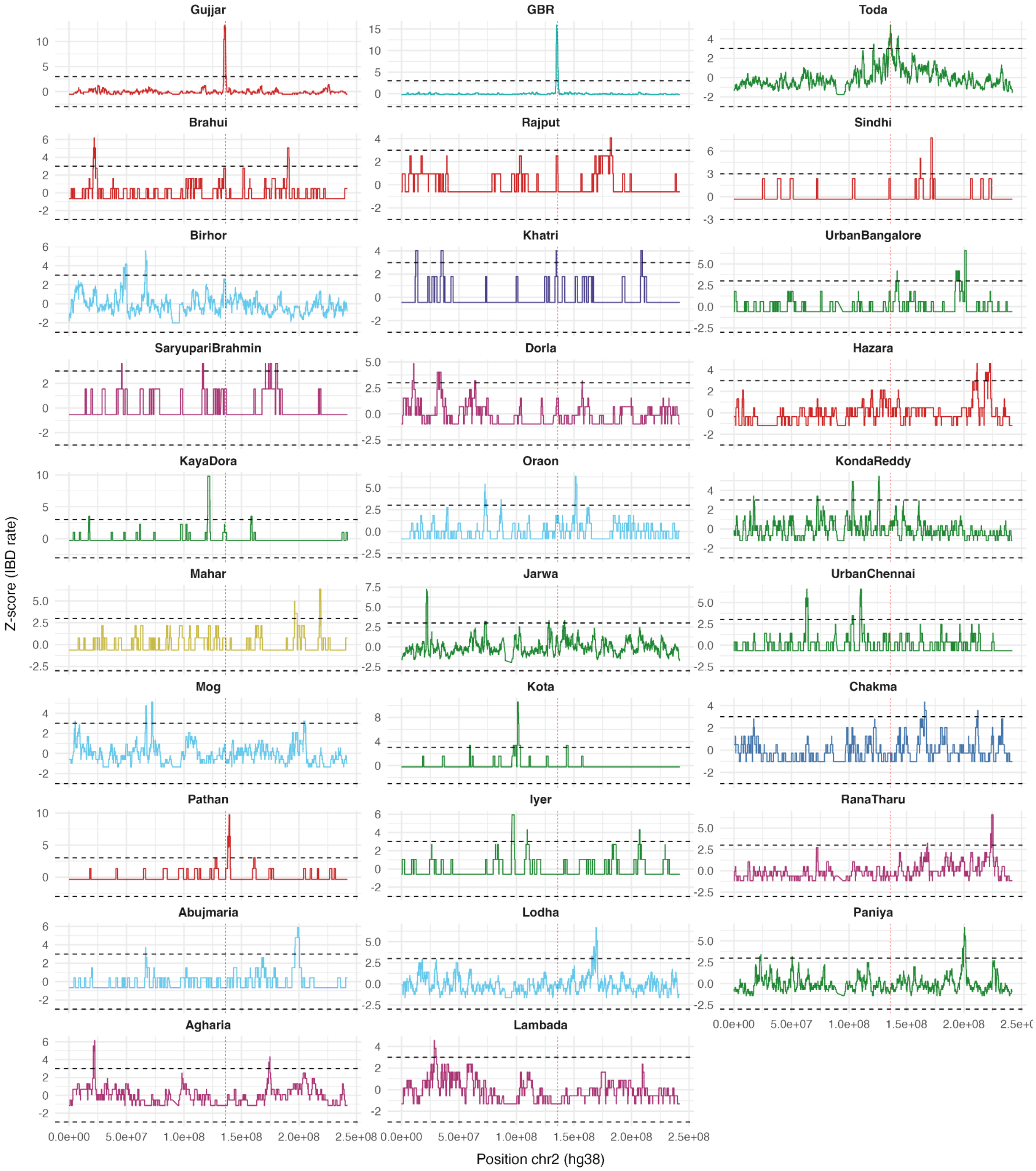


**Figure S7.1. Within population Z-score based on IBD rate on comparing local enrichment at *LCT* locus vs. genome-wide expectation.** We estimated IBD rates per endogamous group with more than 10 representatives in the dataset on chromosome 2 for (A) *LCT* locus from 134,000,00 to 137,500,000 bp (in hg38) and (B) genome-wide. The dashed horizontal lines represent the significance level 3 and -3.

#### 7.2 IBD Sharing between Populations

To investigate the shared origin of the shared haplotypes, we measured pairwise IBD sharing between individuals from Toda, Gujjar and GBR. Similarly to the within population analysis, we compute the number of IBD segments overlapping the -13.910:C>T variant and compare it to the chromosome-wide average statistics. We identified 74 shared IBD segments in GBR/Gujjar, 13 in GBR/Toda, and 42 in Gujjar/Toda individuals, translating to IBD rates of 0.038, 0.008 and 0.029, respectively (Table S7.1). For all pairs of populations, these estimates exceed the chromosome 2 average by more than ten standard deviations (Z = 13.9 for GBR/Gujjar, Z = 12.58 for GBR/Toda and Z = 9.4for Gujjar/Toda, Table S7.1). This high level of sharing of IBD segments across populations suggests shared origin across the two Eurasian subcontinents.

**Table S7.1. IBD over -13.910:C>T variant between South Asian pastoralist groups and the GBR population.**

| **Populations (number of individuals)** | **IBD rate of segments overlapping -13.910:C>T variant (number of IBD segment)** | **Average IBD rate (mean +/-standard deviation)** | **Average number of IBD segments on chr2 (mean +/- standard deviation)** |
|---|---|---|---|
| GBR/Gujjar( n=20,24) | 0.038 (74) | 0.0003 +/- 0.0028 | 0.57 +/- 5.29 |
| GBR/Toda (n=18,24) | 0.008 (13) | 6e-05 +/- 0.0006 | 0.100 +/- 1.03 |
| Gujjar/Toda (n=18,20) | 0.029 (42) | 0.0005 +/- 0.0030 | 0.73 +/- 4.4 |

*after filtering out IBD segments overlapping the -13.910:C>T variant, and segments shorter than 1Mb.

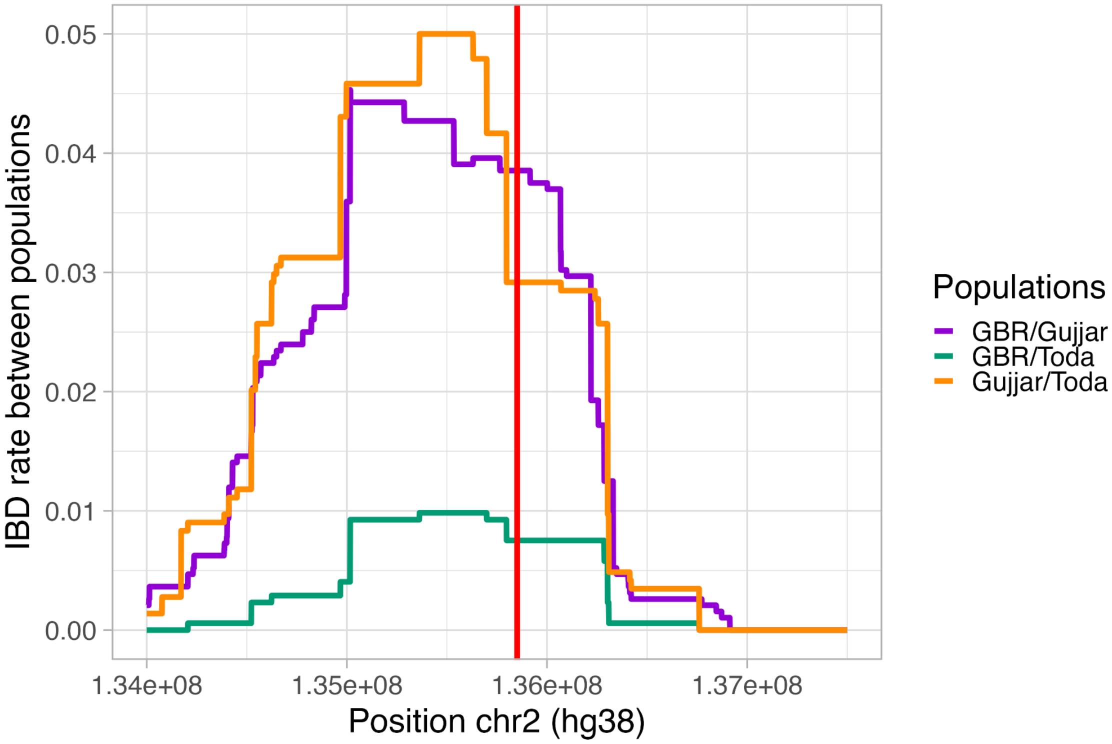


**Figure S7.2. Between populations IBD distribution on the -13.910:C>T variant.** IBD rate between individuals of three different populations: GBR/Gujjar in purple, GBR/Toda in orange and Gujjar/Toda in green. On chromosome 2, from 134,000,00 to 137,500,000 bp (in hg38), in red the position of the -13.910:C>T variant.
